# A repurposed drug screen identifies compounds that inhibit the binding of the COVID-19 spike protein to ACE2

**DOI:** 10.1101/2021.04.08.439071

**Authors:** Kaleb B. Tsegay, Christiana M. Adeyemi, Edward P. Gniffke, D. Noah Sather, John K. Walker, Stephen E. P. Smith

**Affiliations:** Center for Integrative Brain Research, Seattle Children’s Research Institute, Seattle, WA 98101; St. Louis University School of Medicine, Department of Pharmacology and Physiology, St. Louis, MO 63110; Center for Global Infectious Disease Research, Seattle Children’s Research Institute, Seattle, WA, 98101; University of Washington, Department of Pediatrics, Seattle, WA 98195; Henry and Amelia Nasrallah Center for Neuroscience, Saint Louis University St. Louis, MO 63110; Graduate Program in Neuroscience, University of Washington, Seattle, WA 98195

## Abstract

Repurposed drugs that block the interaction between the SARS-CoV-2 spike protein and its receptor ACE2 could offer a rapid route to novel COVID-19 treatments or prophylactics. Here, we screened 2701 compounds from a commercial library of drugs approved by international regulatory agencies for their ability to inhibit the binding of recombinant, trimeric SARS-CoV-2 spike protein to recombinant human ACE2. We identified 56 compounds that inhibited binding by <90%, measured the EC_50_ of binding inhibition, and computationally modeled the docking of the best inhibitors to both Spike and ACE2. These results highlight an effective screening approach to identify compounds capable of disrupting the Spike-ACE2 interaction as well as identifying several potential inhibitors that could serve as templates for future drug discovery efforts.

## Introduction

COVID-19 is currently a global pandemic, causing extensive mortality and economic impact. While the success of rapidly developed vaccines offers hope to control the virus^1^, treatments that improve disease outcomes are also critically needed. Remdesivir, a nucleoside inhibitor of viral RNA polymerase, is the only drug approved by the FDA to treat COVID-19^2^, though its efficacy is disputed^3^. Monoclonal antibodies^4^ or recombinant Ace2^5^, which block the interaction between the SARS2 spike protein and its obligatory receptor ACE2, have also shown promise, but are expensive and suffer production limitations. Repurposing already-approved small molecule drugs could allow for rapid deployment of low-cost and widely available therapeutics^6^, but candidates studied in robust clinical trials have thus far failed^3,7,8^. Here, we took an unbiased approach to screen 2701 drugs approved by global regulatory agencies for the ability to block the interaction between recombinant, trimeric SARS2 spike protein^9^ and latex-bead-conjugated recombinant human ACE2.

## Methods

### Drug Screening

The “FDA-approved drug screening library” (Cat # L1300) was purchased from Selleck Chemicals. In a 96-well plate format, we briefly incubated recombinant, biotinylated, trimeric spike protein^9^ with either 200uM or 1mM of each drug, in duplicate, then added 5-micron flow cytometry beads (Luminex) coated with recombinant ACE2 (for detailed methods, see^10^). Three replicates per plate of positive (vehicle) controls and negative no-spike-protein controls were included, 31 plates in total. After washing and the addition of streptavidin-PE to bind spike attached to ACE2, plates were washed again on a magnetic plate washer and read on an Acea Novocyte flow cytometer. Data were expressed as the median PE fluorescence intensity (MFI), and converted to % inhibition using the formula 1-(MFI_drug_/MFI_positive control_). For EC_50_ studies, serial dilutions of drugs were performed, in duplicate, and run as above.

### In Silico Modeling

The S1 subunit of SARS-CoV-2 spike protein comprising the receptor binding domain (RBD) was modelled using the SWISS-MODEL server^11^, the FASTA sequence (residues 316-530) was retrieved from UniProtKB - P0DTC2 and used as a query sequence, PDB ID: 6VSB with 100% sequence identity was used as a template^12,13^. The structure quality of the modelled protein was validated on PROCHECK, which showed 89.2% residues in the core regions, the quality factor of 89.77% was obtained from Verify 3D on SAVES server and ProSA web gave a Z-score of -6.15 ^14–16^. The crystal structure of ACE2 protein was retrieved from PDB 1D: 2AJF and used for docking into the RBD interface. The SDF structures of the selected FDA-approved drugs were downloaded from Selleck chemicals and PubChem. Ligand and protein preparation was performed using the Ligprep and protein preparation wizard tool on Schrödinger Maestro version 12.2. Structural-based docking to the RBD interface of the Spike and ACE2 was also performed using Schrödinger Maestro and BIOVIA Discovery Studio (Dassault Systems) for docking analysis and visualization.

## Results

We took an unbiased approach to screen 2701 drugs approved by global regulatory agencies for the ability to block the interaction between recombinant, trimeric SARS2 spike protein^9^ and latex-bead-conjugated recombinant human ACE2 (**Figure 1A**). The use of a cell-free system prevented potential cytotoxic effects of drugs inherent to cell-culture based live-virus assays, and allowed us to focus solely on inhibition of the Spike-ACE2 interaction. Using a previously validated assay^10^, we quantified the amount of Spike-ACE2 co-association in the presence of high concentrations (200uM-1mM) of each drug, in duplicate, and calculated the percent inhibition using six replicates of vehicle control per plate (31 plates total). In this first-round screen, 114 drugs that exhibited 90% or greater inhibition were identified (**Figure 1B** and **Table S1**).

**Figure 1:**
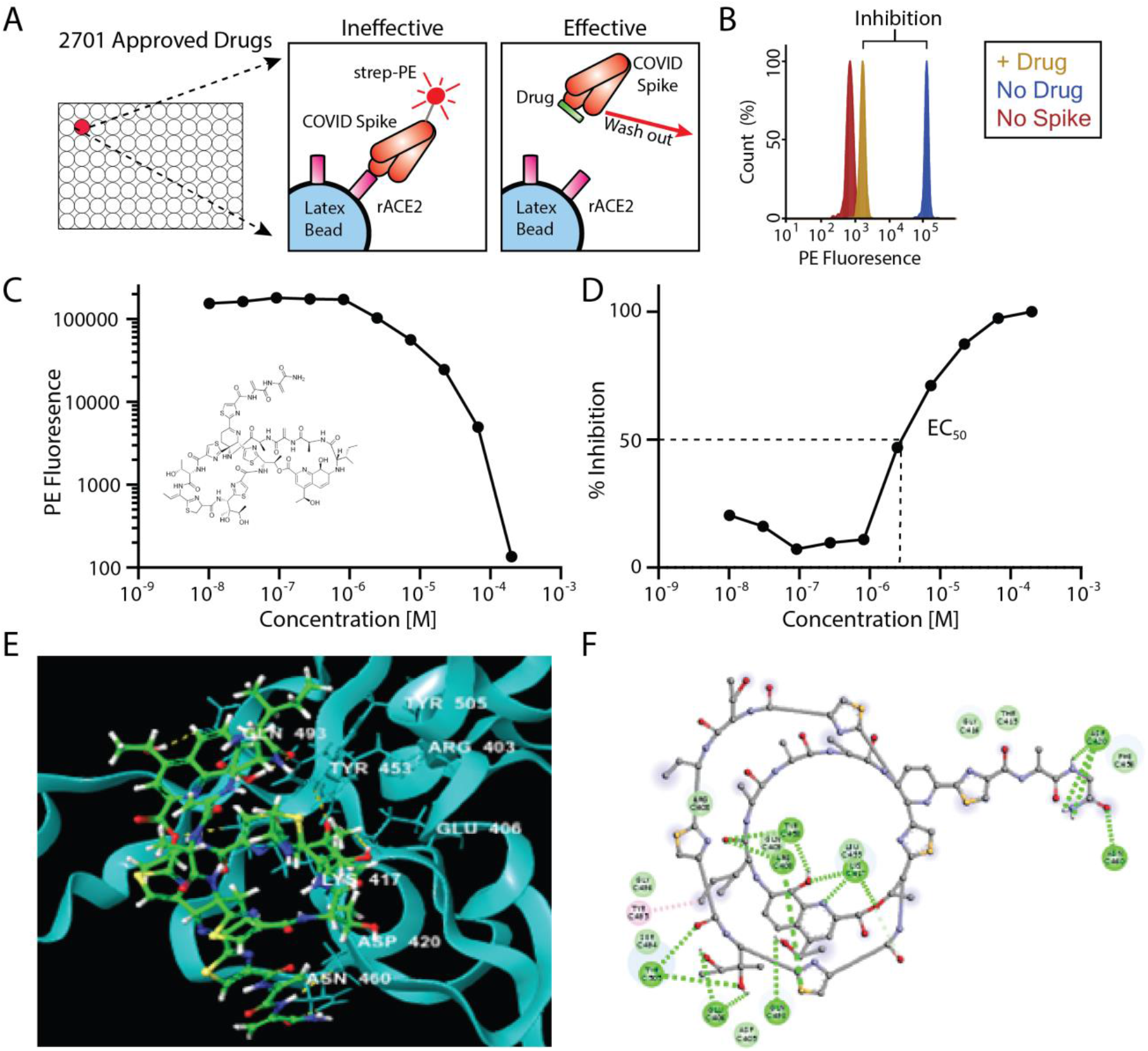
Inhibition of Spike-ACE2 binding by repurposed drugs. A) In vitro assay design showing inhibition of ACE2-Spike binding by “effective” drugs. B) Example histogram from primary screen showing >90% inhibition of ACE2-Spike binding following drug addition. C, D) EC_50_ data for the top candidate, Thiostrepton (structure inset) expressed as PE fluorescence (C) or percent inhibition (D). E) Three-dimensional and F) two-dimensional computational rendering of Thiostrepton binding to the SARS-CoV-2 spike protein.

We next performed serial dilutions of these 114 drugs, in duplicate, to measure EC_50_s of the ACE2 – spike interaction (**Figure 1C, D**). Fifty-eight of the drugs were revealed to be either false positive hits (they showed no inhibition upon re-screening), or showed inhibition only at the highest concentration tested, and were eliminated. The drug with the highest EC_50_ was Thiostrepton (EC_50_ = 3.95 x 10^−6^ M), a cyclic oligopeptide used as a topical antibiotic in animals, and that interacts with the transcription factor FOXM1 to inhibit the growth of breast cancer cells in vitro^17^. Next was Oxytocin (EC_50_ = 4.17 ⨯ 10^−6^ M), a peptide hormone that is administered to induce childbirth, and that may increase social cognition when administered intranasally^18^. The next four best candidates were actually two closely related pairs of drugs, which demonstrates the robustness of our screen in identifying each compound twice. Nilotinib (EC_50_ = 4.20 ⨯ 10^−6^ M), which was identified as both a free base and an HCl salt, is a selective tyrosine kinase inhibitor used to treat chronic myelogenous leukemia^19^.

Hydroxycamptothecine (EC_50_ = 7.25 ⨯ 10^−6^ M) and its stereoisomer S-10-Hydroxycamptothecine (EC_50_ = 7.07 × 10^−6^ M) are DNA topoisomerase I inhibitors with anti-cancer activity^20^. Interestingly, three derivatives that have also been approved for cancer therapy, Topotecan, Irinotecan, and Belotecan were included in the screening panel, but did not inhibit spike-ACE2 binding. The EC_50_s of all 56 compounds are listed in **Table S2**; given the decreased severity of COVID-19 in females^21^, it is notable that Estradiol Benzoate (a synthetic estrogen) inhibited the interaction (EC_50_ = 1.75 ⨯ 10^−5^ M).

We next performed molecular docking studies with both spike and ACE2 to unravel the binding and crucial molecular interactions of the top 12 candidates, focusing on the Spike-Ace2 binding interface. The compounds with the lowest EC_50_s also gave the lowest docking (Glide^22^) scores (**Table 1**), cross-validating our results. The two top hits, Thiostrepton and Oxytocin, bound more favorably to Spike than Ace2, and interacted with several key residues that mediate Spike-ACE2 binding^23^ **(Figure 1, Figure S1-S2**, and **Table S3)**. Thiostrepton in particular bound extensively to Spike residues, and an OH-group in Thiostrepton bound simultaneously with Lys417 of Spike and Asp30 of ACE2. Simultaneous binding of these two critical interface resides would likely disrupt the Spike-ACE2 interaction, resulting in the low observed EC50 value. Nilotinib and Hydroxycamptothecine exhibited glide scores that were slightly more favorable for ACE2, but also showed moderate binding with Spike. Interestingly, both interacted with Arg393 on ACE2, a critical spike-binding residue^16^, and both also interacted with the Spike receptor binding interface (**Figure S3-S6** and **Table S3**). The remaining compounds generally gave higher glide scores, consistent with their lower affinity, but still docked appreciable to ACE2 and/or Spike (see **Figure S7-S12, Table S3** and **Supplementary Discussion**).

**Table 1:** Summary of the top 12 drug candidates. Computationally modeled glide scores for ACE2 and spike binding and EC_50_ values measured with the recombinant Spike-ACE2 binding inhibition assay are displayed.

|  | <u>Glide Score</u> |  | EC<sub>50</sub> |
| --- | --- | --- | --- |
|  | Spike | Ace2 |  |
| Thiostrepton | -7.173 | -4.819 | 3.95E-06 |
| Oxytocin | -7.024 | -5.205 | 4.17E-06 |
| Nilotinib AMN-107 | -5.669 | -6.207 | 4.20E-06 |
| S-(10)-Hydroxycamptothecin | -5.335 | -5.673 | 7.07E-06 |
| Hydroxycamptothecin | -5.321 | -5.679 | 7.25E-06 |
| Nilotinib HCl | -5.659 | -6.209 | 8.38E-06 |
| Selamectin | -4.207 | -3.503 | 8.46E-06 |
| Picropodophyllin | -4.072 | -4.158 | 9.97E-06 |
| Docetaxel | -4.737 | -5.359 | 1.03E-05 |
| Doramectin | -4.65 | -3.964 | 1.28E-05 |
| Anidulafungin | -4.649 | -5.846 | 1.31E-05 |
| Estradiol Benzoate | -4.003 | -5.345 | 1.73E-05 |

## Discussion

Overall, this study identified 56 approved drugs that show some efficacy in blocking the interaction between the COVID spike protein and its receptor, ACE2. Many of the identified drugs are already approved for clinical use in humans (see **Table 2**). However, aiming towards clinical translation, there are several obvious issues. First, many of the drugs are chemotherapy agents, and have toxic side-effects that would not be tolerable in COVID-19 patients. For example, Nilotinib inhibits a kinase important in B cell signaling^19,24^, and may prevent normal immune function, while Picropodophyllin^25^ and Docetaxel^26^ both produce moderate to severe side effects when used in the context of chemotherapy. Secondly, several of the drugs have peak plasma concentrations that are orders of magnitude lower than the concentration required to inhibit spike-Ace2 binding in the in vitro assay, for example Oxytocin^27^ and Doramectin^28^ (0.005 vs 4.2 uM and 0.014 vs 12.8 uM, respectively). Finally, Oxytocin’s clearance rate would necessitate continuous infusion, which seems impractical.

**Table 2:** Reported pharmacokinetic properties of top hits^24–33^. Drugs in grey indicate dosing studies were performed on non-human animals.

|  | <u>Indication</u> | <u>C<sub>max</sub> (ng/ml)</u> | <u>MW (g/mol)</u> | <u>C<sub>max</sub> (uM)</u> | <u>EC<sub>50</sub> (uM)</u> | <u>Elimination T<sub>1/2</sub></u> | <u>Dosing Regimen</u> | <u>Reference</u> |
| --- | --- | --- | --- | --- | --- | --- | --- | --- |
| Thiostrepton | Topical antibiotic, vetrinary |  | 1664 |  | 4.0 |  |  | N/A |
| Oxytocin | Modify social behavior (expeirmental) | 0.005 | 1007 | 0.005 | 4.2 | >1 hour | Intranasal Single dose, 44ug | Gossen et al 2012 |
|  | Induction of labor | 0.005 | 1007 | 0.005 | 4.2 | >>1 hour | IV infusion 6.7ng/min | Leake et al 1980 |
| Nilotinib | Kinase inhibitor, Cancer treatment | 1360 | 529 | 2.57 | 4.2 | 16 hours | Oral BID 300mg | Tian et al 2018 |
| Hydroxycamptothecin (in Rats) |  | 15930 | 364.4 | 43.7 | 7.3 | 428 min | RATS 10mg/kg IV | Zhang 1998 |
| Selamectin (in Dogs) |  | 86.5 | 770 | 0.11 | 8.5 | 266 hours | DOGS topical 24 mg/kg | Sarasola 2002 |
|  |  | 7630 | 770 | 9.9 | 8.5 | 45.7 hours | DOGS oral 24 mg/kg | Sarasola 2002 |
| Picropodophyllin (PPP) | IGF inhibitor, Cancer treatment (aka AXL1717) | 207-1035 | 414 | 0.5-2.5 | 10.0 | 2 hours | Oral 390 or 520 mg BID | Ekman et al 2015 |
| Docetaxel | Microtubule inhibitor, Cancer treatment | 933 | 808 | 1.23 | 10.3 | 25.4 hours | IV infusion 30-36 mg/m<sup>2</sup> | Gustafson 2003 |
| Doramectin | Antiparasitic, vetrinary | 12.2 | 899 | 0.0135 | 12.8 | 10 days | CATTLE topical 500 ug/kg | Gayrard et al 1999 |
| Anidulafungin | Antifungal | 2500 | 1140 | 2.2 | 13.1 | 27 hours | IV infusion 100mg | Wasmann 2018 |
| Estradiol Benzoate | Contraceptive | 0.75 | 376 | 0.002 | 17.3 | 3 days | IM injection, 5mg | Oriowo et al 1980 |

In light of known side-effects and pharmacokinetic data of many of our high-ranking drug candidates, Selamectin may be the top candidate for further study. Selamectin is an anti-parasitic used in dogs and cats to prevent infestation with nematode and arthropod species. Oral Selamectin is well tolerated in dogs and can achieve a peak plasma concentrations (9.9uM) comparable to the measured EC_50_ (8.5 uM)^29^. However, we were not able to identify any studies using Selamectin in humans, since Ivermectin is the standard alternative. Thiostrepton would also be a top candidate, but pharmacokinetic data are lacking.

An advantage of our recombinant approach, using a cell-free system, is that the effects of drug toxicity on cell growth do not confound the readout of Spike-Ace2 binding. However, our results would need to be replicated in a cell or animal model using live virus to ensure that anti-SARS effects appear at drug concentrations low enough to prevent toxicity. The relatively weak (micromolar) binding kinetics of the drugs identified here, as well as their known toxicities, bioactivities and/or high clearance rates, suggest that many would currently be unlikely to be viable for treating acute disease or for prophylactic use. However, they could serve as starting points for future medicinal chemistry optimization efforts to rationally design derivatives that are both less toxic and bind to the COVID spike with higher affinity.

## Supporting information

Supplemental Figure Legends and Discussion

## Acknowledgements

The authors thank Jason McLellan and Barney Graham for their generous gift of the SARS-CoV-2 recombinant trimer and hACE2 expression constructs; This study was funded by a COVID-19 research award from the University of Washington Institute for Translational Heath Sciences (via grant UL1 TR002319) and by a COVID19 award from the Research Integration Hub at Seattle Children’s Research Institute (to SEPS), and by R01 AI140951 (to DNS). The authors declare no conflicts of interest.

## Author Contributions

SEPS conceived the study. KT, CA, EPG, JW and SEPS planned experiments. NS provided recombinant COVID proteins for the inhibition assay. KT and EPG performed microsphere-based experiments, CA performed modeling studies under the supervision of JW. KT, CA, JW and SEPS, analyzed the data. KT, CA, JW and SEPS wrote the manuscript, and all authors read and approved the manuscript.

## Notes

### Competing Interest Statement

SCRI has submitted a provisional patent on the assay and compounds described in this manuscript.

## References

1. Polack, F. P. et al. Safety and Efficacy of the BNT162b2 mRNA Covid-19 Vaccine. N. Engl. J. Med. 383, 2603–2615 (2020).

2. Beigel, J. H. et al. Remdesivir for the Treatment of Covid-19 - Final Report. N. Engl. J. Med. 383, 1813–1826 (2020).

3. Repurposed Antiviral Drugs for Covid-19 — Interim WHO Solidarity Trial Results ä NEJM. N. Engl. J. Med.

4. Marovich, M., Mascola, J. R. & Cohen, M. S. Monoclonal Antibodies for Prevention and Treatment of COVID-19. JAMA 324, 131 (2020).

5. Monteil, V. et al. Inhibition of SARS-CoV-2 Infections in Engineered Human Tissues Using Clinical-Grade Soluble Human ACE2. Cell 181, 905-913.e7 (2020).

6. Saul, S. & Einav, S. Old Drugs for a New Virus: Repurposed Approaches for Combating COVID-19. ACS Infect. Dis. 6, 2304–2318 (2020).

7. Cao, B. et al. A Trial of Lopinavir–Ritonavir in Adults Hospitalized with Severe Covid-19. N. Engl. J. Med. (2020) doi:10.1056/NEJMoa2001282.

8. Hydroxychloroquine in patients mainly with mild to moderate COVID–19: an open– label, randomized, controlled trial ä medRxiv. https://www.medrxiv.org/content/10.1101/2020.04.10.20060558v2.

9. Wrapp, D. et al. Cryo-EM structure of the 2019-nCoV spike in the prefusion conformation. Science 367, 1260–1263 (2020).

10. Gniffke, E. P. et al. Plasma From Recovered COVID-19 Patients Inhibits Spike Protein Binding to ACE2 in a Microsphere-Based Inhibition Assay. J. Infect. Dis. 222, 1965–1973 (2020).

11. Waterhouse, A. et al. SWISS-MODEL: homology modelling of protein structures and complexes. Nucleic Acids Res. 46, W296–W303 (2018).

12. Pundir, S., Martin, M. J., O’Donovan, C., & UniProt Consortium. UniProt Tools. Curr. Protoc. Bioinforma. 53, 1.29.1-1.29.15 (2016).

13. Berman, H. M. et al. The Protein Data Bank. Nucleic Acids Res. 28, 235–242 (2000).

14. Laskowski, R. A., MacArthur, M. W., Moss, D. S. & Thornton, J. M. PROCHECK: a program to check the stereochemical quality of protein structures. J. Appl. Crystallogr. 26, 283–291 (1993).

15. Eisenberg, D., Lüthy, R. & Bowie, J. U. VERIFY3D: assessment of protein models with three-dimensional profiles. Methods Enzymol. 277, 396–404 (1997).

16. Wiederstein, M. & Sippl, M. J. ProSA-web: interactive web service for the recognition of errors in three-dimensional structures of proteins. Nucleic Acids Res. 35, W407–410 (2007).

17. Hegde, N. S., Sanders, D. A., Rodriguez, R. & Balasubramanian, S. The transcription factor FOXM1 is a cellular target of the natural product thiostrepton. Nat. Chem. 3, 725–731 (2011).

18. Keech, B., Crowe, S. & Hocking, D. R. Intranasal oxytocin, social cognition and neurodevelopmental disorders: A meta-analysis. Psychoneuroendocrinology 87, 9– 19 (2018).

19. Tokuhira, M. et al. Efficacy and safety of nilotinib therapy in patients with newly diagnosed chronic myeloid leukemia in the chronic phase. Med. Oncol. Northwood Lond. Engl. 35, 38 (2018).

20. Fei, B., Chi, A. L. & Weng, Y. Hydroxycamptothecin induces apoptosis and inhibits tumor growth in colon cancer by the downregulation of survivin and XIAP expression. World J. Surg. Oncol. 11, 120 (2013).

21. Peckham, H. et al. Male sex identified by global COVID-19 meta-analysis as a risk factor for death and ITU admission. Nat. Commun. 11, 6317 (2020).

22. Friesner, R. A. et al. Extra precision glide: docking and scoring incorporating a model of hydrophobic enclosure for protein-ligand complexes. J. Med. Chem. 49, 6177–6196 (2006).

23. Lan, J. et al. Structure of the SARS-CoV-2 spike receptor-binding domain bound to the ACE2 receptor. Nature 581, 215–220 (2020).

24. Tian, X. et al. Clinical Pharmacokinetic and Pharmacodynamic Overview of Nilotinib, a Selective Tyrosine Kinase Inhibitor. J. Clin. Pharmacol. 58, 1533–1540 (2018).

25. Ekman, S. et al. A novel oral insulin-like growth factor-1 receptor pathway modulator and its implications for patients with non-small cell lung carcinoma: A phase I clinical trial. Acta Oncol. 55, 140–148 (2016).

26. Gustafson, D. L. et al. Analysis of docetaxel pharmacokinetics in humans with the inclusion of later sampling time-points afforded by the use of a sensitive tandem LCMS assay. Cancer Chemother. Pharmacol. 52, 159–166 (2003).

27. Gossen, A. et al. Oxytocin plasma concentrations after single intranasal oxytocin administration -a study in healthy men. Neuropeptides 46, 211–215 (2012).

28. Gayrard, V., Alvinerie, M. & Toutain, P. L. Comparison of pharmacokinetic profiles of doramectin and ivermectin pour-on formulations in cattle. Vet. Parasitol. 81, 47–55 (1999).

29. Sarasola, P. et al. Pharmacokinetics of selamectin following intravenous, oral and topical administration in cats and dogs. J. Vet. Pharmacol. Ther. 25, 265–272 (2002).

30. Leake, R. D., Weitzman, R. E. & Fisher, D. A. Pharmacokinetics of oxytocin in the human subject. Obstet. Gynecol. 56, 701–704 (1980).

31. Zhang, R. et al. Preclinical pharmacology of the natural product anticancer agent 10-hydroxycamptothecin, an inhibitor of topoisomerase I. Cancer Chemother. Pharmacol. 41, 257–267 (1998).

32. Wasmann, R. E. et al. Pharmacokinetics of Anidulafungin in Obese and Normal-Weight Adults. Antimicrob. Agents Chemother. 62, (2018).

33. Oriowo, M. A., Landgren, B. M., Stenström, B. & Diczfalusy, E. A comparison of the pharmacokinetic properties of three estradiol esters. Contraception 21, 415–424 (1980).

