## Supplemental Figure Legends and Discussion for "A repurposed drug screen identifies compounds that inhibit the binding of the COVID-19 spike protein to ACE2"

**-for-**

**Figures S1-11.pptx**

**Tables S1-S3.xlsx**

**Supplementary Discussion**

**File “Figures S1-11.pdf”**

**Figure legends included in the file**

**File “Table S1.xlsx”**

**Table S1: Results of primary screen of 2701 compounds from the Selleckchem “FDA approved drug library”.** Compounds are listed with their plate and well location, and the percent inhibition in the initial screen, measured in duplicate, is presented. Compounds that met the primary screening criteria of >90% inhibition are highlighted in red.

**File “Table S2.xlsx”**

**Table S2: Results of the $EC_{50}$ screening showing the 56 compounds that produced measurable $EC_{50}$ values.** Compounds are listed with their plate and well location, the percent inhibition in the initial screen (measured in duplicate), and the EC50 value measured by serial dilution, in duplicate, and computed in Prizm Graphpad 9.0. The list is sorted by $EC_{50}$, with the most strongly inhibiting compounds on top.

**File “Table S3.xlsx”**

**Table S3: Computationally modeled binding interactions between the top 12 candidates (by $EC_{50}$) and Spike and ACE2 protein residues.** Interactions between drug candidates and protein targets were modeled using Schrödinger Maestro version 12.2. Interactions are listed by bond type, and resides that are critical to the Spike-ACE2 binding [cite Lan 2020] are highlighted in bold type. Critical residues include: **On spike:** Lys417, Tyr449, Tyr453, Leu455, Phe456, Gln474, Phe486, Asn487, Tyr489, Phe490, Gln493, Ser494, Gly496, Gln498, Thr500, Asn501, Gly502 and Tyr505.[12] **On ACE2**: Gln24, Asp30, Lys31, His34, Glu35, Glu37, Asp38, Tyr41, Gln42, Met82, Tyr83, Lys353, Gly354 and Arg393.[11-13]

**Supplemental Discussion: Detailed Results and Discussion related to Computational Modeling studies.**

Molecular docking studies were performed on the top 12 drug candidates to evaluate their potential binding and molecular interactions with the Spike protein of SARS-CoV-2 and human ACE2 . Residues actively participating in the interaction between the receptor binding domain of Spike and ACE2[1] were targeted and used for structure-based screening. The important S-protein RDB residues at the interface are: Lys417, Tyr449, Tyr453, Leu455, Phe456, Gln474, Phe486, Asn487, Tyr489, Phe490, Gln493, Gly496, Ser494, Gln498, Thr500, Asn501, Gly502 and Tyr505[1]. The crucial residues in ACE2 involved in binding to the Spike protein include: Gln24, Asp30, Lys31, His34, Glu35, Glu37, Asp38, Tyr41, Gln42, Met82, Tyr83, Lys353, Gly354 and Arg393[1,2]. The docking experiments were generally consistent with the experimentally obtained binding data, with compounds displaying the lowest $EC_{50}$s also giving the lowest docking scores (See **Table 1**). Thiostrepton and oxytocin yielded the lowest glide scores and displayed a higher affinity toward the Spike Receptor Binding Domain (S-RBD) residues by interacting with hotspot Lys417, which interact with Asp30 of ACE2 via hydrogen bond and salt bridge[1–3] (**Table S3**). These compounds also showed some interactions with ACE2. Conversely, nilotinib, docetaxel, anidulafungin and estradiol appear to prefer binding to ACE2 by interacting with Arg393 (**Table S3**), while S(10)-Hydroxycamptothecin and Hydroxycamptothecin, exhibited glide scores that were comparable for both spike and ACE2 protein.

Thiostrepton interacts with S-RBD residues with the lowest glide score (-7.173). It has the highest number of hydrogen bonds interacting with Arg403, Glu406, Lys417, Asp420, Tyr453, Asn460, Gln493, Tyr505 and an alkyl bond with Tyr495. Notably, H-bond interactions with Lys417, Tyr453, Gln493 and Tyr505, which are crucial residues in S-RBD[3], indicate strong interactions between Thiostrepton and spike glycoprotein (**Table S3, Figure 1**). The docking studies of Thiostrepton and ACE2 revealed hydrogen bonding interactions with Asp30, Asp38 and Arg393. Intriguingly, the same OH-group in

Thiostrepton binds with Lys417 of Spike protein, and also binds with Asp30 of ACE2 (**Table S3 and Figure S1**). Simultaneous binding to two critical resides would likely cause a major disruption in the Spike-ACE2 interaction, thereby resulting in the lowest EC50 value. Van der Waals interactions with Lys353 residues in ACE2 were also observed. These data demonstrate that Thiostrepton can bind and block the interaction between Spike and ACE2 at the S-RBD interface.

Oxytocin showed similar interactions as Thiostrepton (**Figure S2**). The S-RBD docking results (-7.024) showed molecular interactions between Oxytocin and important S-RDB residues Lys417, Tyr449, Gln493, Gln498 and Try501 (**Table S3**). Oxytocin interacts with the ACE2 residues His34, Glu35, Glu42, and Glu75 *via* hydrogen bonds and forms an alkyl bond with Lys31. It also forms a salt bridge with Asp38.

Nilotinib AMN-107 and Nilotinib HCl have the same interactions with S-RBD, with glide scores of -5.669 and -5.659 respectively. Interactions with the crucial S-RBD residues Tyr449, Gln493, Tyr495, Gly496 and Asn501 were observed (**Table S3** and **Figures S3 and S6**). The pyridine, pyrimidine, and phenyl rings of Nilotinib formed π-π T-shaped interactions with Gly496, Tyr495, and Tyr449 of S-RBD, respectively. Interestingly, the fluoro groups formed halogen bonds with Phe490 and Leu492. Some van der Waals interactions with Ser494, Gln493, Tyr505, and Gln498 of S-protein RBD were observed. Nilotinib AMN-107 and Nilotinib HCl also showed similar interactions with ACE2, forming H-bonds with Arg393, π-π T-shaped bonds with Tyr349, and the three fluoro groups also formed halogen bond with Asp350 and Asp382 (**Table S3** and **Figures S3 and S6**).

S-(10)-Hydroxycamptothecin and Hydroxycamptothecin are stereoisomers, inhibitors of topoisomerase isolated from the Chinese tree *C. acuminate*. They both have similar binding poses for S-RBD and ACE2 (**Figures S4 and S5)**. The docked compounds interact with Arg403, Gln414, and Lys417 of the spike protein, with hydrogen bond interactions between the carbonyl groups and Lys417. S-(10)-Hydroxycamptothecin and Hydroxycamptothecin bind to Asp30 and Glu37 of ACE2 *via* hydrogen bonds, and to Gln388 and Arg393 via π-stacked and π-cation bond interactions, respectively.

Selamectin is a macrocyclic lactone used as an antiparasitic in dogs and cats. Selamectin binds with S-RBD residues via hydrogen bond interactions with Arg403, Gln409, Gln414,

Lys417, Try453, and an alkyl bond with Lys417 (**Table S3** and **Figure S7)**. Note that Lys417 and Try453 have a crucial role in the interaction of S-RBD and ACE2. Selamectin also interacts with ACE2 through hydrogen bonds with residues Asp30, Gly354, Arg393 and Lys353, which is a hotspot in RBD interface. Interaction with His34 *via* π-alkyl bond was also observed (**Table S3** and **Figure S7**).

Picropodophyllin is a plant-derived inhibitor of the insulin-like growth factor-1 receptor, and has antineoplastic activity. In docking with S-RBD, it forms hydrogen bonds with Lys458 and Gln474, and displays a π-stacked interaction with Ala475 and a π-cation interaction with Lys458. Note that the binding involves Gln474 and Ala475, which are crucial for S-RBD and ACE2 interactions (**Figure S8**). Picropodophyllin also binds to Asp350 and Phe40 of ACE2 through hydrogen bond and π-stacked interactions, respectively (**Table S3** and **Figure S8**).

Docetaxel is a semi synthetic analogue of paclitaxel (taxol), which is a strong chemotherapy agent used to treat a number of different cancers such as head and neck cancer, breast cancer, stomach cancer, prostate cancer and non-small-cell lung cancer. The docking pose with S-RBD showed interactions with residues Tyr449, Gln493, Asn501, and an alkyl bond with Tyr449 (**Table S3** and **Figure S9**). All of these residues play important roles in the Spike-ACE2 binding and may inhibit the interaction. The docking pose of Docetaxel with ACE2 showed hydrogen bond interactions with Asn394, Arg393 and Lys562 residues (**Table S3** and **Figure S9**).

Doramectin is a macrocyclic lactone is used for the treatment of gastrointestinal roundworms, eyeworms, lungworms, grubs, sucking lice and mange mites in cattle. It showed hydrogen bond interactions with S-RBD residues Ile468, Thr470, Leu492, and Ser494. We observed an alkyl bond with Leu452 and Leu455, and a π-sigma interaction with Phe490. Notably, Ser494 on S-RBD provides support for Lys353 – a crucial residue in ACE2 because it interacts with Tyr505, Gly502, Gly496, Asn501 of the Spike protein[2]. Doramectin also binds to ACE2 at Gly354, which is an adjacent residue to Lys353 and implies a mechanism of Spike-ACE2 inhibition (**Table S3** and **Figure S10**).

Anidulafungin, which was previously known as LY303366, is an antifungal antibiotic that is used to treat yeast infections. It showed hydrogen bond interactions with Asp405 and

Arg408 of S-RBD, and displayed π- π T shaped and π-alkyl bond interactions with Tys505 and Tyr449 respectively. Anidulafungin also interacts with ACE2 via Glu35, Glu75, Lys353 and Ala386 (**Table S3** and **Figure S11**)

Estradiol benzoate is an estrogen analogue used for menopausal symptoms as hormone therapy; it is also used in the treatment of prostate cancer in men. Estradiol interacts with S-RBD *via* hydrogen bonds with Phe490, π-alkyl interactions with Cys488 and an alkyl bond with Phe486. Phe486 and Phe490 have physiological roles in the Spike-ACE2 binding, since they make critical contacts with Tyr83 and Lys31 in ACE2[2]. Estradiol also showed hydrogen bond interactions with Gln102 and π-alkyl bond with Arg393 of ACE2 (**Table S3** and **Figure 12**).


1. Lan, J. *et al.* Structure of the SARS-CoV-2 spike receptor-binding domain bound to the ACE2 receptor. *Nature* **581**, 215–220 (2020).
2. Wan, Y., Shang, J., Graham, R., Baric, R. S. & Li, F. Receptor Recognition by the Novel Coronavirus from Wuhan: an Analysis Based on Decade-Long Structural Studies of SARS Coronavirus. *J. Virol.* **94**, (2020).
3. Yan, R. *et al.* Structural basis for the recognition of SARS-CoV-2 by full-length human ACE2. *Science* **367**, 1444–1448 (2020).
