## Supplementary material for "A repurposed drug screen identifies compounds that inhibit the binding of the COVID-19 spike protein to ACE2": Suppl. Figs 1-12

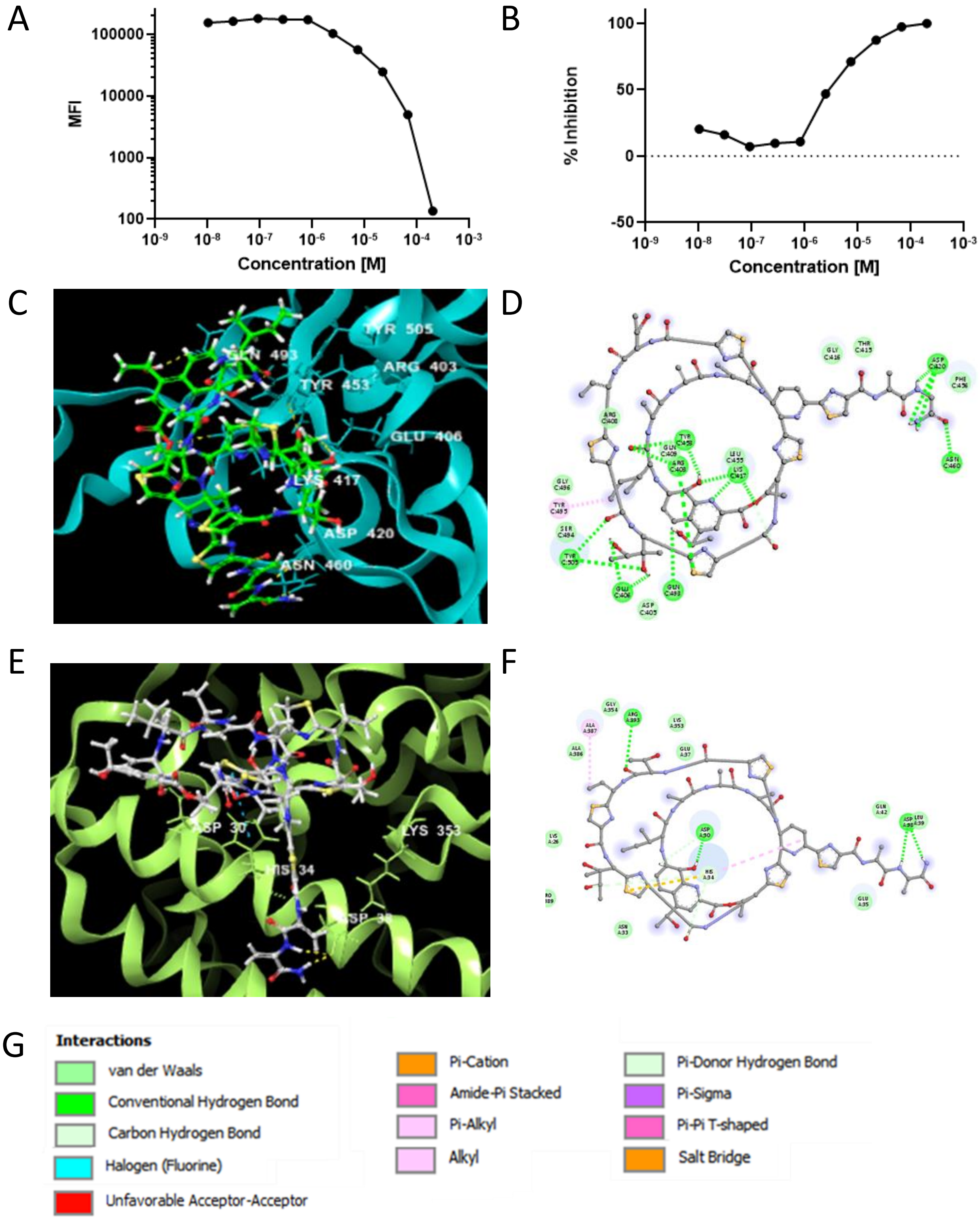


**Figure S1: EC50 and computational binding data for Thiostrepton**. A,B) EC50 data are expressed as median fluorescent intensity (MFI, A) or percent inhibition (B). C) Three dimensional and D) two dimensional computationally determined lowest-energy docking poses of Thiostrepton and the SARS-CoV-2 spike protein. E) Three dimensional and F) two dimensional computational docking of Thiostrepton and human ACE2. G) Key showing color-codes for interaction types. Note A-D are identical to Figure 1 C-F.

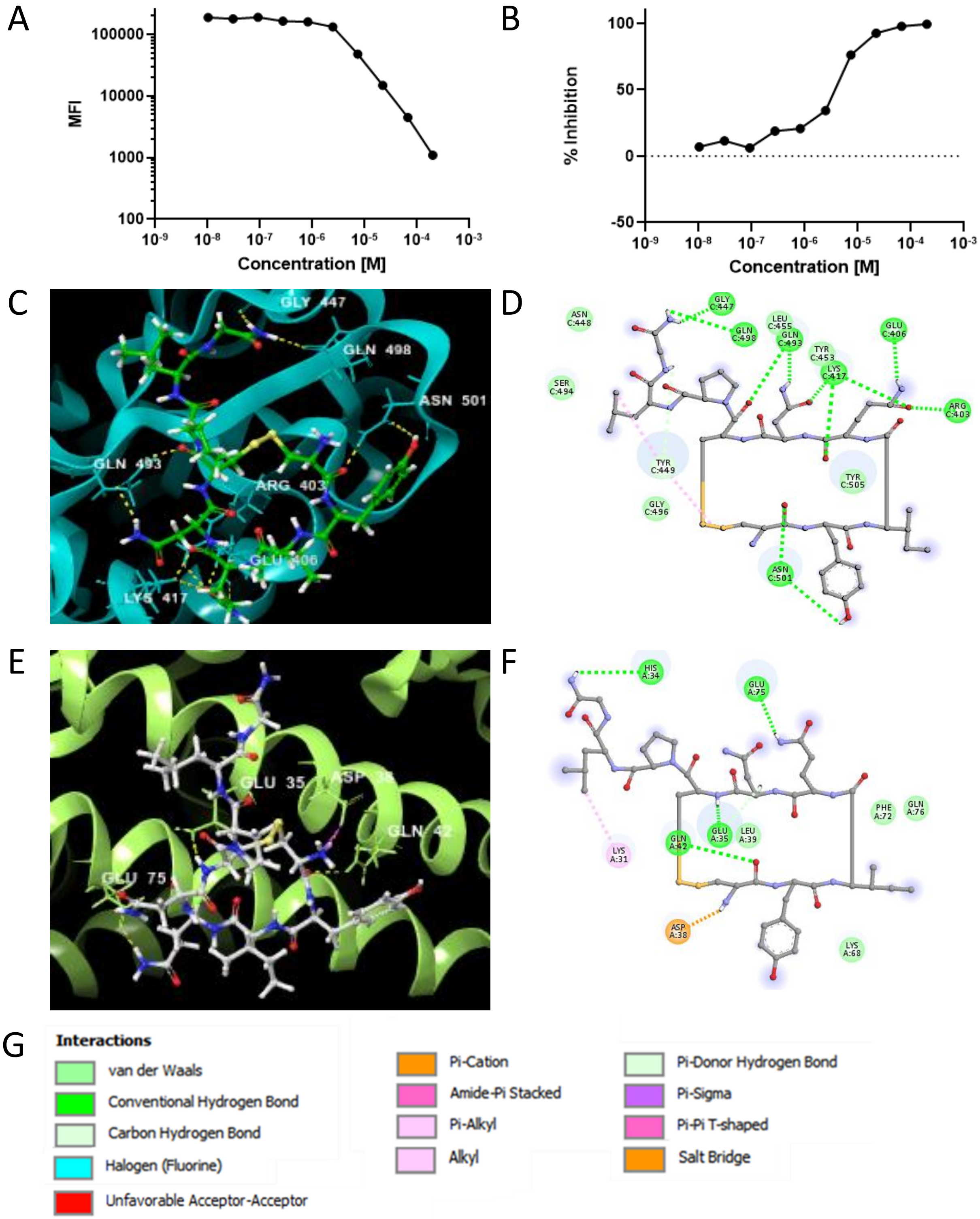


**Figure S2: EC50 and computational binding data for Oxytocin**. A,B) EC50 data are expressed as median fluorescent intensity (MFI, A) or percent inhibition (B). C) Three dimensional and D) two dimensional computationally determined lowest-energy docking poses of Oxytocin and the SARS-CoV-2 spike protein. E) Three dimensional and F) two dimensional computational docking of Oxytocin and human ACE2. G) Key showing color-codes for interaction types.

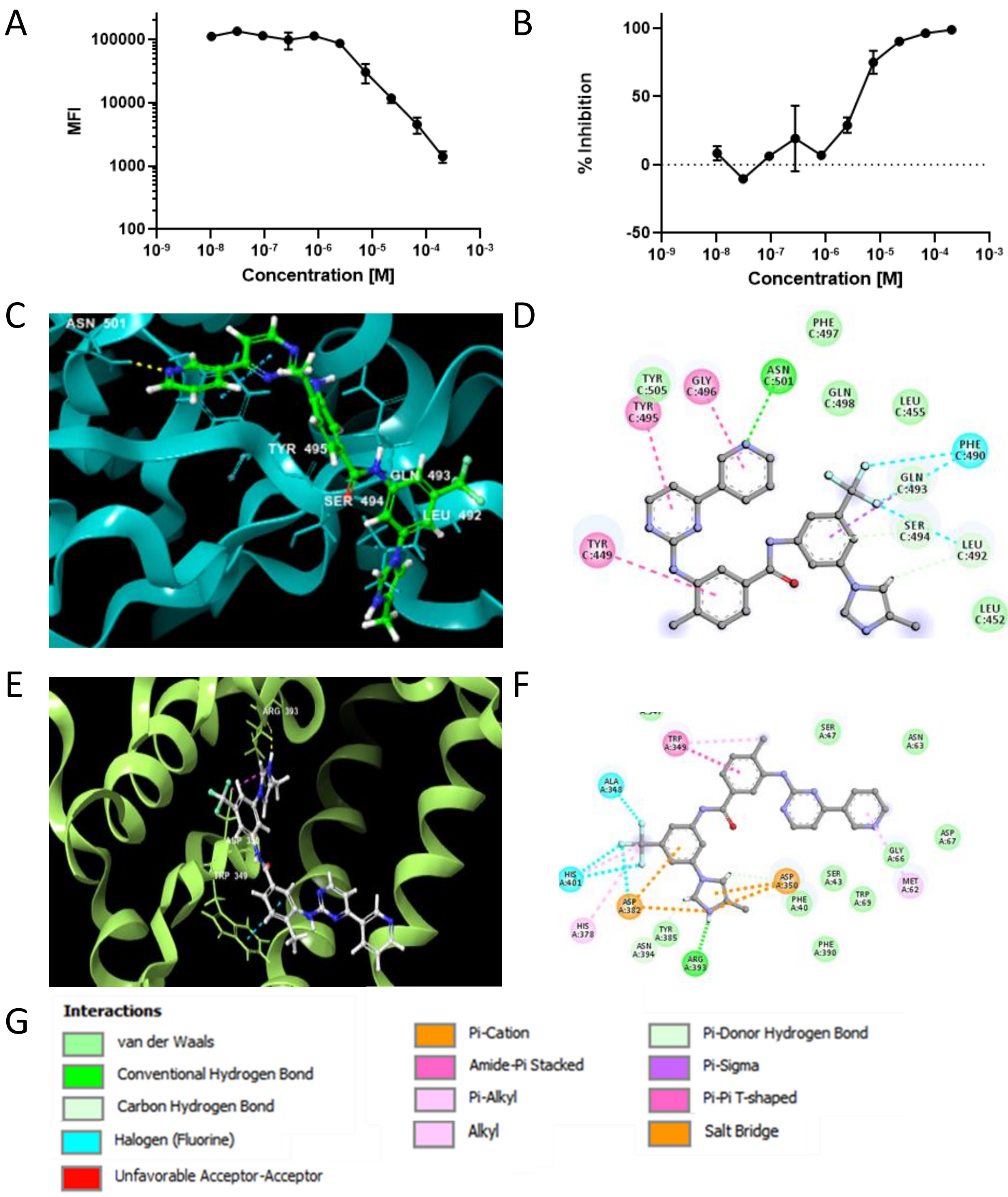


**Figure S3: EC50 and computational binding data for Nilotinib (AMN-107)**. A,B) EC50 data are expressed as median fluorescent intensity (MFI, A) or percent inhibition (B). C) Three dimensional and D) two dimensional computationally determined lowest-energy docking poses of Nilotinib and the SARS-CoV-2 spike protein. E) Three dimensional and F) two dimensional computational docking of Nilotinib and human ACE2. G) Key showing color-codes for interaction types.

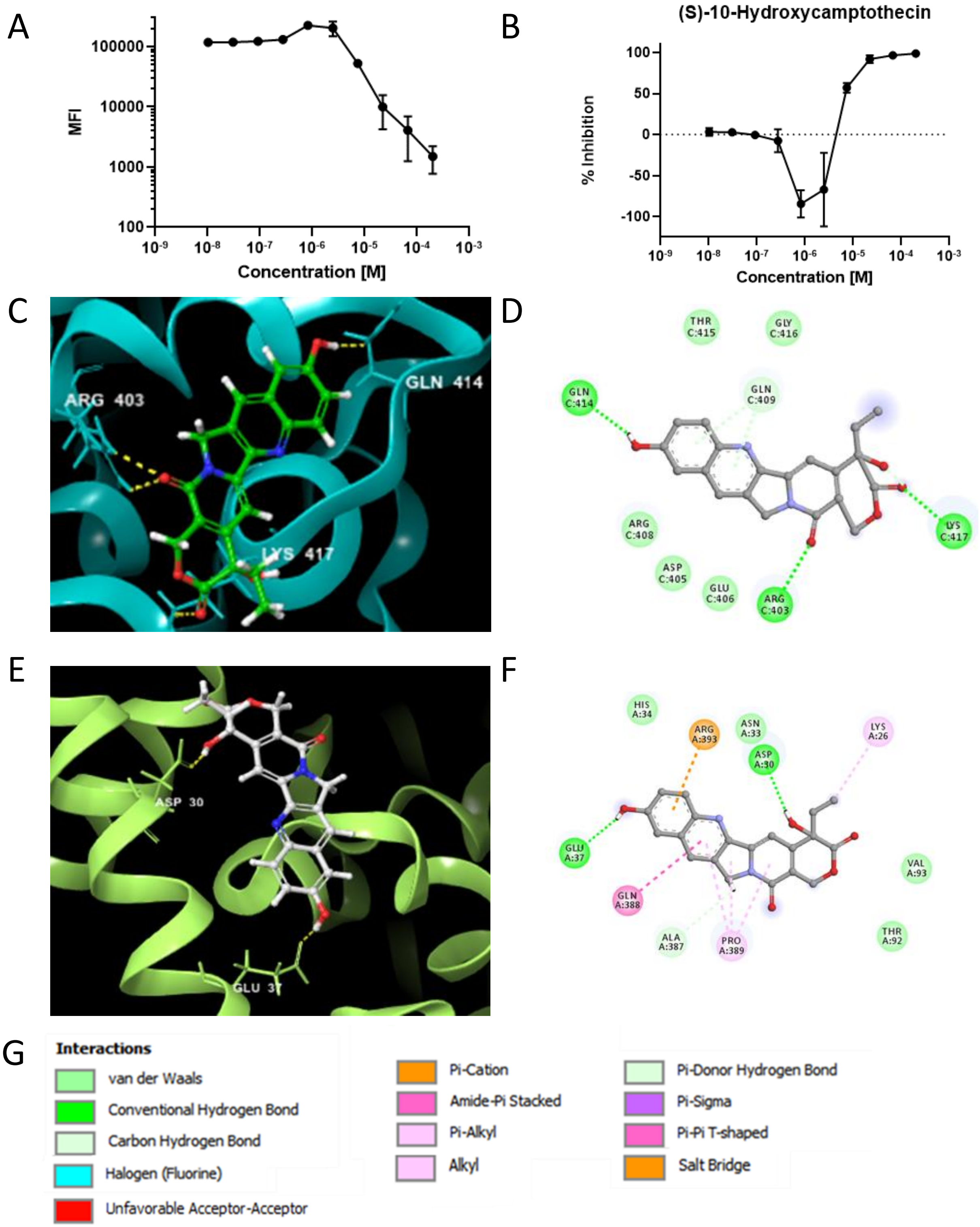


**Figure S4: EC50 and computational binding data for (S)-10-Hydroxycamptothecin**. A,B) EC50 data are expressed as median fluorescent intensity (MFI, A) or percent inhibition (B). C) Three dimensional and D) two dimensional computationally determined lowest-energy docking poses of (S)-10-Hydroxycamptothecin and the SARS-CoV-2 spike protein. E) Three dimensional and F) two dimensional computational docking of (S)-10-Hydroxycamptothecin and human ACE2. G) Key showing color-codes for interaction types.

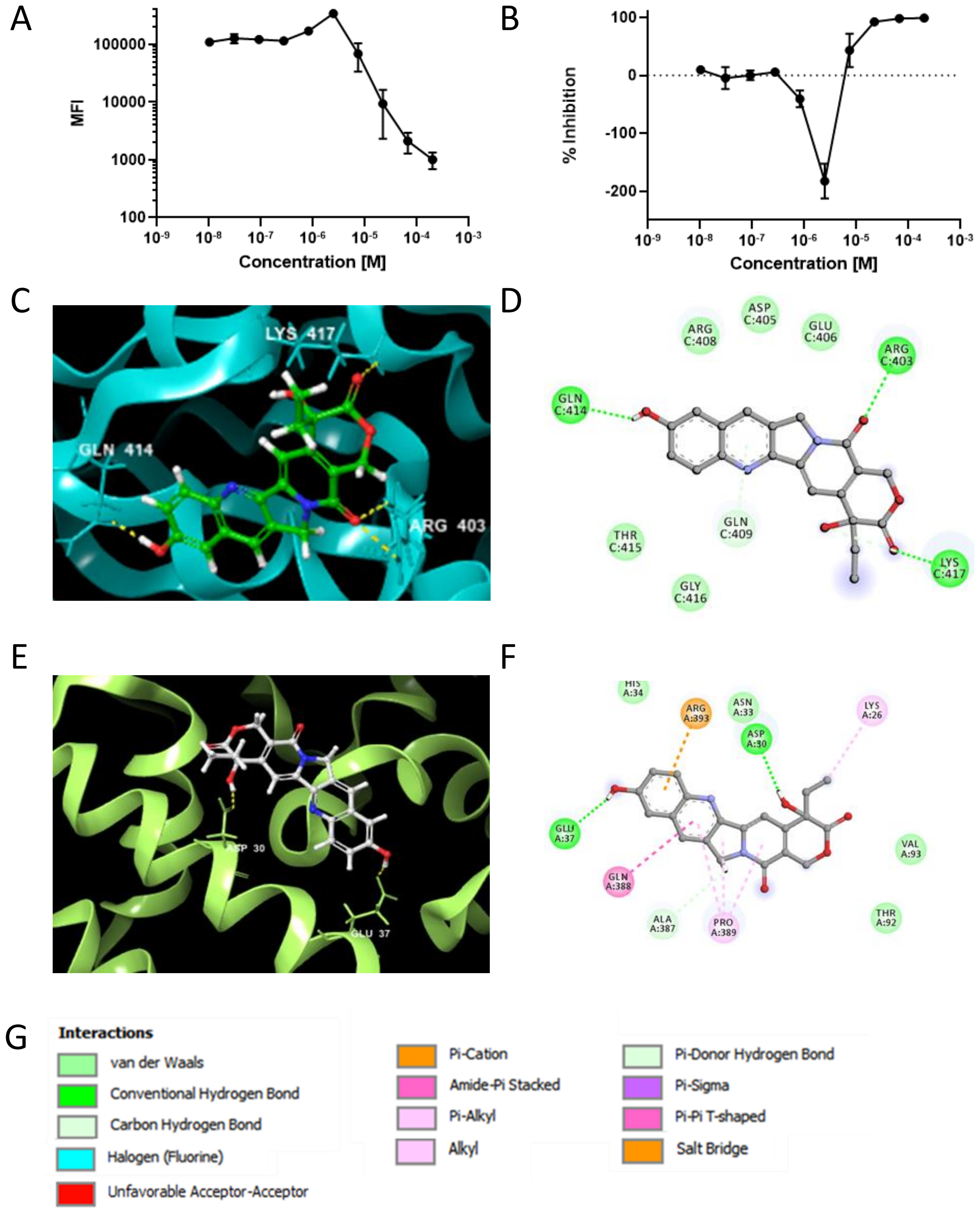


**Figure S5: EC50 and computational binding data for Hydroxy Camptothecine**. A,B) EC50 data are expressed as median fluorescent intensity (MFI, A) or percent inhibition (B). C) Three dimensional and D) two dimensional computationally determined lowest-energy docking poses of Hydroxy Camptothecine and the SARS-CoV-2 spike protein. E) Three dimensional and F) two dimensional computational docking of Hydroxy Camptothecine and human ACE2. G) Key showing color-codes for interaction types.

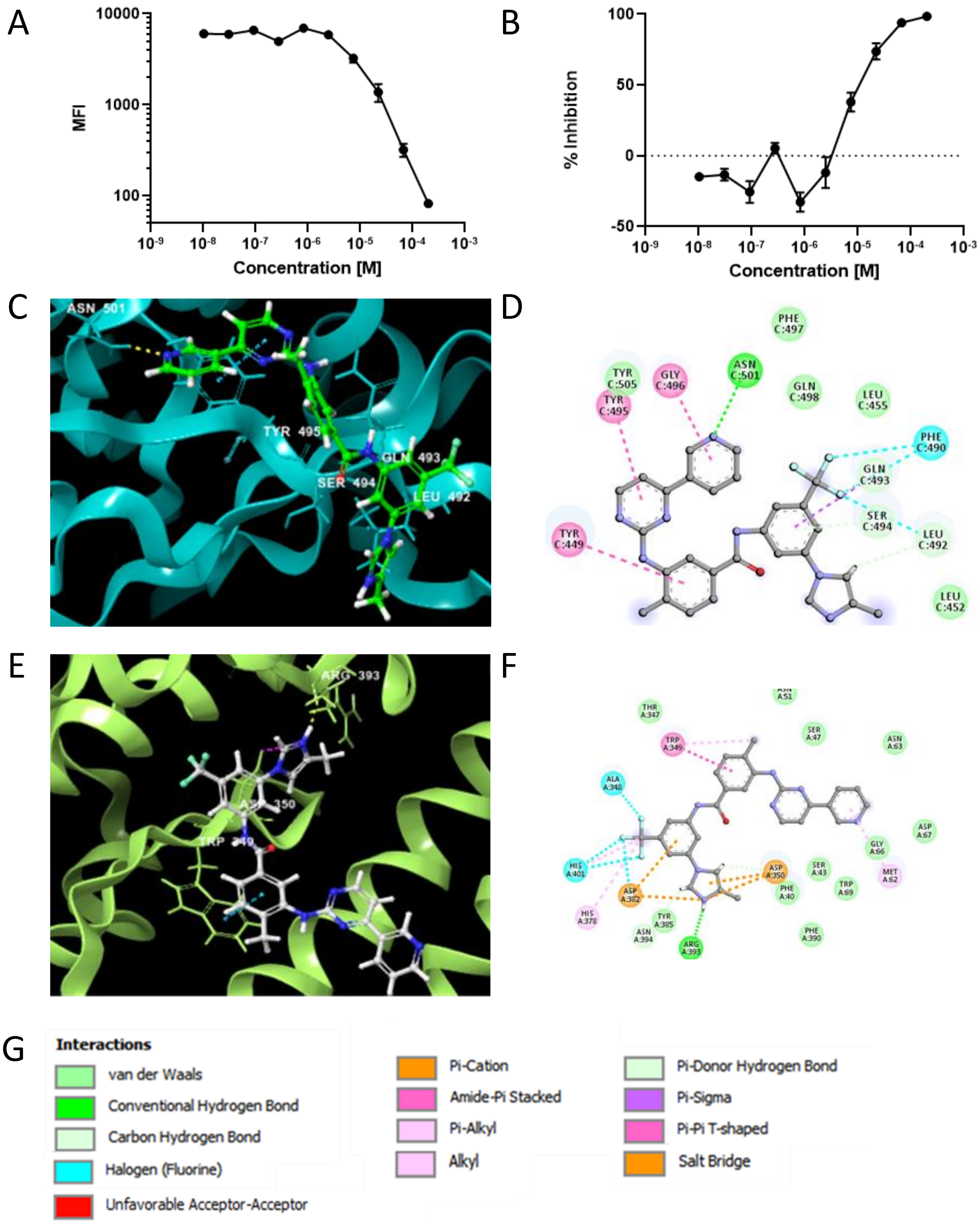


**Figure S6: EC50 and computational binding data for Nilotinib HCl**. A,B) EC50 data are expressed as median fluorescent intensity (MFI, A) or percent inhibition (B). C) Three dimensional and D) two dimensional computationally determined lowest-energy docking poses of Nilotinib HCl and the SARS-CoV-2 spike protein. E) Three dimensional and F) two dimensional computational docking of Nilotinib HCl and human ACE2. G) Key showing color-codes for interaction types.

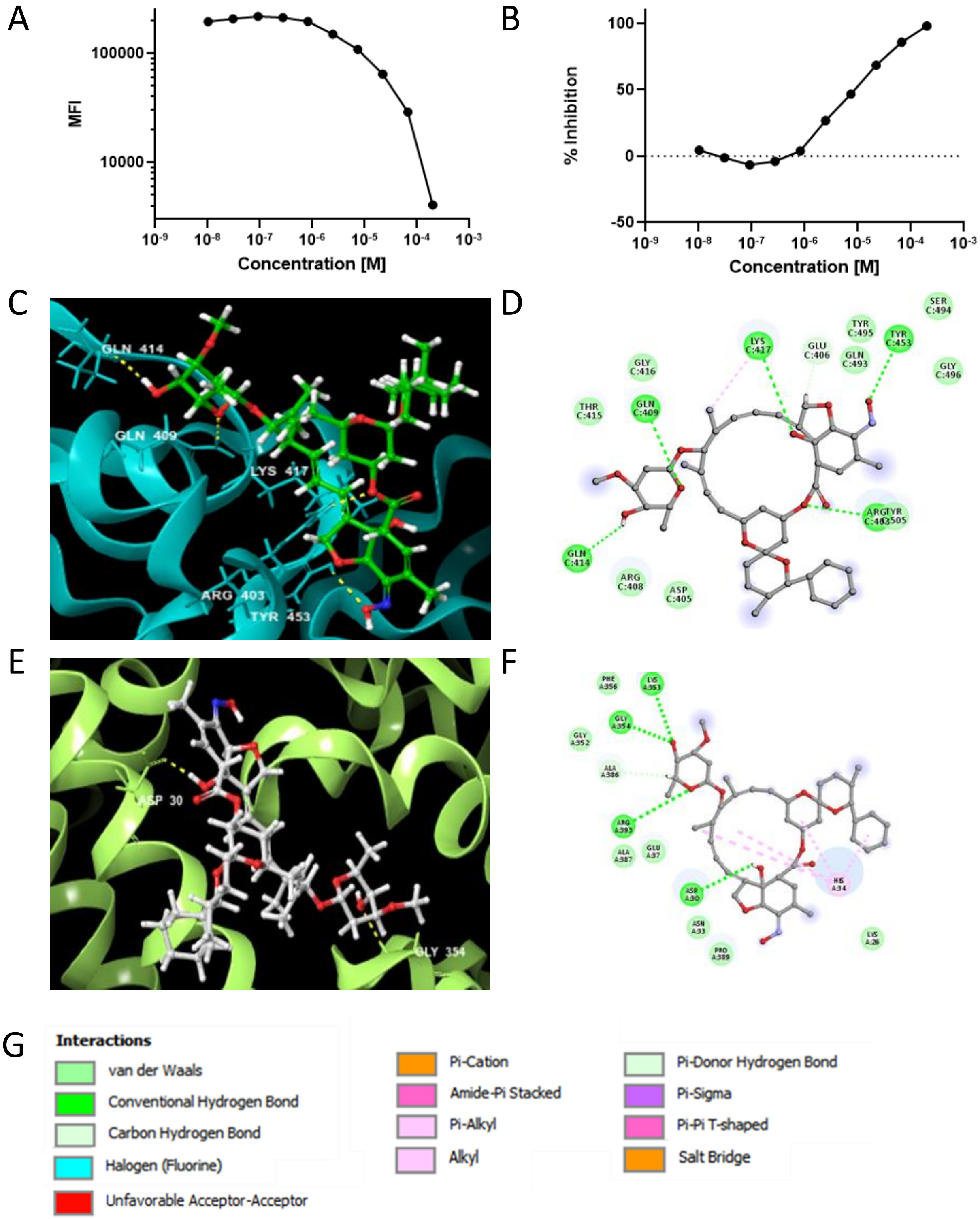


**Figure S7: EC50 and computational binding data for Selamectin**. A,B) EC50 data are expressed as median fluorescent intensity (MFI, A) or percent inhibition (B). C) Three dimensional and D) two dimensional computationally determined lowest-energy docking poses of Selamectin and the SARS-CoV-2 spike protein. E) Three dimensional and F) two dimensional computational docking of Selamectin and human ACE2. G) Key showing color-codes for interaction types.

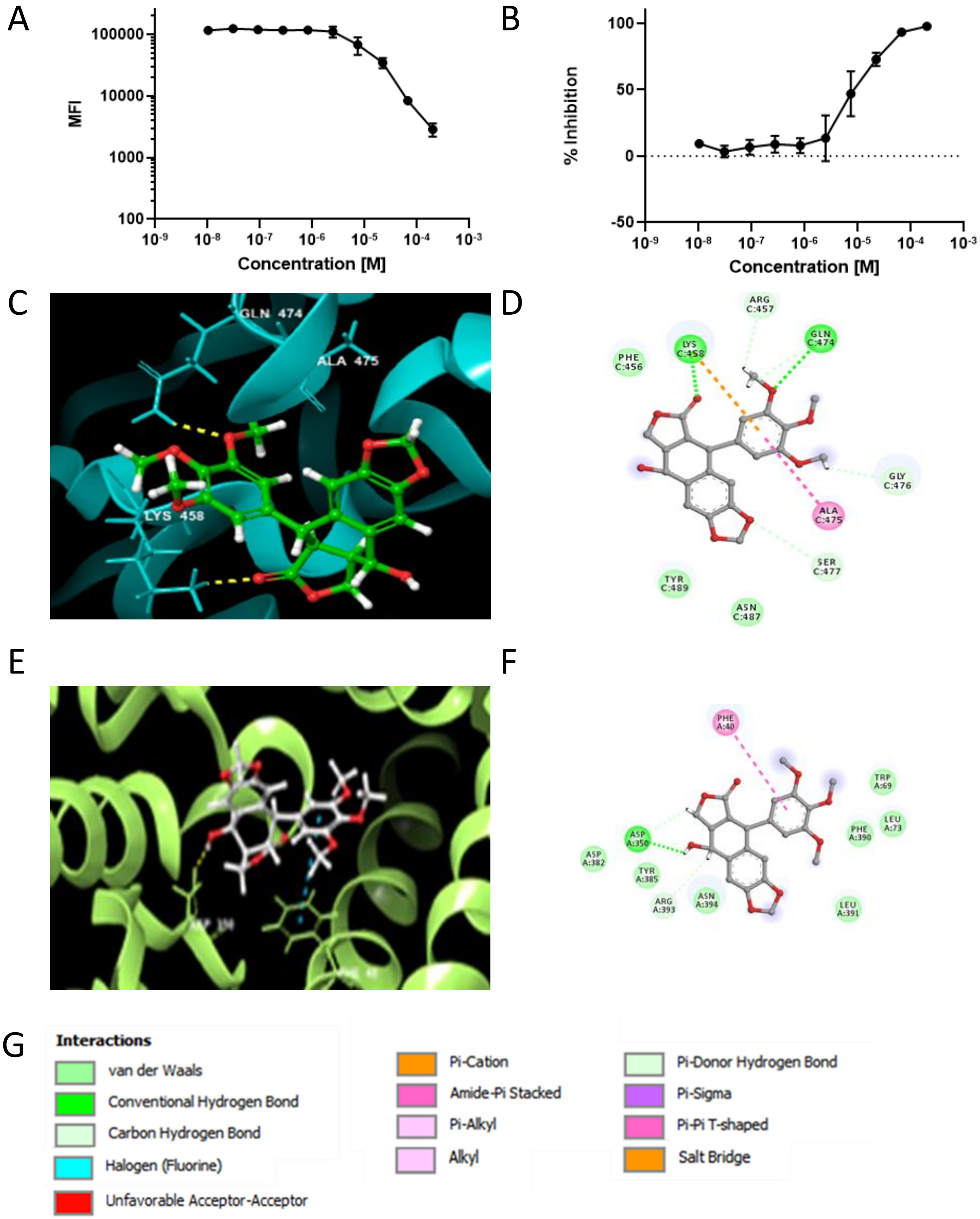


**Figure S8: EC50 and computational binding data for Picropodophyllin**. A,B) EC50 data are expressed as median fluorescent intensity (MFI, A) or percent inhibition (B). C Three dimensional and D) two dimensional computationally determined lowest-energy docking poses of Picropodophyllin and the SARS-CoV-2 spike protein. E) Three dimensional and F) two dimensional computational docking of Picropodophyllin and human ACE2. G) Key showing color-codes for interaction types.

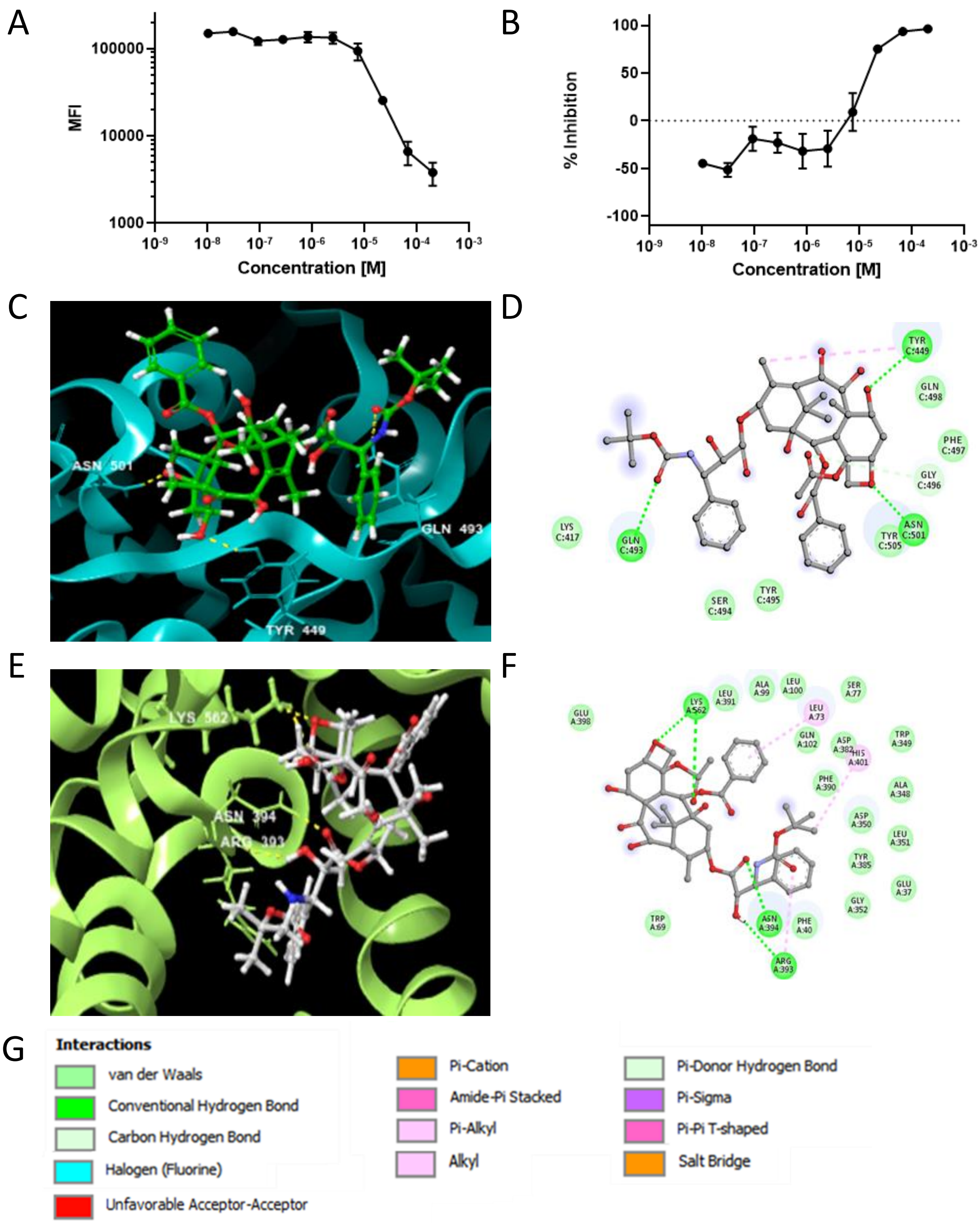


**Figure S9: EC50 and computational binding data for Docetaxel**. A,B) EC50 data are expressed as median fluorescent intensity (MFI, A) or percent inhibition (B). C) Three dimensional and D) two dimensional computationally determined lowest-energy docking poses of Docetaxel and the SARS-CoV-2 spike protein. E) Three dimensional and F) two dimensional computational docking of Docetaxel and human ACE2. G) Key showing color-codes for interaction types.

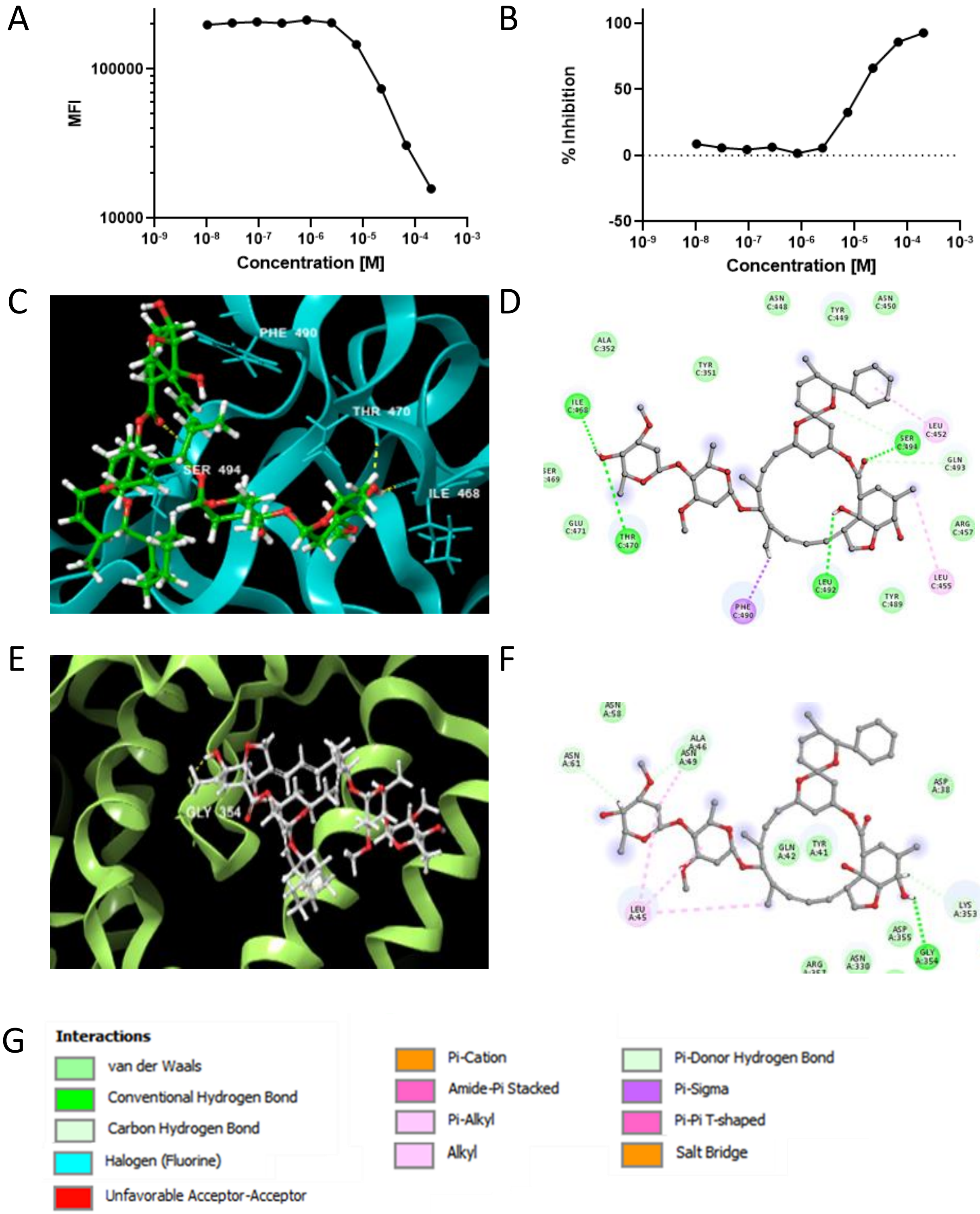


**Figure S10: EC50 and computational binding data for Doramectin**. A,B) EC50 data are expressed as median fluorescent intensity (MFI, A) or percent inhibition (B). C) Three dimensional and D) two dimensional computationally determined lowest-energy docking poses of Doramectin and the SARS-CoV-2 spike protein. E) Three dimensional and F) two dimensional computational docking of Doramectin and human ACE2. G) Key showing color-codes for interaction types.

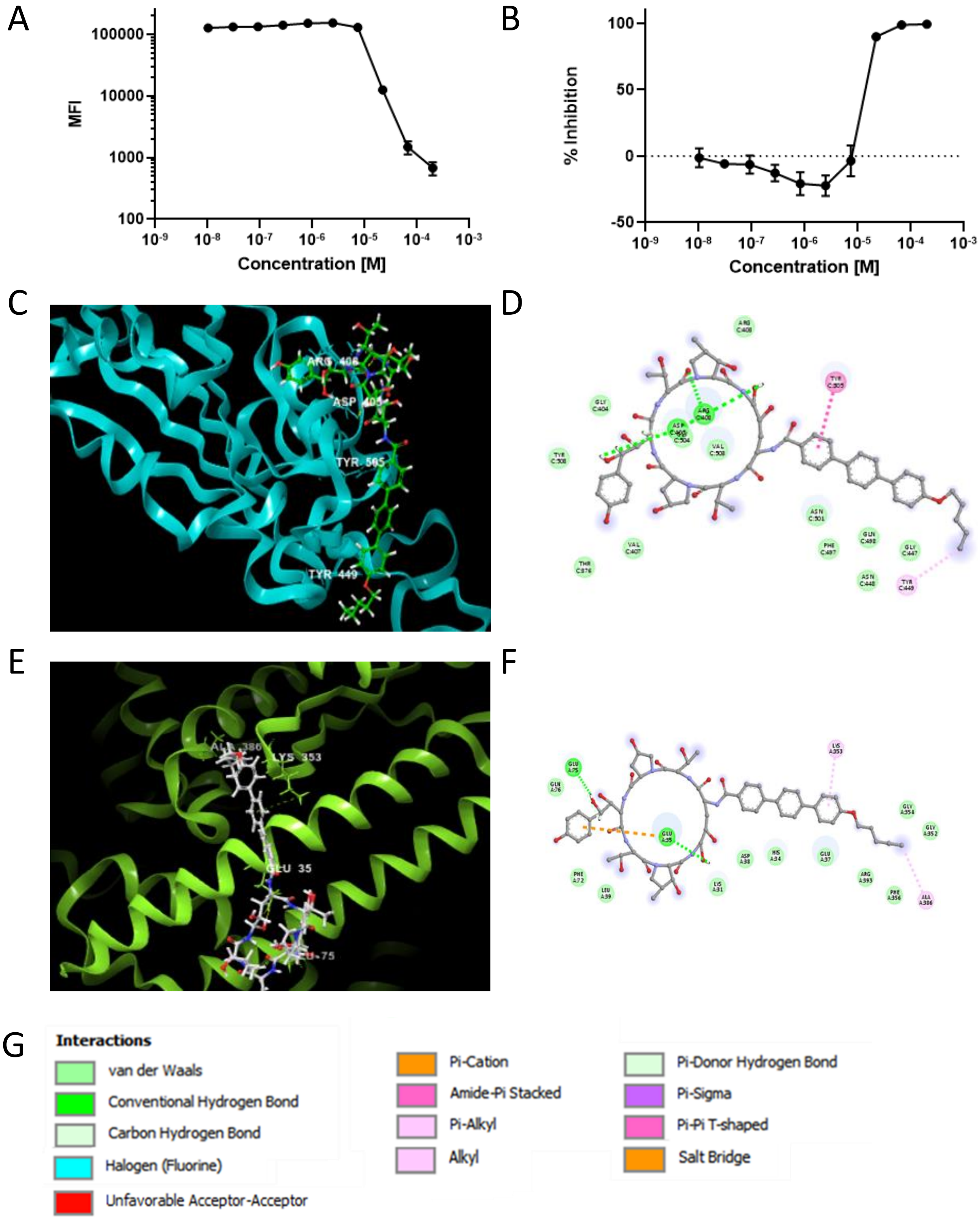


**Figure S11: EC50 and computational binding data for Anidulafungin**. A,B) EC50 data are expressed as median fluorescent intensity (MFI, A) or percent inhibition (B). C) Three dimensional and D) two dimensional computationally determined lowest-energy docking poses of Anidulafungin and the SARS-CoV-2 spike protein. E) Three dimensional and F) two dimensional computational docking of Anidulafungin and human ACE2. G) Key showing color-codes for interaction types.

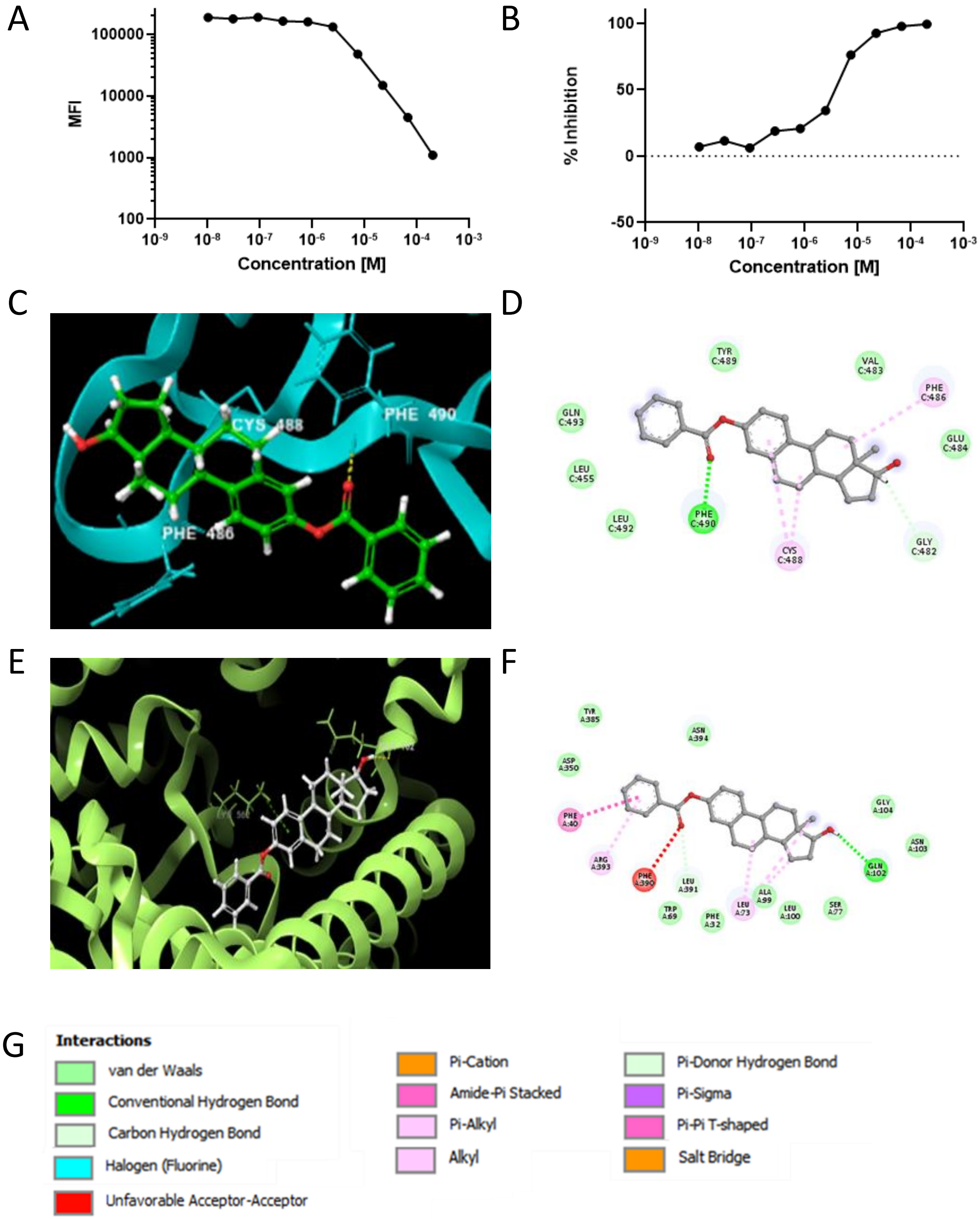


**Figure S12: EC50 and computational binding data for Estradiol Benzoate**.

A,B) EC50 data are expressed as median fluorescent intensity (MFI, A) or percent inhibition (B). C) Three dimensional and D) two dimensional computationally determined lowest-energy docking poses of Estradiol Benzoate and the SARS-CoV-2 spike protein. E) Three dimensional and F) two dimensional computational docking of Estradiol Benzoate and human ACE2. G) Key showing color-codes for interaction types.
