## Supplementary material for "A repurposed drug screen identifies compounds that inhibit the binding of the COVID-19 spike protein to ACE2": Suppl. Table 1

| Drug Name | Plate Number | Well Location | % Inhibition, Screen |
|---|---|---|---|
| Axitinib | 1 | 1 | 20 |
| Gefitinib (ZD1839) | 1 | 2 | 70 |
| Sorafenib Tosylate | 1 | 3 | 21 |
| Crizotinib (PF-02341066) | 1 | 4 | 55 |
| Docetaxel | 1 | 5 | 98 |
| Anastrozole | 1 | 6 | 25 |
| Cladribine | 1 | 7 | 23 |
| Methotrexate | 1 | 8 | -187 |
| Letrozole | 1 | 9 | 65 |
| Entecavir Hydrate | 1 | 10 | 48 |
| Roxadustat (FG-4592) | 1 | 11 | 19 |
| Imatinib Mesylate (STI571) | 1 | 12 | 0 |
| Sunitinib Malate | 1 | 13 | 34 |
| Vismodegib (GDC-0449) | 1 | 14 | 64 |
| Paclitaxel | 1 | 15 | 89 |
| Aprepitant | 1 | 16 | 94 |
| Decitabine | 1 | 17 | -79 |
| Bendamustine HCl | 1 | 18 | 19 |
| Temozolomide | 1 | 19 | -111 |
| Nepafenac | 1 | 20 | 24 |
| Nintedanib (BIBF 1120) | 1 | 21 | -43 |
| Lapatinib (GW-572016) Ditosylate | 1 | 22 | 88 |
| Temsirolimus (CCI-779, NSC 683864) | 1 | 23 | 96 |
| Belinostat (PXD101) | 1 | 24 | 46 |
| Capecitabine | 1 | 25 | 19 |
| Bicalutamide | 1 | 26 | 83 |
| Dutasteride | 1 | 27 | 68 |
| Epirubicin HCl | 1 | 28 | -59 |
| Tamoxifen | 1 | 29 | 30 |
| Rufinamide | 1 | 30 | 96 |
| Afatinib (BIBW2992) | 1 | 31 | -54 |
| Lenalidomide (CC-5013) | 1 | 32 | 19 |
| Vorinostat (SAHA, MK0683) | 1 | 33 | 38 |
| Rucaparib (AG-014699,PF-01367338) p | 1 | 34 | 14 |
| Lenvatinib (E7080) | 1 | 35 | 80 |
| Fulvestrant | 1 | 36 | 76 |
| Melatonin | 1 | 37 | 15 |
| Etoposide | 1 | 38 | -69 |
| Vincristine sulfate | 1 | 39 | 61 |
| Posaconazole | 1 | 40 | 97 |
| Bortezomib (PS-341) | 1 | 41 | 71 |
| Panobinostat (LBH589) | 1 | 42 | 41 |
| Entinostat (MS-275) | 1 | 43 | 26 |
| Cabozantinib (XL184, BMS-907351) | 1 | 44 | 79 |
| Valproic acid sodium salt (Sodium valp | 1 | 45 | 7 |
| Raltitrexed | 1 | 46 | 39 |

| | | | |
|---|---|---|---|
| Bisoprolol fumarate | 1 | 47 | -23 |
| Raloxifene HCl | 1 | 48 | 97 |
| Agomelatine | 1 | 49 | 35 |
| Prasugrel | 1 | 50 | -24 |
| Bosutinib (SKI-606) | 1 | 51 | 85 |
| Nilotinib (AMN-107) | 1 | 52 | 99 |
| Enzastaurin (LY317615) | 1 | 53 | -12 |
| Everolimus (RAD001) | 1 | 54 | 94 |
| Regorafenib (BAY 73-4506) | 1 | 55 | 24 |
| Thalidomide | 1 | 56 | 40 |
| Tivozanib (AV-951) | 1 | 57 | 86 |
| Fludarabine Phosphate | 1 | 58 | 12 |
| Leflunomide | 1 | 59 | 83 |
| Ramelteon | 1 | 60 | 46 |
| Dasatinib | 1 | 61 | 97 |
| Pazopanib HCl (GW786034 HCl) | 1 | 62 | 88 |
| Olaparib (AZD2281, Ku-0059436) | 1 | 63 | 25 |
| Malotilate | 1 | 64 | 65 |
| Danoprevir (ITMN-191) | 1 | 65 | -31 |
| Exemestane | 1 | 66 | 70 |
| Doxorubicin (Adriamycin) HCl | 1 | 67 | -5 |
| Topotecan HCl | 1 | 68 | 15 |
| Enzalutamide (MDV3100) | 1 | 69 | 95 |
| Cinacalcet HCl | 1 | 70 | 35 |
| Ridaforolimus (Deforolimus, MK-8669 | 1 | 71 | 51 |
| Rapamycin (Sirolimus) | 1 | 72 | 97 |
| Masitinib (AB1010) | 1 | 73 | 71 |
| Ivacaftor (VX-770) | 1 | 74 | 53 |
| Ritonavir | 1 | 75 | 94 |
| Finasteride | 1 | 76 | 68 |
| Fluorouracil (5-Fluoracil, 5-FU) | 1 | 77 | 8 |
| 2-Methoxyestradiol (2-MeOE2) | 1 | 78 | 82 |
| Dienogest | 1 | 79 | 53 |
| Celecoxib | 1 | 80 | 93 |
| Vemurafenib | 2 | 1 | 45 |
| Benazepril HCl | 2 | 2 | 17 |
| Tegafur | 2 | 3 | -80 |
| Edaravone | 2 | 4 | -15 |
| Fluvoxamine Maleate | 2 | 5 | 70 |
| Loratadine | 2 | 6 | 83 |
| Ruxolitinib | 2 | 7 | 22 |
| Omeprazole | 2 | 8 | 10 |
| Tenofovir Disoproxil Fumarate | 2 | 9 | 63 |
| Clopidogrel | 2 | 10 | 82 |
| Acarbose | 2 | 11 | -44 |
| Budesonide | 2 | 12 | 80 |
| Ifosfamide | 2 | 13 | 8 |

| | | | |
|---|---|---|---|
| Etodolac | 2 | 14 | -19 |
| Gatifloxacin | 2 | 15 | 11 |
| Losartan Potassium | 2 | 16 | -11 |
| Isotretinoin | 2 | 17 | -27 |
| Ondansetron HCl | 2 | 18 | 79 |
| Tenofovir | 2 | 19 | -62 |
| Ranolazine 2HCl | 2 | 20 | -1 |
| Adapalene | 2 | 21 | 70 |
| Bumetanide | 2 | 22 | 8 |
| Megestrol Acetate | 2 | 23 | 51 |
| Etomidate | 2 | 24 | 33 |
| Genistein | 2 | 25 | 37 |
| Amonafide | 2 | 26 | -32 |
| Lopinavir | 2 | 27 | 92 |
| Oxcarbazepine | 2 | 28 | 9 |
| Tigecycline | 2 | 29 | 15 |
| Repaglinide | 2 | 30 | 42 |
| Altretamine | 2 | 31 | 38 |
| Carmofur | 2 | 32 | -23 |
| Mercaptopurine | 2 | 33 | -19 |
| Felbamate | 2 | 34 | -3 |
| Glimepiride | 2 | 35 | 36 |
| Acitretin | 2 | 36 | -59 |
| Meropenem | 2 | 37 | -12 |
| Pirarubicin | 2 | 38 | 78 |
| Trilostane | 2 | 39 | 1 |
| Rolipram | 2 | 40 | -20 |
| Amisulpride | 2 | 41 | -59 |
| Cetirizine DiHCl | 2 | 42 | 94 |
| Streptozotocin | 2 | 43 | -58 |
| Fluconazole | 2 | 44 | -3 |
| Ivermectin | 2 | 45 | 89 |
| Daptomycin | 2 | 46 | 25 |
| Mianserin HCl | 2 | 47 | 75 |
| Pizotifen Malate | 2 | 48 | 32 |
| Vecuronium Bromide | 2 | 49 | -79 |
| Sildenafil Citrate | 2 | 50 | 84 |
| Aniracetam | 2 | 51 | 5 |
| Cilnidipine | 2 | 52 | 33 |
| Costunolide | 2 | 53 | 41 |
| Flumazenil | 2 | 54 | -40 |
| Lansoprazole | 2 | 55 | 81 |
| Doripenem Hydrate | 2 | 56 | -61 |
| Mosapride Citrate | 2 | 57 | 72 |
| Resveratrol | 2 | 58 | -17 |
| Bimatoprost | 2 | 59 | 57 |
| Sumatriptan Succinate | 2 | 60 | -62 |

| | | | |
|---|---|---|---|
| Artemisinin | 2 | 61 | 34 |
| Cilostazol | 2 | 62 | 79 |
| Dexamethasone | 2 | 63 | 74 |
| Fluoxetine HCl | 2 | 64 | 57 |
| Levetiracetam | 2 | 65 | 7 |
| Gestodene | 2 | 66 | 43 |
| Nafamostat Mesylate | 2 | 67 | -56 |
| Rocuronium Bromide | 2 | 68 | 25 |
| Linezolid | 2 | 69 | -31 |
| Tamsulosin | 2 | 70 | -109 |
| Asenaprine Maleate | 2 | 71 | 56 |
| Floxuridine | 2 | 72 | -21 |
| Doxazosin Mesylate | 2 | 73 | 59 |
| Flupirtine Maleate | 2 | 74 | 52 |
| Lidocaine | 2 | 75 | 52 |
| Drospirenone | 2 | 76 | 21 |
| Naftopidil DiHCl | 2 | 77 | 25 |
| Stavudine | 2 | 78 | -22 |
| Alfuzosin HCl | 2 | 79 | -46 |
| Tianeptine Sodium | 2 | 80 | 30 |
| Tizanidine HCl | 3 | 1 | -23 |
| Tipifarnib | 3 | 2 | 98 |
| Daclatasvir | 3 | 3 | 99 |
| Betamethasone | 3 | 4 | -16 |
| Natamycin | 3 | 5 | -76 |
| Pomalidomide | 3 | 6 | 17 |
| Olmesartan Medoxomil | 3 | 7 | -40 |
| Silodosin | 3 | 8 | -63 |
| Naproxen Sodium | 3 | 9 | -8 |
| Amphotericin B | 3 | 10 | -103 |
| Topiramate | 3 | 11 | -4 |
| Atazanavir Sulfate | 3 | 12 | 93 |
| Iloperidone | 3 | 13 | -58 |
| Mycophenolate Mofetil | 3 | 14 | 7 |
| Telaprevir | 3 | 15 | 29 |
| Tazarotene | 3 | 16 | 2 |
| Cefdinir | 3 | 17 | -58 |
| Riluzole | 3 | 18 | 23 |
| Nitazoxanide | 3 | 19 | -33 |
| Ibuprofen | 3 | 20 | -31 |
| Tranilast | 3 | 21 | -1 |
| VX-745 | 3 | 22 | -4 |
| HMN-214 | 3 | 23 | 45 |
| Dyphylline | 3 | 24 | 5 |
| Saxagliptin | 3 | 25 | -1 |
| Fasudil HCl | 3 | 26 | -66 |
| Clotrimazole | 3 | 27 | 39 |

| | | | |
|---|---|---|---|
| Sulfameter | 3 | 28 | -5 |
| Triamcinolone Acetonide | 3 | 29 | 14 |
| Amprenavir | 3 | 30 | 14 |
| Venlafaxine HCl | 3 | 31 | -22 |
| Moxifloxacin HCl | 3 | 32 | -9 |
| Naratriptan HCl | 3 | 33 | -46 |
| Aztreonam | 3 | 34 | 27 |
| Febuxostat | 3 | 35 | -29 |
| Sulfasalazine | 3 | 36 | -23 |
| Rizatriptan Benzoate | 3 | 37 | -39 |
| Prilocaine | 3 | 38 | -11 |
| Orlistat | 3 | 39 | 17 |
| Albendazole | 3 | 40 | 18 |
| Voriconazole | 3 | 41 | 43 |
| Calcitriol | 3 | 42 | 59 |
| Ponatinib | 3 | 43 | 67 |
| Alprostadil | 3 | 44 | 16 |
| Dapagliflozin | 3 | 45 | 37 |
| Candesartan | 3 | 46 | 2 |
| Pyridostigmine Bromide | 3 | 47 | 10 |
| Darunavir Ethanolate | 3 | 48 | 22 |
| Allopurinol | 3 | 49 | 5 |
| Chlorothiazide | 3 | 50 | -7 |
| Zileuton | 3 | 51 | 58 |
| Doxercalciferol | 3 | 52 | 17 |
| Fludarabine | 3 | 53 | 18 |
| Lactulose | 3 | 54 | 53 |
| Nebivolol HCl | 3 | 55 | -9 |
| Apixaban | 3 | 56 | 69 |
| Methimazole | 3 | 57 | 10 |
| Prednisone | 3 | 58 | 2 |
| Allopurinol Sodium | 3 | 59 | 19 |
| Ursodiol | 3 | 60 | -37 |
| Ziprasidone HCl | 3 | 61 | 82 |
| Alfacalcidol | 3 | 62 | 55 |
| Pralatrexate | 3 | 63 | 46 |
| Tadalafil | 3 | 64 | 96 |
| Pimobendan | 3 | 65 | 92 |
| Reserpine | 3 | 66 | 87 |
| Metolazone | 3 | 67 | 27 |
| Acetylcysteine | 3 | 68 | 2 |
| Zafirlukast | 3 | 69 | -10 |
| Nitrofural | 3 | 70 | -57 |
| Zonisamide | 3 | 71 | 62 |
| Safinamide Mesylate | 3 | 72 | 22 |
| Cefaclor | 3 | 73 | 54 |
| Cyclosporine | 3 | 74 | 63 |

| | | | |
|---|---|---|---|
| VX-809 (Lumacaftor) | 3 | 75 | 33 |
| Furosemide | 3 | 76 | 17 |
| Cefoperazone | 3 | 77 | 1 |
| Ethinyl Estradiol | 3 | 78 | 39 |
| Erythromycin | 3 | 79 | 14 |
| Ketoprofen | 3 | 80 | 13 |
| Ketorolac | 4 | 1 | -69 |
| Enalaprilat Dihydrate | 4 | 2 | -57 |
| Aminoglutethimide | 4 | 3 | -105 |
| Ipratropium Bromide | 4 | 4 | -97 |
| Hydrocortisone | 4 | 5 | -104 |
| Deferasirox | 4 | 6 | -211 |
| Azathioprine | 4 | 7 | -79 |
| Meloxicam | 4 | 8 | 1 |
| Nevirapine | 4 | 9 | -103 |
| Pitavastatin Calcium | 4 | 10 | -115 |
| Adenosine | 4 | 11 | -77 |
| Dofetilide | 4 | 12 | -225 |
| Aminophylline | 4 | 13 | -142 |
| Sulfanilamide | 4 | 14 | -72 |
| Desonide | 4 | 15 | 32 |
| Piroxicam | 4 | 16 | -16 |
| Indomethacin | 4 | 17 | -173 |
| Mesna | 4 | 18 | -155 |
| NEXIUM (esomeprazole magnesium) | 4 | 19 | 34 |
| Rifapentine | 4 | 20 | -71 |
| Zolmitriptan | 4 | 21 | -78 |
| Isradipine | 4 | 22 | 80 |
| Lubiprostone | 4 | 23 | 43 |
| Betamethasone Dipropionate | 4 | 24 | 86 |
| Didanosine | 4 | 25 | -109 |
| Gemcitabine | 4 | 26 | -114 |
| Terbinafine | 4 | 27 | -34 |
| Methocarbamol | 4 | 28 | -98 |
| Nicotinic Acid | 4 | 29 | -10 |
| Suprofen | 4 | 30 | 23 |
| Telbivudine | 4 | 31 | 2 |
| Estrone | 4 | 32 | 76 |
| Amorolfine HCl | 4 | 33 | -109 |
| Meprednisone | 4 | 34 | -24 |
| Divalproex Sodium | 4 | 35 | -89 |
| Glipizide | 4 | 36 | -209 |
| Levonorgestrel | 4 | 37 | 30 |
| Prednisolone | 4 | 38 | 14 |
| Nimodipine | 4 | 39 | 96 |
| Pyrazinamide | 4 | 40 | 54 |
| Monobenzone | 4 | 41 | 52 |

| | | | |
|---|---|---|---|
| Flucytosine | 4 | 42 | -34 |
| Chloramphenicol | 4 | 43 | -77 |
| Betamethasone Valerate | 4 | 44 | 92 |
| Emtricitabine | 4 | 45 | -34 |
| Glyburide (Glibenclamide) | 4 | 46 | -71 |
| Gemfibrozil | 4 | 47 | 0 |
| Telmisartan | 4 | 48 | 68 |
| Nisoldipine | 4 | 49 | 72 |
| Quetiapine Fumarate | 4 | 50 | 65 |
| Tretinoin | 4 | 51 | -390 |
| Trichlormethiazide | 4 | 52 | -15 |
| Flurbiprofen | 4 | 53 | -52 |
| Praziquantel | 4 | 54 | -53 |
| Progesterone | 4 | 55 | 48 |
| Fomepizole | 4 | 56 | -50 |
| Indapamide | 4 | 57 | -63 |
| Thiabendazole | 4 | 58 | -55 |
| Octocrylene | 4 | 59 | 24 |
| Rifampin | 4 | 60 | -65 |
| Phenylbutazone | 4 | 61 | 85 |
| Loteprednol etabonate | 4 | 62 | 50 |
| Disulfiram | 4 | 63 | 13 |
| Busulfan | 4 | 64 | -32 |
| Lamivudine | 4 | 65 | -14 |
| Adefovir Dipivoxil | 4 | 66 | 92 |
| Mitotane | 4 | 67 | 53 |
| Guaifenesin | 4 | 68 | 25 |
| Oxybutynin | 4 | 69 | 86 |
| Cefditoren Pivoxil | 4 | 70 | 96 |
| Ezetimibe | 4 | 71 | 74 |
| Mesalamine | 4 | 72 | -66 |
| Carbamazepine | 4 | 73 | -81 |
| Estradiol | 4 | 74 | -130 |
| Zalcitabine | 4 | 75 | -64 |
| Methylprednisolone | 4 | 76 | 25 |
| Rifabutin | 4 | 77 | -2 |
| Enoxacin | 4 | 78 | -61 |
| Sulfadiazine | 4 | 79 | -68 |
| Chlorprothixine | 5 | 1 | -69 |
| Vidarabine | 5 | 2 | -25 |
| Ranolazine | 5 | 3 | 0 |
| Fenoprofen Calcium | 5 | 4 | -24 |
| Butoconazole Nitrate | 5 | 5 | 64 |
| Curcumin | 5 | 6 | 23 |
| Diphenhydramine HCl | 5 | 7 | -66 |
| Felodipine | 5 | 8 | 46 |
| Tropisetron HCl | 5 | 9 | -99 |

| | | | |
|---|---|---|---|
| Fluvastatin Sodium | 5 | 10 | -20 |
| Oxytetracycline | 5 | 11 | -59 |
| Verteporfin | 5 | 12 | -181 |
| Ranitidine Hydrochloride | 5 | 13 | -69 |
| Erdosteine | 5 | 14 | -11 |
| Azithromycin | 5 | 15 | 0 |
| Daidzein | 5 | 16 | 11 |
| Dapoxetine HCl | 5 | 17 | -174 |
| Deflazacort | 5 | 18 | 16 |
| Nicotinamide | 5 | 19 | -14 |
| Tioconazole | 5 | 20 | 65 |
| Thioguanine | 5 | 21 | -29 |
| Teniposide | 5 | 22 | 97 |
| Acipimox | 5 | 23 | -9 |
| Betaxolol HCl | 5 | 24 | -34 |
| Albendazole Oxide | 5 | 25 | 1 |
| Bifonazole | 5 | 26 | -1 |
| Valaciclovir HCl | 5 | 27 | 1 |
| Nizatidine | 5 | 28 | -25 |
| Vitamin B12 | 5 | 29 | -4 |
| Tropicamide | 5 | 30 | -29 |
| Thiotepa | 5 | 31 | 54 |
| Tetrabenazine | 5 | 32 | 37 |
| Aciclovir | 5 | 33 | -17 |
| Proparacaine HCl | 5 | 34 | -16 |
| Chloroxine | 5 | 35 | -62 |
| Perfloxacin Mesylate | 5 | 36 | -14 |
| Ganciclovir | 5 | 37 | -3 |
| Carbidopa | 5 | 38 | -31 |
| Diclofenac Sodium | 5 | 39 | -26 |
| Pregnenolone | 5 | 40 | -73 |
| Toremifene Citrate | 5 | 41 | 61 |
| Rifaximin | 5 | 42 | 33 |
| Nifedipine | 5 | 43 | 69 |
| Pranlukast | 5 | 44 | -197 |
| Lomustine | 5 | 45 | 10 |
| Metoprolol Tartrate | 5 | 46 | -11 |
| Roxatidine Acetate HCl | 5 | 47 | -56 |
| Valsartan | 5 | 48 | -22 |
| Avobenzone | 5 | 49 | 12 |
| Sulfamethoxazole | 5 | 50 | -25 |
| Ethionamide | 5 | 51 | -7 |
| Simvastatin | 5 | 52 | 89 |
| Amiloride HCl | 5 | 53 | -105 |
| Oxfendazole | 5 | 54 | 96 |
| Chenodeoxycholic Acid | 5 | 55 | 12 |
| Dienestrol | 5 | 56 | 52 |

| | | | |
|---|---|---|---|
| Protionamide | 5 | 57 | 16 |
| Dipyridamole | 5 | 58 | 79 |
| Amlodipine | 5 | 59 | 70 |
| Sulfisoxazole | 5 | 60 | -43 |
| Trifluridine | 5 | 61 | -6 |
| Ramipril | 5 | 62 | -11 |
| Amlodipine Besylate | 5 | 63 | 63 |
| Carvedilol | 5 | 64 | 87 |
| Cimetidine | 5 | 65 | -44 |
| Diethylstilbestrol | 5 | 66 | 95 |
| Idoxuridine | 5 | 67 | 3 |
| Hydroxyurea | 5 | 68 | -12 |
| Metronidazole | 5 | 69 | -5 |
| Crystal Violet | 5 | 70 | 98 |
| Azacitidine | 5 | 71 | 14 |
| Fenofibrate | 5 | 72 | -69 |
| Chlorpeniramine Maleate | 5 | 73 | -57 |
| Atracurium Besylate | 5 | 74 | -35 |
| Clemastine Fumarate | 5 | 75 | 70 |
| Diltiazem HCl | 5 | 76 | 30 |
| Sparfloxacin | 5 | 77 | -22 |
| Potassium Iodide | 5 | 78 | 12 |
| Flutamide | 5 | 79 | 2 |
| Haloperidol | 5 | 80 | -77 |
| Phenindione | 6 | 1 | 7 |
| Methoxsalen | 6 | 2 | 5 |
| Nefiracetam | 6 | 3 | -35 |
| Mometasone furoate | 6 | 4 | 98 |
| Pyrimethamine | 6 | 5 | -17 |
| Nelarabine | 6 | 6 | -8 |
| Aspartame | 6 | 7 | -27 |
| Doxifluridine | 6 | 8 | 9 |
| Tolnaftate | 6 | 9 | 10 |
| Ozagrel HCl | 6 | 10 | -19 |
| Triamcinolone | 6 | 11 | -6 |
| Miconazole Nitrate | 6 | 12 | 69 |
| Nicorandil | 6 | 13 | -67 |
| Propylthiouracil | 6 | 14 | 3 |
| Sulindac | 6 | 15 | 22 |
| Ketotifen Fumarate | 6 | 16 | -20 |
| Candesartan Cilexetil | 6 | 17 | 9 |
| Pioglitazone HCl | 6 | 18 | -126 |
| Terazosin HCl Dihydrate | 6 | 19 | 23 |
| Argatroban | 6 | 20 | 43 |
| Nystatin (Fungicidin) | 6 | 21 | -71 |
| Sulfamethizole | 6 | 22 | -9 |
| Tamoxifen Citrate | 6 | 23 | 31 |

| | | | |
|---|---|---|---|
| Capsaicin(Vanilloid) | 6 | 24 | 83 |
| Pramipexole 2HCl Monohydrate | 6 | 25 | -35 |
| Urapidil HCl | 6 | 26 | -37 |
| Phentolamine Mesylate | 6 | 27 | 9 |
| Captopril | 6 | 28 | -11 |
| Bromhexine HCl | 6 | 29 | -4 |
| Prulifloxacin (NM441, AF 3013) | 6 | 30 | 28 |
| Isoniazid | 6 | 31 | -65 |
| Sulbactam | 6 | 32 | -76 |
| Meglumine | 6 | 33 | -37 |
| Fluticasone propionate | 6 | 34 | -9 |
| Suplatast Tosylate | 6 | 35 | -27 |
| Diclazuril | 6 | 36 | -49 |
| Nimesulide | 6 | 37 | -13 |
| Oxytetracycline Dihydrate | 6 | 38 | -43 |
| Lovastatin | 6 | 39 | 87 |
| Rosiglitazone HCl | 6 | 40 | 34 |
| Levofloxacin | 6 | 41 | 14 |
| Tolfenamic Acid | 6 | 42 | -2 |
| Aripiprazole | 6 | 43 | 22 |
| Lacidipine | 6 | 44 | 24 |
| Mirtazapine | 6 | 45 | 47 |
| Uridine | 6 | 46 | -14 |
| Dyclonine HCl | 6 | 47 | 74 |
| Cytidine | 6 | 48 | -4 |
| Tiopronin | 6 | 49 | -51 |
| Atorvastatin Calcium | 6 | 50 | -87 |
| Enalapril Maleate | 6 | 51 | 6 |
| Pranoprofen | 6 | 52 | 16 |
| Methscopolamine | 6 | 53 | 25 |
| Elvitegravir (GS-9137, JTK-303) | 6 | 54 | 64 |
| Benidipine HCl | 6 | 55 | 47 |
| Flunarizine 2HCl | 6 | 56 | 64 |
| Cyproterone Acetate | 6 | 57 | 62 |
| Orphenadrine Citrate | 6 | 58 | 25 |
| Balofloxacin | 6 | 59 | 41 |
| Famotidine | 6 | 60 | 4 |
| Menadione | 6 | 61 | -26 |
| Rimantadine | 6 | 62 | -44 |
| Amiodarone HCl | 6 | 63 | -18 |
| Maraviroc | 6 | 64 | 2 |
| Formoterol Hemifumarate | 6 | 65 | -36 |
| Fenticonazole Nitrate | 6 | 66 | 33 |
| Memantine HCl | 6 | 67 | 21 |
| Gimeracil | 6 | 68 | -41 |
| Lafutidine | 6 | 69 | -13 |
| Moexipril HCl | 6 | 70 | -10 |

| | | | |
|---|---|---|---|
| Metformin HCl | 6 | 71 | 5 |
| Primidone | 6 | 72 | -5 |
| Adenine HCl | 6 | 73 | -25 |
| Raltegravir (MK-0518) | 6 | 74 | -12 |
| Chlormezanone | 6 | 75 | -22 |
| Rebamipide | 6 | 76 | -5 |
| Cyproheptadine HCl | 6 | 77 | 51 |
| Cyclophosphamide Monohydrate | 6 | 78 | -6 |
| Moxonidine | 6 | 79 | 2 |
| Clevidipine Butyrate | 6 | 80 | 78 |
| Procaterol HCl | 7 | 1 | -24 |
| Almotriptan Malate | 7 | 2 | -283 |
| Pantoprazole | 7 | 3 | 22 |
| Probucol | 7 | 4 | -83 |
| Atropine Sulfate Monohydrate | 7 | 5 | -88 |
| Dichlorphenamide | 7 | 6 | -34 |
| Eltrombopag Olamine | 7 | 7 | 3 |
| Arbutin | 7 | 8 | -73 |
| Chlorogenic Acid | 7 | 9 | -1 |
| Emodin | 7 | 10 | 47 |
| Duloxetine HCl | 7 | 11 | -21 |
| Ambrisentan | 7 | 12 | -59 |
| Flunixin Meglumin | 7 | 13 | -44 |
| Arbidol HCl | 7 | 14 | 95 |
| Roflumilast | 7 | 15 | -127 |
| Ixazomib | 7 | 16 | -52 |
| Esomeprazole Sodium | 7 | 17 | 19 |
| Artemether | 7 | 18 | 9 |
| Cinchonidine | 7 | 19 | -81 |
| Enoxolone | 7 | 20 | -42 |
| Trimebutine | 7 | 21 | 65 |
| Bexarotene | 7 | 22 | -225 |
| Imidapril HCl | 7 | 23 | 28 |
| Dextrose | 7 | 24 | 1 |
| Sonidegib | 7 | 25 | 41 |
| Ixazomib Citrate | 7 | 26 | 0 |
| Fesoterodine Fumarate | 7 | 27 | 3 |
| Artesunate | 7 | 28 | -16 |
| Cinchonidine (LA 40221) | 7 | 29 | -252 |
| Formononetin | 7 | 30 | -54 |
| Ivabradine HCl | 7 | 31 | -98 |
| Temocapril HCl | 7 | 32 | 20 |
| Lapatinib | 7 | 33 | 33 |
| Xylose | 7 | 34 | -25 |
| Dabigatran Etexilate | 7 | 35 | 57 |
| Aliskiren Hemifumarate | 7 | 36 | 43 |
| 4-Methylumbelliferone | 7 | 37 | -4 |

| | | | |
|---|---|---|---|
| Baicalein | 7 | 38 | -81 |
| Colchicine | 7 | 39 | 3 |
| Ferulic Acid | 7 | 40 | -3 |
| Rivastigmine Tartrate | 7 | 41 | -31 |
| Gabexate Mesylate | 7 | 42 | -83 |
| Cisatracurium Besylate | 7 | 43 | -144 |
| Mestranol | 7 | 44 | 46 |
| Tebipenem Pivoxil | 7 | 45 | 67 |
| Formestane | 7 | 46 | -36 |
| Esculin | 7 | 47 | -7 |
| Baicalin | 7 | 48 | -61 |
| Cytisine | 7 | 49 | -19 |
| Glycyrrhizic Acid | 7 | 50 | -38 |
| Dexmedetomidine HCl | 7 | 51 | -72 |
| Rasagiline Mesylate | 7 | 52 | -6 |
| Dronedarone HCl | 7 | 53 | 69 |
| Naftopidil | 7 | 54 | 71 |
| Bazedoxifene Acetate | 7 | 55 | 78 |
| Irinotecan HCl Trihydrate | 7 | 56 | -124 |
| Amygdalin | 7 | 57 | -34 |
| Bergenin | 7 | 58 | -100 |
| Daidzin | 7 | 59 | -14 |
| Gramine | 7 | 60 | -61 |
| Betaxolol | 7 | 61 | 1 |
| Naltrexone HCl | 7 | 62 | -21 |
| Conivaptan HCl | 7 | 63 | 92 |
| S-(+)-Rolipram | 7 | 64 | 29 |
| Rosuvastatin Calcium | 7 | 65 | 24 |
| Apatinib Mesylate | 7 | 66 | 94 |
| Andrographolide | 7 | 67 | 31 |
| Berberine Chloride | 7 | 68 | -40 |
| Dihydroartemisinin | 7 | 69 | 20 |
| Hesperidin | 7 | 70 | 1 |
| Detomidine HCl | 7 | 71 | 4 |
| Levosulpiride | 7 | 72 | -4 |
| Ibutilide Fumarate | 7 | 73 | -89 |
| Bazedoxifene HCl | 7 | 74 | 68 |
| Telotristat Etiprate | 7 | 75 | 88 |
| Idelalisib | 7 | 76 | 40 |
| Apigenin | 7 | 77 | -6 |
| Caffeic Acid | 7 | 78 | -55 |
| DL-Carnitine HCl | 7 | 79 | -19 |
| Honokiol | 7 | 80 | 42 |
| Hyodeoxycholic Acid (HDCA) | 8 | 1 | 15 |
| Nalidixic Acid | 8 | 2 | 46 |
| Paeonol | 8 | 3 | 46 |
| Sclareol | 8 | 4 | 88 |

| | | | |
|---|---|---|---|
| Troxerutin | 8 | 5 | 10 |
| D-Mannitol | 8 | 6 | -2 |
| Sorbitol | 8 | 7 | -16 |
| Rotundine | 8 | 8 | 20 |
| Amfebutamone (Bupropion) HCl | 8 | 9 | 39 |
| Pramipexole | 8 | 10 | -54 |
| Kaempferol | 8 | 11 | 19 |
| Naringin | 8 | 12 | 50 |
| (-)-Parthenolide | 8 | 13 | 29 |
| Silibinin | 8 | 14 | -5 |
| Ursolic Acid | 8 | 15 | 36 |
| Gastrodin | 8 | 16 | -64 |
| Salidroside | 8 | 17 | 34 |
| Synephrine HCl | 8 | 18 | 36 |
| Benserazide HCl | 8 | 19 | -5 |
| Domperidone | 8 | 20 | 92 |
| Kinetin | 8 | 21 | 78 |
| Neohesperidin Dihydrochalcone | 8 | 22 | 53 |
| Piperine | 8 | 23 | 66 |
| Silymarin | 8 | 24 | 40 |
| Yohimbine HCl | 8 | 25 | -24 |
| Hematoxylin | 8 | 26 | 30 |
| Palmatine Chloride | 8 | 27 | 7 |
| Guanosine | 8 | 28 | 12 |
| Bupivacaine HCl | 8 | 29 | 53 |
| Estriol | 8 | 30 | 3 |
| L-(+)-Rhamnose Monohydrate | 8 | 31 | 21 |
| Neohesperidin | 8 | 32 | 14 |
| Puerarin | 8 | 33 | 52 |
| Sinomenine | 8 | 34 | -28 |
| S-hydroxytryptophan | 8 | 35 | -2 |
| Indirubin | 8 | 36 | 40 |
| Sodium Danshensu | 8 | 37 | -16 |
| Inosine | 8 | 38 | 37 |
| Bethanechol Chloride | 8 | 39 | 2 |
| Famciclovir | 8 | 40 | -6 |
| Lappaconitine | 8 | 41 | 76 |
| Oleanolic Acid | 8 | 42 | 12 |
| Quercetin Dihydrate | 8 | 43 | 68 |
| Synephrine | 8 | 44 | -8 |
| Aloin | 8 | 45 | 38 |
| L-carnitine | 8 | 46 | 18 |
| Paeoniflorin | 8 | 47 | 31 |
| Tolbutamide | 8 | 48 | 56 |
| Chlorpromazine HCl | 8 | 49 | 96 |
| Fenbendazole | 8 | 50 | 28 |
| Luteolin | 8 | 51 | 48 |

| | | | |
|---|---|---|---|
| Orotic Acid | 8 | 52 | -27 |
| Rotenone | 8 | 53 | 83 |
| Tanshinone I | 8 | 54 | -69 |
| Ammonium Glycyrrhizinate | 8 | 55 | -28 |
| Naringin Dihydrochalcone | 8 | 56 | -27 |
| Geniposide | 8 | 57 | -37 |
| Levosimendan | 8 | 58 | 38 |
| Clindamycin HCl | 8 | 59 | 56 |
| Fluocinolone Acetonide | 8 | 60 | 44 |
| Magnolol | 8 | 61 | 80 |
| Osthole | 8 | 62 | 24 |
| Rutin | 8 | 63 | 25 |
| Tanshinone IIA | 8 | 64 | -64 |
| Butylscopolamine Bromide | 8 | 65 | 1 |
| Polydatin | 8 | 66 | 27 |
| Ipriflavone | 8 | 67 | -50 |
| Equol | 8 | 68 | 29 |
| Clonidine HCl | 8 | 69 | -15 |
| Gallamine Triethiodide | 8 | 70 | -57 |
| Morin Hydrate | 8 | 71 | 29 |
| Oxymatrine | 8 | 72 | 22 |
| Salicin | 8 | 73 | 14 |
| Taxifolin | 8 | 74 | -6 |
| Diosmetin | 8 | 75 | 38 |
| Quercetin | 8 | 76 | 53 |
| (S)-10-Hydroxycamptothecin | 8 | 77 | 100 |
| Amantadine HCl | 8 | 78 | -59 |
| Clozapine | 8 | 79 | 38 |
| Hexestrol | 8 | 80 | 0 |
| Imatinib (STI571) | 9 | 1 | -83 |
| Mycophenolic acid | 9 | 2 | 18 |
| Phenoxybenzamine HCl | 9 | 3 | -133 |
| Scopolamine HBr | 9 | 4 | -28 |
| Naphazoline HCl | 9 | 5 | -47 |
| Isoconazole nitrate | 9 | 6 | 43 |
| Tiotropium Bromide hydrate | 9 | 7 | -79 |
| Rosiglitazone | 9 | 8 | -17 |
| Phenylephrine HCl | 9 | 9 | -24 |
| Thiamphenicol | 9 | 10 | -29 |
| Lincomycin HCl | 9 | 11 | -55 |
| Nateglinide | 9 | 12 | 66 |
| Propafenone HCl | 9 | 13 | 72 |
| Sotalol HCl | 9 | 14 | -4 |
| Epinephrine bitartrate | 9 | 15 | 18 |
| Econazole nitrate | 9 | 16 | 82 |
| Trospium chloride | 9 | 17 | -23 |
| Terbinafine HCl | 9 | 18 | -138 |

| | | | |
|---|---|---|---|
| Prednisolone Acetate | 9 | 19 | -9 |
| Clobetasol propionate | 9 | 20 | 73 |
| Loperamide HCl | 9 | 21 | 53 |
| Nitrendipine | 9 | 22 | 59 |
| Pyrantel Pamoate | 9 | 23 | -79 |
| Spectinomycin 2HCl | 9 | 24 | -7 |
| L-Adrenaline | 9 | 25 | -11 |
| Miconazole | 9 | 26 | 59 |
| Tolterodine tartrate | 9 | 27 | -88 |
| Cortisone acetate | 9 | 28 | 33 |
| Tetracaine HCl | 9 | 29 | 31 |
| Brompheniramine hydrogen maleate | 9 | 30 | -96 |
| Manidipine | 9 | 31 | -62 |
| Novobiocin Sodium | 9 | 32 | -27 |
| Quinine HCl Dihydrate | 9 | 33 | -94 |
| Sulfadoxine | 9 | 34 | -21 |
| Phenytoin Sodium | 9 | 35 | 0 |
| Secnidazole | 9 | 36 | 5 |
| Sulbactam sodium | 9 | 37 | 15 |
| Clomifene citrate | 9 | 38 | -125 |
| Tetracycline HCl | 9 | 39 | -39 |
| Dimethyl Fumarate | 9 | 40 | 0 |
| Manidipine 2HCl | 9 | 41 | 96 |
| Olanzapine | 9 | 42 | -59 |
| Racecadotril | 9 | 43 | -77 |
| Tenoxicam | 9 | 44 | 11 |
| Phenytoin | 9 | 45 | 7 |
| Acetanilide | 9 | 46 | 1 |
| Azelastine HCl | 9 | 47 | -4 |
| Cloxacillin Sodium | 9 | 48 | -18 |
| Xylometazoline HCl | 9 | 49 | -112 |
| Miglitol | 9 | 50 | -40 |
| Milrinone | 9 | 51 | 1 |
| Olopatadine HCl | 9 | 52 | -19 |
| Ribavirin | 9 | 53 | -13 |
| Vardenafil HCl Trihydrate | 9 | 54 | 84 |
| Ciclopirox | 9 | 55 | 34 |
| Clomipramine HCl | 9 | 56 | 79 |
| 5-Aminolevulinic acid HCl | 9 | 57 | -3 |
| Amoxicillin Sodium | 9 | 58 | -8 |
| Phenacetin | 9 | 59 | -17 |
| Pioglitazone | 9 | 60 | -106 |
| Mitoxantrone 2HCl | 9 | 61 | -590 |
| Ozagrel | 9 | 62 | -99 |
| Rosiglitazone maleate | 9 | 63 | 5 |
| Xylazine HCl | 9 | 64 | -46 |
| Dopamine HCl | 9 | 65 | -35 |

| | | | |
|---|---|---|---|
| Phenformin HCl | 9 | 66 | -69 |
| Daphnetin | 9 | 67 | -14 |
| Isoprenaline HCl | 9 | 68 | -13 |
| Zidovudine | 9 | 69 | -24 |
| Tolvaptan | 9 | 70 | 67 |
| Moroxydine HCl | 9 | 71 | -2 |
| Pancuronium dibromide | 9 | 72 | -113 |
| Roxithromycin | 9 | 73 | -19 |
| Maprotiline HCl | 9 | 74 | -26 |
| Ritodrine HCl | 9 | 75 | -65 |
| Ceftiofur HCl | 9 | 76 | -12 |
| Clarithromycin | 9 | 77 | -2 |
| Medroxyprogesterone acetate | 9 | 78 | -11 |
| Quinapril HCl | 9 | 79 | 28 |
| Pramiracetam | 9 | 80 | -11 |
| Clindamycin Palmitate HCl | 10 | 1 | -8 |
| Buflomedil HCl | 10 | 2 | -59 |
| Clinofibrate | 10 | 3 | -16 |
| Canagliflozin | 10 | 4 | -22 |
| Dabrafenib | 10 | 5 | -54 |
| Alogliptin Benzoate | 10 | 6 | -45 |
| Icotinib | 10 | 7 | -16 |
| Amoxicillin | 10 | 8 | -60 |
| Fenoprofen Calcium Hydrate | 10 | 9 | -54 |
| Cinepazide Maleate | 10 | 10 | -11 |
| L-Thyroxine | 10 | 11 | -72 |
| Fluocinonide | 10 | 12 | 83 |
| Ciprofibrate | 10 | 13 | -44 |
| Alectinib | 10 | 14 | 2 |
| MPEP | 10 | 15 | -51 |
| Camostat Mesilate | 10 | 16 | -121 |
| Carbazochrome Sodium Sulfonate | 10 | 17 | -65 |
| Aspirin | 10 | 18 | -36 |
| Linagliptin | 10 | 19 | -95 |
| Otilonium Bromide | 10 | 20 | 56 |
| Gliclazide | 10 | 21 | 3 |
| Inulin | 10 | 22 | -16 |
| Dolutegravir | 10 | 23 | 2 |
| MK-2048 | 10 | 24 | -7 |
| Alpelisib | 10 | 25 | 37 |
| Prucalopride | 10 | 26 | -91 |
| Clevudine | 10 | 27 | -20 |
| Niflumic Acid | 10 | 28 | -127 |
| Vildagliptin | 10 | 29 | -7 |
| Bosentan Hydrate | 10 | 30 | -55 |
| Acemetacin | 10 | 31 | 26 |
| Lonidamine | 10 | 32 | 19 |

| | | | |
|---|---|---|---|
| Trametinib | 10 | 33 | 13 |
| Laquinimod | 10 | 34 | 7 |
| Clindamycin | 10 | 35 | -17 |
| Acesulfame Potassium | 10 | 36 | -41 |
| Rivaroxaban | 10 | 37 | 70 |
| Ciclopiroxethanolamine | 10 | 38 | -13 |
| Daunorubicin HCl | 10 | 39 | -124 |
| Rupatadine Fumarate | 10 | 40 | 1 |
| Tioxolone | 10 | 41 | -49 |
| Clorsulon | 10 | 42 | -9 |
| Ibrutinib | 10 | 43 | 51 |
| Tofacitinib | 10 | 44 | -10 |
| Epiandrosterone | 10 | 45 | -27 |
| Cobicistat | 10 | 46 | -91 |
| Prostaglandin E2 | 10 | 47 | -34 |
| Rimonabant | 10 | 48 | -28 |
| Pravastatin Sodium | 10 | 49 | 0 |
| Azelnidipine | 10 | 50 | 9 |
| Dehydroepiandrosterone | 10 | 51 | 39 |
| Arecoline HBr | 10 | 52 | -5 |
| Nilvadipine | 10 | 53 | 15 |
| Istradefylline | 10 | 54 | 19 |
| Apalutamide | 10 | 55 | 57 |
| S-Ruxolitinib | 10 | 56 | -16 |
| Paroxetine HCl | 10 | 57 | -88 |
| Cabazitaxel | 10 | 58 | 86 |
| Bepotastine Besilate | 10 | 59 | -76 |
| Alverine Citrate | 10 | 60 | -134 |
| Idebenone | 10 | 61 | -17 |
| Noradrenaline Bitartrate Monohydrat | 10 | 62 | -13 |
| Dacomitinib | 10 | 63 | 31 |
| Torcetrapib | 10 | 64 | 4 |
| Baricitinib | 10 | 65 | -10 |
| Lumiracoxib | 10 | 66 | -2 |
| Zaltoprofen | 10 | 67 | -85 |
| Bufexamac | 10 | 68 | -31 |
| Fosaprepitant Dimeglumine Salt | 10 | 69 | 63 |
| Azilsartan Medoxomil | 10 | 70 | -44 |
| Mifepristone | 10 | 71 | 48 |
| Fostamatinib | 10 | 72 | -7 |
| Niraparib | 10 | 73 | -91 |
| Sofosbuvir | 10 | 74 | -18 |
| Carfilzomib | 10 | 75 | 22 |
| Pirfenidone | 10 | 76 | -46 |
| Pazopanib | 10 | 77 | 61 |
| Lamotrigine | 10 | 78 | -14 |
| Rofecoxib | 10 | 79 | -38 |

| | | | |
|---|---|---|---|
| Medetomidine HCl | 10 | 80 | -15 |
| Bleomycin Sulfate | 11 | 1 | -63 |
| Chlorhexidine 2HCl | 11 | 2 | 31 |
| Atovaquone | 11 | 3 | -105 |
| Pyridoxine HCl | 11 | 4 | -17 |
| Biotin (Vitamin B7) | 11 | 5 | 91 |
| Entacapone | 11 | 6 | -34 |
| Tylosin Tartrate | 11 | 7 | -27 |
| Brinzolamide | 11 | 8 | -12 |
| Ropinirole HCl | 11 | 9 | -63 |
| Iopromide | 11 | 10 | -28 |
| Clofarabine | 11 | 11 | 8 |
| Piracetam | 11 | 12 | -32 |
| Etravirine | 11 | 13 | 58 |
| Vitamin C | 11 | 14 | -18 |
| Sulfamerazine | 11 | 15 | -7 |
| Estradiol Valerate | 11 | 16 | 26 |
| Benztropine Mesylate | 11 | 17 | -73 |
| Carbenicillin Disodium | 11 | 18 | 37 |
| Ticarcillin Sodium | 11 | 19 | -4 |
| Fexofenadine HCl | 11 | 20 | -13 |
| Dacarbazine | 11 | 21 | -56 |
| Vanillin | 11 | 22 | -8 |
| Ulipristal | 11 | 23 | 58 |
| Sulfathiazole | 11 | 24 | 2 |
| Sulfamethazine | 11 | 25 | -31 |
| Articaine HCl | 11 | 26 | -41 |
| Altrenogest | 11 | 27 | 29 |
| Eletriptan HBr | 11 | 28 | -163 |
| Azlocillin Sodium Salt | 11 | 29 | -58 |
| Moclobemide | 11 | 30 | -74 |
| Dexrazoxane HCl | 11 | 31 | 25 |
| Chlorthalidone | 11 | 32 | -5 |
| Indacaterol Maleate | 11 | 33 | 84 |
| Oxybutynin Chloride | 11 | 34 | -22 |
| Sodium Salicylate | 11 | 35 | -34 |
| Gliquidone | 11 | 36 | 65 |
| Ampicillin Sodium | 11 | 37 | -64 |
| Flumequine | 11 | 38 | -9 |
| Reboxetine Mesylate | 11 | 39 | 36 |
| Triptolide | 11 | 40 | -8 |
| Epinephrine HCl | 11 | 41 | -28 |
| Dexmedetomidine | 11 | 42 | 12 |
| 2-Thiouracil | 11 | 43 | -12 |
| Doxepin HCl | 11 | 44 | -59 |
| Methylthiouracil | 11 | 45 | -3 |
| Butenafine HCl | 11 | 46 | -45 |

| | | | |
|---|---|---|---|
| Anagrelide HCl | 11 | 47 | 13 |
| Amitriptyline HCl | 11 | 48 | 43 |
| Triflusal | 11 | 49 | -30 |
| Borneol | 11 | 50 | 15 |
| Diclofenac Potassium | 11 | 51 | -38 |
| Moguisteine | 11 | 52 | 15 |
| Ornidazole | 11 | 53 | -13 |
| Mepivacaine HCl | 11 | 54 | -63 |
| Antipyrine | 11 | 55 | -14 |
| Adrenalone HCl | 11 | 56 | -38 |
| Trifluoperazine 2HCl | 11 | 57 | -14 |
| Fangchinoline | 11 | 58 | 63 |
| Diclofenac Diethylamine | 11 | 59 | 3 |
| Tazobactam | 11 | 60 | -20 |
| Nadifloxacin | 11 | 61 | 4 |
| Dexamethasone Acetate | 11 | 62 | 67 |
| Milnacipran HCl | 11 | 63 | -131 |
| Ethynodiol Diacetate | 11 | 64 | -1 |
| Atomoxetine HCl | 11 | 65 | 0 |
| Azatadine Dimaleate | 11 | 66 | -57 |
| Catharanthine | 11 | 67 | 22 |
| Berbamine | 11 | 68 | 12 |
| Naloxone HCl | 11 | 69 | -48 |
| Beclomethasone Dipropionate | 11 | 70 | 58 |
| Pidotimod | 11 | 71 | -29 |
| Trimethoprim | 11 | 72 | -59 |
| Darifenacin HBr | 11 | 73 | -23 |
| Sertaconazole Nitrate | 11 | 74 | 22 |
| Betahistine 2HCl | 11 | 75 | -54 |
| (+,-)-Octopamine HCl | 11 | 76 | -36 |
| Meptazinol HCl | 11 | 77 | -80 |
| (+)-Fangchinoline | 11 | 78 | 43 |
| Rosmarinic acid | 12 | 1 | -49 |
| Medroxyprogesterone | 12 | 2 | -14 |
| Sulfamonomethoxine | 12 | 3 | -21 |
| Promestriene | 12 | 4 | 38 |
| Quinestrol | 12 | 5 | -4 |
| 4-Biphenylacetic acid | 12 | 6 | -22 |
| Phenethyl alcohol | 12 | 7 | -7 |
| Lifitegrast | 12 | 8 | -38 |
| Selexipag | 12 | 9 | -121 |
| Umeclidinium bromide | 12 | 10 | 36 |
| Dehydrocostus Lactone | 12 | 11 | -78 |
| Kitasamycin | 12 | 12 | 69 |
| Fludrocortisone acetate | 12 | 13 | 9 |
| Cefonicid sodium | 12 | 14 | -90 |
| Ethidium bromide | 12 | 15 | 4 |

| | | | |
|---|---|---|---|
| 4-Methylbenzylidene camphor | 12 | 16 | -103 |
| Flibanserin | 12 | 17 | 78 |
| Vilanterol Trifenate | 12 | 18 | 67 |
| Travoprost | 12 | 19 | -125 |
| Asiaticoside | 12 | 20 | 16 |
| Cefpirome sulfate | 12 | 21 | -87 |
| Thimerosal | 12 | 22 | -46 |
| Afloqualone | 12 | 23 | 7 |
| Sulfaphenazole | 12 | 24 | -53 |
| Pargyline hydrochloride | 12 | 25 | -32 |
| Chlorobutanol | 12 | 26 | -35 |
| Testosterone Enanthate | 12 | 27 | 52 |
| Grazoprevir | 12 | 28 | -102 |
| Calcipotriene | 12 | 29 | 79 |
| Acetylspiramycin (ASPM) | 12 | 30 | 3 |
| Cefamandole nafate | 12 | 31 | -102 |
| Mafenide Acetate | 12 | 32 | -25 |
| Flupenthixol dihydrochloride | 12 | 33 | -104 |
| Levamlodipine | 12 | 34 | 67 |
| N-Ethylmaleimide (NEM) | 12 | 35 | 13 |
| Sarpogrelate hydrochloride | 12 | 36 | 59 |
| Leuprolide Acetate | 12 | 37 | -70 |
| Iron sucrose | 12 | 38 | -139 |
| Benznidazole | 12 | 39 | -22 |
| Pazufloxacin mesylate | 12 | 40 | 21 |
| Tacrine hydrochloride hydrate | 12 | 41 | -105 |
| Amlexanox | 12 | 42 | 9 |
| Ilaprazole | 12 | 43 | 59 |
| Umbelliferone | 12 | 44 | -41 |
| Glucosamine hydrochloride | 12 | 45 | 10 |
| Ethopabate | 12 | 46 | -13 |
| Topiroxostat | 12 | 47 | -26 |
| Metaxalone | 12 | 48 | 71 |
| Cholic acid | 12 | 49 | 50 |
| Diammonium Glycyrrhizinate | 12 | 50 | -24 |
| Methoxyphenamine Hydrochloride | 12 | 51 | -72 |
| Tauroursodeoxycholic Acid (TUDCA) | 12 | 52 | -22 |
| Thymopentin | 12 | 53 | -69 |
| Cinnamic acid | 12 | 54 | -21 |
| Mafenide hydrochloride | 12 | 55 | -38 |
| Sulfachloropyridazine | 12 | 56 | -52 |
| Isavuconazole | 12 | 57 | -10 |
| Tipiracil hydrochloride | 12 | 58 | -30 |
| Ceftibuten dihydrate | 12 | 59 | -100 |
| Osalmid | 12 | 60 | -2 |
| Cefepime Dihydrochloride Monohydra | 12 | 61 | -11 |
| Carmustine | 12 | 62 | -37 |

| | | | |
|---|---|---|---|
| Vitamin E Acetate | 12 | 63 | -99 |
| Nortriptyline hydrochloride | 12 | 64 | -41 |
| Carbasalate Calcium | 12 | 65 | -8 |
| Ramosetron Hydrochloride | 12 | 66 | -55 |
| Avibactam sodium | 12 | 67 | -22 |
| Balsalazide disodium | 12 | 68 | -124 |
| Tyramine | 12 | 69 | 2 |
| Amitraz | 12 | 70 | 10 |
| Piribedil | 12 | 71 | 12 |
| Cefsulodin sodium | 12 | 72 | -25 |
| Methacholine chloride | 12 | 73 | 11 |
| Benactyzine hydrochloride | 12 | 74 | 55 |
| Moxidectin | 12 | 75 | 80 |
| Velpatasvir | 12 | 76 | 93 |
| Boceprevir | 12 | 77 | 52 |
| Lumefantrine | 12 | 78 | -107 |
| Levothyroxine sodium | 13 | 1 | -41 |
| Cinnamaldehyde | 13 | 2 | -42 |
| Echinacoside | 13 | 3 | -5 |
| Imperatorin | 13 | 4 | 35 |
| Panaxatriol | 13 | 5 | 77 |
| Harmine | 13 | 6 | -100 |
| Isopsoralen | 13 | 7 | -5 |
| Hederacoside C | 13 | 8 | 73 |
| Nonivamide | 13 | 9 | 79 |
| (+)-Borneol | 13 | 10 | -92 |
| Sodium benzoate | 13 | 11 | 29 |
| Tanshinone IIA sulfonate (sodium) | 13 | 12 | -34 |
| Notoginsenoside R1 | 13 | 13 | -264 |
| Scutellarin | 13 | 14 | -40 |
| D-Galactose | 13 | 15 | -19 |
| Guaiacol | 13 | 16 | 38 |
| Bornyl acetate | 13 | 17 | 17 |
| Lathyrol | 13 | 18 | 69 |
| Valproic acid | 13 | 19 | 14 |
| Vanillyl Butyl Ether | 13 | 20 | 54 |
| Quinidine sulfate | 13 | 21 | -29 |
| Palmatine | 13 | 22 | -112 |
| Carvacrol | 13 | 23 | 71 |
| Ginsenoside Re | 13 | 24 | 46 |
| Glucosamine sulfate | 13 | 25 | -4 |
| Indigo | 13 | 26 | -160 |
| Sophoridine | 13 | 27 | 1 |
| Ginsenoside Rg1 | 13 | 28 | 15 |
| L-Cycloserine | 13 | 29 | 44 |
| Nifuratel | 13 | 30 | 23 |
| 4-Hydroxybenzoic acid | 13 | 31 | 38 |

| | | | |
|---|---|---|---|
| 5-Hydroxymethylfurfural | 13 | 32 | 3 |
| Succinic acid | 13 | 33 | -1 |
| Harmine hydrochloride | 13 | 34 | -10 |
| Camphor | 13 | 35 | -2 |
| Scopoletin | 13 | 36 | -4 |
| Hydroxy Camptothecine | 13 | 37 | 99 |
| Ginsenoside Rb1 | 13 | 38 | 12 |
| Mesterolone | 13 | 39 | -36 |
| Flavone | 13 | 40 | -27 |
| Betaine | 13 | 41 | 41 |
| Tyrosol | 13 | 42 | 29 |
| Palmitic acid | 13 | 43 | -87 |
| Quercitrin | 13 | 44 | -43 |
| Tetrahydropalmatine hydrochloride | 13 | 45 | 59 |
| Protopine | 13 | 46 | -124 |
| Hederagenin | 13 | 47 | 71 |
| (-)-Epicatechin gallate | 13 | 48 | 18 |
| Spermine | 13 | 49 | -87 |
| Histamine | 13 | 50 | 4 |
| Methyl salicylate | 13 | 51 | 44 |
| Ligustrazine hydrochloride | 13 | 52 | 46 |
| Trigonelline Hydrochloride | 13 | 53 | 37 |
| Loganin | 13 | 54 | 49 |
| Allantoin | 13 | 55 | 4 |
| Pyrogallol | 13 | 56 | -49 |
| Astragaloside IV | 13 | 57 | 58 |
| Forsythin | 13 | 58 | 57 |
| Maltitol | 13 | 59 | 17 |
| Veratric acid | 13 | 60 | 33 |
| Sinomenine hydrochloride | 13 | 61 | 9 |
| cis-Anethole | 13 | 62 | 26 |
| Stevioside | 13 | 63 | 64 |
| Isoquercitrin | 13 | 64 | -27 |
| Lawsone | 13 | 65 | 4 |
| L-Rhamnose monohydrate | 13 | 66 | 27 |
| Catalpol | 13 | 67 | 40 |
| Swertiamarin | 13 | 68 | 24 |
| Tannic acid | 13 | 69 | 64 |
| Vindoline | 13 | 70 | 53 |
| Eucalyptol | 13 | 71 | 48 |
| Ginkgolide C | 13 | 72 | 19 |
| Dehydroandrographolide | 13 | 73 | 29 |
| Madecassoside | 13 | 74 | 53 |
| Galanthamine | 13 | 75 | 3 |
| Arteether | 13 | 76 | 37 |
| α-Hederin | 13 | 77 | -134 |
| Liquiritin | 13 | 78 | 59 |

| | | | |
|---|---|---|---|
| Gamma-Oryzanol | 13 | 79 | -10 |
| Fusidine | 13 | 80 | 26 |
| Lobeline Hydrochloride | 14 | 1 | -35 |
| Nordihydroguaiaretic Acid | 14 | 2 | 51 |
| (+)-α-Lipoic Acid | 14 | 3 | -62 |
| Acebutolol HCl | 14 | 4 | -82 |
| Sodium Picosulfate | 14 | 5 | -6 |
| Nafcillin Sodium | 14 | 6 | -81 |
| Valganciclovir HCl | 14 | 7 | -41 |
| Erythromycin Ethylsuccinate | 14 | 8 | 83 |
| Griseofulvin | 14 | 9 | 74 |
| (+)-Catechin Hydrate | 14 | 10 | -71 |
| Methyl 4-Hydroxybenzoate | 14 | 11 | -36 |
| Pergolide Mesylate | 14 | 12 | -57 |
| Ampiroxicam | 14 | 13 | -10 |
| Tolcapone | 14 | 14 | 3 |
| Diphemanil Methylsulfate | 14 | 15 | 6 |
| Tetrahydrozoline HCl | 14 | 16 | -129 |
| Nabumetone | 14 | 17 | 42 |
| Levobupivacaine HCl | 14 | 18 | 30 |
| Decamethonium Bromide | 14 | 19 | -57 |
| Protocatechuic Acid | 14 | 20 | -38 |
| L(+)-Arabinose | 14 | 21 | -33 |
| Cabozantinib Malate | 14 | 22 | 79 |
| Desloratadine | 14 | 23 | 52 |
| Probenecid | 14 | 24 | 45 |
| Vitamin D2 | 14 | 25 | -133 |
| Toltrazuril | 14 | 26 | 63 |
| Sertraline HCl | 14 | 27 | 64 |
| Ronidazole | 14 | 28 | -56 |
| Sodium Nitrite | 14 | 29 | -59 |
| (-)-Borneol | 14 | 30 | 68 |
| L-Tryptophan | 14 | 31 | -48 |
| Sitagliptin Phosphate Monohydrate | 14 | 32 | 54 |
| Hyoscyamine | 14 | 33 | 3 |
| Procaine HCl | 14 | 34 | 8 |
| Doxapram HCl | 14 | 35 | 51 |
| Pheniramine Maleate | 14 | 36 | -88 |
| Spironolactone | 14 | 37 | -15 |
| Cholecaciferol | 14 | 38 | -59 |
| Zinc Pyrithione | 14 | 39 | -84 |
| Zinc Undecylenate | 14 | 40 | 23 |
| D-(+)-Trehalose Dihydrate | 14 | 41 | 19 |
| Lithocholoic Acid | 14 | 42 | 51 |
| Ouabai | 14 | 43 | 18 |
| Homatropine Methylbromide | 14 | 44 | -34 |
| Dibucaine HCl | 14 | 45 | 74 |

| | | | |
|---|---|---|---|
| Estradiol Cypionate | 14 | 46 | 58 |
| Retapamulin | 14 | 47 | 40 |
| Escitalopram Oxalate | 14 | 48 | 25 |
| Propranolol HCl | 14 | 49 | -52 |
| Pyridoxine | 14 | 50 | 40 |
| Guaiazulene | 14 | 51 | -20 |
| Ethambutol 2HCl | 14 | 52 | -35 |
| Allylthiourea | 14 | 53 | -1 |
| Homatropine Bromide | 14 | 54 | -56 |
| Methazolamide | 14 | 55 | -59 |
| Bisacodyl | 14 | 56 | 58 |
| Methyclothiazide | 14 | 57 | 25 |
| Guanabenz Acetate | 14 | 58 | -2 |
| Mequinol | 14 | 59 | 25 |
| Batyl Alcohol | 14 | 60 | -43 |
| Thioctic Acid | 14 | 61 | -74 |
| Pentamidine Isethionate | 14 | 62 | -17 |
| Sennoside B | 14 | 63 | -4 |
| Hydroxyzine 2HCl | 14 | 64 | 93 |
| Norethindrone | 14 | 65 | -22 |
| Carbimazole | 14 | 66 | 8 |
| Ropivacaine HCl | 14 | 67 | -19 |
| Tinidazole | 14 | 68 | -17 |
| Mefenamic Acid | 14 | 69 | 15 |
| Caryophyllene Oxide | 14 | 70 | -21 |
| Oxaceprol | 14 | 71 | -44 |
| Mirabegron | 14 | 72 | 1 |
| Avanafil | 14 | 73 | 99 |
| Aclidinium Bromide | 14 | 74 | 38 |
| Olsalazine Sodium | 14 | 75 | 21 |
| Valdecoxib | 14 | 76 | 20 |
| Sodium Nitroprusside Dihydrate | 14 | 77 | 79 |
| Guanidine HCl | 14 | 78 | -18 |
| Ticagrelor | 14 | 79 | -49 |
| Triamterene | 15 | 1 | 91 |
| Halobetasol Propionate | 15 | 2 | 72 |
| Esmolol HCl | 15 | 3 | -66 |
| Dicloxacillin Sodium | 15 | 4 | -29 |
| Timolol Maleate | 15 | 5 | -48 |
| Cyclizine 2HCl | 15 | 6 | 29 |
| Chlorzoxazone | 15 | 7 | 5 |
| Chlorpropamide | 15 | 8 | 19 |
| Trometamol | 15 | 9 | -16 |
| Domiphen Bromide | 15 | 10 | -51 |
| Sulfacetamide Sodium | 15 | 11 | -13 |
| Fenspiride HCl | 15 | 12 | -77 |
| Voglibose | 15 | 13 | -16 |

| | | | |
|---|---|---|---|
| Desvenlafaxine Succinate | 15 | 14 | -35 |
| Tolazoline HCl | 15 | 15 | -45 |
| Dinitolmide | 15 | 16 | -65 |
| Bezafibrate | 15 | 17 | -7 |
| Cyromazine | 15 | 18 | -15 |
| Uracil | 15 | 19 | -27 |
| Salicylanilide | 15 | 20 | -2 |
| Spiramycin | 15 | 21 | -5 |
| Ifenprodil Tartrate | 15 | 22 | -27 |
| Eprosartan Mesylate | 15 | 23 | 20 |
| Desvenlafaxine | 15 | 24 | -39 |
| Sodium Phenylbutyrate | 15 | 25 | -6 |
| Pentoxyverine Citrate | 15 | 26 | -14 |
| Penicillin G Sodium | 15 | 27 | -4 |
| Teriflunomide | 15 | 28 | -28 |
| Climbazole | 15 | 29 | 62 |
| Sasapyrine | 15 | 30 | 7 |
| Vitamin A Acetate | 15 | 31 | 72 |
| Pramoxine HCl | 15 | 32 | -176 |
| Diminazene Aceturate | 15 | 33 | -77 |
| Triclabendazole | 15 | 34 | 7 |
| Troxipide | 15 | 35 | -31 |
| Azithromycin Dihydrate | 15 | 36 | 4 |
| Benzoic Acid | 15 | 37 | -18 |
| Coumarin | 15 | 38 | -47 |
| Mezlocillin Sodium | 15 | 39 | -44 |
| Cyclandelate | 15 | 40 | 56 |
| Lomerizine 2HCl | 15 | 41 | 86 |
| Difluprednate | 15 | 42 | 92 |
| Closantel Sodium | 15 | 43 | -16 |
| Histamine 2HCl | 15 | 44 | -50 |
| Levodropropizine | 15 | 45 | -32 |
| Ampicillin Trihydrate | 15 | 46 | -75 |
| Benzethonium Chloride | 15 | 47 | -123 |
| Choline Chloride | 15 | 48 | -15 |
| Nicardipine HCl | 15 | 49 | 26 |
| Betamipron | 15 | 50 | -16 |
| Levobetaxolol HCl | 15 | 51 | -29 |
| Droperidol | 15 | 52 | 29 |
| Closantel | 15 | 53 | 21 |
| Pefloxacin Mesylate Dihydrate | 15 | 54 | -59 |
| Clorprenaline HCl | 15 | 55 | -45 |
| Amfenac Sodium Monohydrate | 15 | 56 | 31 |
| Doxycycline Hyclate | 15 | 57 | -21 |
| Cetylpyridinium Chloride | 15 | 58 | -69 |
| Nifuroxazide | 15 | 59 | 77 |
| Chlorquinaldol | 15 | 60 | 27 |

| | | | |
|---|---|---|---|
| Loxapine Succinate | 15 | 61 | 92 |
| Halcinonide | 15 | 62 | -48 |
| Clofazimine | 15 | 63 | -77 |
| Sulconazole Nitrate | 15 | 64 | 94 |
| Carprofen | 15 | 65 | -9 |
| Penfluridol | 15 | 66 | -27 |
| Doxofylline | 15 | 67 | -3 |
| 1-Hexadecanol | 15 | 68 | -35 |
| Penciclovir | 15 | 69 | -5 |
| Broxyquinoline | 15 | 70 | -3 |
| Flumethasone | 15 | 71 | 10 |
| Dexlansoprazole | 15 | 72 | 44 |
| Estradiol Benzoate | 15 | 73 | 93 |
| Tilmicosin | 15 | 74 | -52 |
| Dropropizine | 15 | 75 | -13 |
| Ethamsylate | 15 | 76 | 8 |
| Benzydamine HCl | 15 | 77 | 24 |
| Sulfaguanidine | 15 | 78 | -13 |
| Tiratricol | 15 | 79 | 25 |
| Ethacridine lactate monohydrate | 15 | 80 | -17 |
| Bemegride | 16 | 1 | -32 |
| Chromocarb | 16 | 2 | -134 |
| Azaperone | 16 | 3 | -12 |
| Oxybuprocaine HCl | 16 | 4 | 82 |
| Cetrimonium Bromide | 16 | 5 | 89 |
| Mechlorethamine HCl | 16 | 6 | -90 |
| Efaproxiral Sodium | 16 | 7 | -30 |
| Rotigotine | 16 | 8 | 76 |
| Chloroprocaine HCl | 16 | 9 | -95 |
| Promethazine HCl | 16 | 10 | 67 |
| Tolperisone HCl | 16 | 11 | -157 |
| Chlorocresol | 16 | 12 | 8 |
| Benzbromarone | 16 | 13 | -83 |
| Oxaprozin | 16 | 14 | -15 |
| Deoxycorticosterone Acetate | 16 | 15 | 21 |
| Epinastine HCl | 16 | 16 | -52 |
| Etofibrate | 16 | 17 | 69 |
| Bambuterol HCl | 16 | 18 | -28 |
| Ospemifene | 16 | 19 | 63 |
| Procainamide HCl | 16 | 20 | -82 |
| Florfenicol | 16 | 21 | -227 |
| Benzocaine | 16 | 22 | -9 |
| Piperacillin Sodium | 16 | 23 | -16 |
| Pilocarpine HCl | 16 | 24 | -27 |
| Serotonin HCl | 16 | 25 | -114 |
| Quinacrine 2HCl | 16 | 26 | -13 |
| Nicaraven | 16 | 27 | -23 |

| | | | |
|---|---|---|---|
| Carteolol HCl | 16 | 28 | -64 |
| Anidulafungin | 16 | 29 | 99 |
| Meclofenamate Sodium | 16 | 30 | -86 |
| Verapamil HCl | 16 | 31 | 72 |
| Montelukast Sodium | 16 | 32 | -26 |
| Mevastatin | 16 | 33 | 94 |
| Phenazopyridine HCl | 16 | 34 | 70 |
| Tranylcypromine HCl | 16 | 35 | -1 |
| Busiprone HCl | 16 | 36 | -23 |
| Brimonidine Tartrate | 16 | 37 | -21 |
| Demeclocycline HCl | 16 | 38 | 22 |
| Chloroambucil | 16 | 39 | -17 |
| Salmeterol Xinafoate | 16 | 40 | 63 |
| Furaltadone HCl | 16 | 41 | -52 |
| Dirithromycin | 16 | 42 | 29 |
| Erythritol | 16 | 43 | -38 |
| Primaquine Diphosphate | 16 | 44 | 15 |
| Prucalopride Succinate | 16 | 45 | -122 |
| Alizapride HCl | 16 | 46 | -109 |
| Diacerein | 16 | 47 | -19 |
| Meclofenoxate HCl | 16 | 48 | -17 |
| Metoclopramide HCl | 16 | 49 | 13 |
| Mupirocin | 16 | 50 | 60 |
| Isosorbide | 16 | 51 | -1 |
| Sucralose | 16 | 52 | -31 |
| Mexiletine HCl | 16 | 53 | -20 |
| Cepharanthine | 16 | 54 | 94 |
| Bromfenac Sodium | 16 | 55 | -15 |
| Luliconazole | 16 | 56 | 75 |
| Flufenamic Acid | 16 | 57 | 28 |
| Tasimelteon | 16 | 58 | 41 |
| Digoxin | 16 | 59 | 18 |
| Dicoumarol | 16 | 60 | -5 |
| Cysteamine HCl | 16 | 61 | 27 |
| Valnemulin HCl | 16 | 62 | 89 |
| Fidaxomicin | 16 | 63 | 47 |
| Bergapten | 16 | 64 | 37 |
| Flopropione | 16 | 65 | 51 |
| Vilazodone HCl | 16 | 66 | 99 |
| Vinorelbine Tartrate | 16 | 67 | 49 |
| Nelfinavir Mesylate | 16 | 68 | 94 |
| Labetalol HCl | 16 | 69 | -7 |
| R-(+)-Atenolol HCl | 16 | 70 | 4 |
| Clofibric Acid | 16 | 71 | 51 |
| Liothyronine Sodium | 16 | 72 | 14 |
| Fluorometholone Acetate | 16 | 73 | 95 |
| Doxylamine Succinate | 16 | 74 | -2 |

| | | | |
|---|---|---|---|
| Sulfamethoxypyridazine | 16 | 75 | -3 |
| Tamibarotene | 16 | 76 | -61 |
| Oxiracetam | 16 | 77 | -72 |
| Cyclobenzaprine HCl | 16 | 78 | 54 |
| Diphenidol HCl | 16 | 79 | 15 |
| Anisindione | 16 | 80 | 52 |
| Anisotropine Methylbromide | 17 | 1 | -34 |
| Isoetharine Mesylate | 17 | 2 | -62 |
| Thiostrepton | 17 | 3 | 99 |
| Suxibuzone | 17 | 4 | 34 |
| Pyrilamine Maleate | 17 | 5 | -179 |
| Clofoctol | 17 | 6 | -65 |
| Trimipramine Maleate | 17 | 7 | 57 |
| Aceglutamide | 17 | 8 | -65 |
| Ethylparaben | 17 | 9 | -50 |
| Auranofin | 17 | 10 | 67 |
| Meclocycline Sulfosalicylate | 17 | 11 | 10 |
| Oxethazaine | 17 | 12 | 78 |
| Tacrine HCl | 17 | 13 | -118 |
| Carbenoxolone Sodium | 17 | 14 | -123 |
| Fosfomycin Tromethamine | 17 | 15 | -28 |
| Digoxigenin | 17 | 16 | -28 |
| Mefloquine HCl | 17 | 17 | -71 |
| Acetylleucine | 17 | 18 | -60 |
| Fenbufen | 17 | 19 | 7 |
| Benzthiazide | 17 | 20 | -16 |
| Medrysone | 17 | 21 | 37 |
| Oxprenolol HCl | 17 | 22 | -68 |
| Pimozide | 17 | 23 | 63 |
| Bendroflumet Hiazide | 17 | 24 | 0 |
| Diperodon HCl | 17 | 25 | 74 |
| Eltrombopag | 17 | 26 | 54 |
| Ademetionine Disulfate Tosylate | 17 | 27 | 96 |
| Fenofibric Acid | 17 | 28 | -58 |
| Bromocriptine Mesylate | 17 | 29 | 96 |
| Mesoridazine Besylate | 17 | 30 | -91 |
| Pentoxifylline | 17 | 31 | -22 |
| Carbachol | 17 | 32 | -38 |
| Dicyclomine HCl | 17 | 33 | -14 |
| Bentiromide | 17 | 34 | -31 |
| Oxeladin Citrate | 17 | 35 | 66 |
| 6-Mercaptopurine | 17 | 36 | -88 |
| (+)-Camphor | 17 | 37 | -16 |
| Furazolidone | 17 | 38 | 32 |
| Carbadox | 17 | 39 | 84 |
| Metaproterenol Sulfate | 17 | 40 | -69 |
| Piromidic Acid | 17 | 41 | -82 |

| | | | |
|---|---|---|---|
| Cinoxacin | 17 | 42 | -58 |
| Mepenzolate Bromide | 17 | 43 | -58 |
| Bephenium Hydroxynaphthoate | 17 | 44 | -58 |
| Pasiniazid | 17 | 45 | -63 |
| Vinblastine Sulfate | 17 | 46 | 27 |
| Cefotaxime Sodium | 17 | 47 | -85 |
| Iopamidol | 17 | 48 | -37 |
| Clorgyline HCl | 17 | 49 | 32 |
| Metaraminol Bitartrate | 17 | 50 | -60 |
| Procyclidine HCl | 17 | 51 | -24 |
| Glafenine HCl | 17 | 52 | 73 |
| Aceclidine HCl | 17 | 53 | -10 |
| Brucine Sulfate Salt Hydrate | 17 | 54 | -76 |
| Picrotoxinin | 17 | 55 | -11 |
| Acetazolamide | 17 | 56 | -34 |
| Chloroxylenol | 17 | 57 | 25 |
| Methylene Blue | 17 | 58 | -44 |
| Disopyramide Phosphate | 17 | 59 | -26 |
| Methoxamine HCl | 17 | 60 | -65 |
| Ractopamine HCl | 17 | 61 | -39 |
| Pthalylsulfacetamide | 17 | 62 | -25 |
| Imipramine HCl | 17 | 63 | -3 |
| Camylofin Chlorhydrate | 17 | 64 | 87 |
| Pindolol | 17 | 65 | -70 |
| 4-Aminoantipyrine | 17 | 66 | -66 |
| Nitrofurantoin | 17 | 67 | -46 |
| Ethoxzolamide | 17 | 68 | 64 |
| Meticrane | 17 | 69 | -23 |
| Terfenadine | 17 | 70 | 57 |
| Pinacidil | 17 | 71 | -10 |
| Proadifen HCl | 17 | 72 | -143 |
| Cephapirin Sodium | 17 | 73 | -111 |
| Misoprostol | 17 | 74 | -136 |
| 4-Aminobenzoic Acid | 17 | 75 | -57 |
| 2-Aminoheptane | 17 | 76 | -24 |
| Pantoprazole Sodium | 17 | 77 | -75 |
| Salicylic Acid | 18 | 1 | 31 |
| Azelaic Acid | 18 | 2 | 33 |
| Diatrizoic Acid | 18 | 3 | 20 |
| Sulfabenzamide | 18 | 4 | 5 |
| Succinylsulfathiazole | 18 | 5 | -1 |
| Cefixime | 18 | 6 | -60 |
| Diflunisal | 18 | 7 | -2 |
| Alcaftadine | 18 | 8 | 1 |
| Sodium Sulfadiazine | 18 | 9 | -18 |
| Ciclesonide | 18 | 10 | 66 |
| Triclosan | 18 | 11 | 74 |

| | | | |
|---|---|---|---|
| Bithionol | 18 | 12 | -17 |
| Diethylcarbamazine Citrate | 18 | 13 | 13 |
| Terpin Hydrate | 18 | 14 | -3 |
| Docusate Sodium | 18 | 15 | -58 |
| Lercanidipine Hydrochloride | 18 | 16 | 68 |
| Mebendazole | 18 | 17 | 92 |
| Ethosuximide | 18 | 18 | -22 |
| Cyproheptadine Hydrochloride | 18 | 19 | 54 |
| Cefmenoxime Hydrochloride | 18 | 20 | 38 |
| Trihexyphenidyl Hydrochloride | 18 | 21 | 17 |
| Bronopol | 18 | 22 | 68 |
| Diiodohydroxyquinoline | 18 | 23 | -2 |
| Tyloxapol | 18 | 24 | -114 |
| Amodiaquine Dihydrochloride Dihydra | 18 | 25 | 76 |
| Benzyl Benzoate | 18 | 26 | 72 |
| Dapson | 18 | 27 | -1 |
| (+/-)-Sulfinpyrazone | 18 | 28 | 14 |
| Teneligliptin Hydrobromide | 18 | 29 | -14 |
| Dantrolene Sodium Hemiheptahydrate | 18 | 30 | -40 |
| Trimetazidine Dihydrochloride | 18 | 31 | 43 |
| Carzenide | 18 | 32 | 45 |
| DL-Panthenol | 18 | 33 | 34 |
| Resorcinol | 18 | 34 | 46 |
| Dithranol | 18 | 35 | 89 |
| Benzyl Alcohol | 18 | 36 | 55 |
| Dextromethorphan Hydrobromide | 18 | 37 | 59 |
| Chlorotrianisene | 18 | 38 | -11 |
| Prasugrel Hydrochloride | 18 | 39 | 31 |
| Atipamezole Hydrochloride | 18 | 40 | 79 |
| Urethane | 18 | 41 | 22 |
| Citiolone | 18 | 42 | 47 |
| Fluphenazine Dihydrochloride | 18 | 43 | -119 |
| Hydroquinone | 18 | 44 | 19 |
| Nitroxoline | 18 | 45 | 19 |
| Clioquinol | 18 | 46 | -42 |
| Fenoldopam Mesylate | 18 | 47 | -2 |
| Diazoxide | 18 | 48 | 10 |
| Desogestrel | 18 | 49 | 89 |
| Atipamezole | 18 | 50 | 53 |
| Xylitol | 18 | 51 | 2 |
| Cloxiquine | 18 | 52 | -7 |
| Halothane | 18 | 53 | 19 |
| Triacetin | 18 | 54 | 23 |
| Chlormadinone Acetate | 18 | 55 | 79 |
| Acetohydroxamic Acid | 18 | 56 | 24 |
| Itopride Hydrochloride | 18 | 57 | -9 |
| Prochlorperazine Dimaleate Salt | 18 | 58 | -69 |

| | | | |
|---|---|---|---|
| Brexpiprazole | 18 | 59 | 32 |
| Etoricoxib | 18 | 60 | 29 |
| 8-Hydroxyquinoline | 18 | 61 | 88 |
| Danthron | 18 | 62 | 89 |
| Hexylresorcinol | 18 | 63 | 93 |
| Butamben | 18 | 64 | 81 |
| Cephalothin | 18 | 65 | 26 |
| Gallic Acid | 18 | 66 | 63 |
| Cefuroxime Sodium | 18 | 67 | 37 |
| Hexachlorophene | 18 | 68 | 1 |
| Lesinurad | 18 | 69 | 26 |
| Sulisobenzone | 18 | 70 | 35 |
| Aminoguanidine Hydrochloride | 18 | 71 | 33 |
| Dehydrocholic Acid | 18 | 72 | -13 |
| Piperazine | 18 | 73 | -13 |
| Butylparaben | 18 | 74 | 95 |
| Cefazolin Sodium | 18 | 75 | 26 |
| Levofloxacin Hydrate | 18 | 76 | 38 |
| Methylbenactyzine Bromide | 18 | 77 | 21 |
| Isosorbide Mononitrate | 18 | 78 | 45 |
| Tedizolid Phosphate | 18 | 79 | 11 |
| Sulpiride | 18 | 80 | 36 |
| Parecoxib | 19 | 1 | -116 |
| Rebeprazole sodium | 19 | 2 | -87 |
| Gluconolactone | 19 | 3 | -86 |
| Rivastigmine | 19 | 4 | -123 |
| Vitamin K1 | 19 | 5 | -2 |
| Evans Blue | 19 | 6 | 28 |
| Perphenazine | 19 | 7 | -104 |
| Sulfacetamide sodium salt hydrate | 19 | 8 | -62 |
| Harmaline | 19 | 9 | -22 |
| Daminozide | 19 | 10 | -86 |
| Eslicarbazepine Acetate | 19 | 11 | -67 |
| Sivelestat sodium tetrahydrate | 19 | 12 | -77 |
| Povidone iodine | 19 | 13 | 0 |
| Deoxycholic acid | 19 | 14 | 36 |
| Etretinate | 19 | 15 | 21 |
| Isatin | 19 | 16 | 2 |
| Retigabine | 19 | 17 | 83 |
| Cisapride hydrate | 19 | 18 | -173 |
| Menadiol Diacetate | 19 | 19 | -56 |
| Thymidine | 19 | 20 | -59 |
| Hydroquinidine | 19 | 21 | -108 |
| Lidocaine hydrochloride | 19 | 22 | -81 |
| Terazosin HCl | 19 | 23 | -145 |
| Escin | 19 | 24 | -249 |
| 2-Deoxy-D-glucose | 19 | 25 | -173 |

| | | | |
|---|---|---|---|
| Acetylcholine iodide | 19 | 26 | -12 |
| Retigabine 2HCl | 19 | 27 | 98 |
| Corticosterone | 19 | 28 | -5 |
| Benzyl isothiocyanate | 19 | 29 | -95 |
| Ceftizoxime | 19 | 30 | -72 |
| Glycopyrrolate | 19 | 31 | -103 |
| Procaine | 19 | 32 | -93 |
| Protirelin | 19 | 33 | -148 |
| Oxybenzone | 19 | 34 | 40 |
| Eugenol | 19 | 35 | 13 |
| (+)-Catechin | 19 | 36 | -108 |
| Salvianolic acid B | 19 | 37 | -49 |
| Betulin | 19 | 38 | -299 |
| N-Acetylneuraminic acid | 19 | 39 | -79 |
| Cefuroxime axetil | 19 | 40 | -50 |
| Tiagabine hydrochloride | 19 | 41 | 0 |
| Benzocaine hydrochloride | 19 | 42 | 30 |
| Loxoprofen | 19 | 43 | -36 |
| Guanfacine Hydrochloride | 19 | 44 | -155 |
| Oleic Acid | 19 | 45 | -62 |
| (-)Epicatechin | 19 | 46 | -79 |
| Trapidil | 19 | 47 | -54 |
| Dihydrotestosterone(DHT) | 19 | 48 | -148 |
| Drostanolone Propionate | 19 | 49 | 1 |
| L-Cysteine HCl | 19 | 50 | -50 |
| Atazanavir | 19 | 51 | 33 |
| Etonogestrel | 19 | 52 | 87 |
| Sildenafil Mesylate | 19 | 53 | 88 |
| D panthenol | 19 | 54 | 6 |
| Latanoprost | 19 | 55 | -307 |
| Benzenesulfonamide | 19 | 56 | 20 |
| Psoralen | 19 | 57 | -201 |
| p-Coumaric Acid | 19 | 58 | -19 |
| Trenbolone acetate | 19 | 59 | 77 |
| Diatrizoate sodium | 19 | 60 | -11 |
| Fusidate Sodium | 19 | 61 | 26 |
| Hydroxyprogesterone caproate | 19 | 62 | 68 |
| Efavirenz | 19 | 63 | 70 |
| CarbinoxaMine Maleate | 19 | 64 | -64 |
| Esculetin | 19 | 65 | -57 |
| Lauric Acid | 19 | 66 | -121 |
| Ondansetron Hydrochloride Dihydrate | 19 | 67 | 65 |
| Melibiose | 19 | 68 | -25 |
| Methandrostenolone | 19 | 69 | 33 |
| Atenolol | 19 | 70 | -62 |
| Molsidomine | 19 | 71 | -49 |
| Tiagabine | 19 | 72 | -116 |

| | | | |
|---|---|---|---|
| Vitamin E | 19 | 73 | 18 |
| Saxagliptin hydrate | 19 | 74 | -44 |
| (-)-Menthol | 19 | 75 | 83 |
| Cinnarizine | 19 | 76 | -90 |
| Citalopram HBr | 19 | 77 | 44 |
| L-5-Hydroxytryptophan | 19 | 78 | -20 |
| Nicergoline | 19 | 79 | 88 |
| Saccharin | 19 | 80 | 13 |
| Diastase | 20 | 1 | -143 |
| Ibudilast | 20 | 2 | 36 |
| Faropenem Sodium | 20 | 3 | -6 |
| Iproniazid | 20 | 4 | -19 |
| Sulfamethoxazole Sodium | 20 | 5 | -21 |
| Cefathiamidine | 20 | 6 | -79 |
| Cytosine | 20 | 7 | 7 |
| Maltol | 20 | 8 | -2 |
| Iminostilbene | 20 | 9 | -41 |
| Tiamulin Fumarate | 20 | 10 | 6 |
| Maltose | 20 | 11 | -4 |
| Acotiamide Hydrochloride | 20 | 12 | -9 |
| Dalbavancin | 20 | 13 | 82 |
| Triacetonamine | 20 | 14 | -3 |
| Cefodizime Sodium | 20 | 15 | -117 |
| Calcium Dobesilate | 20 | 16 | 7 |
| Elagolix Sodium | 20 | 17 | -46 |
| Nonanoic Acid | 20 | 18 | 5 |
| Fimasartan | 20 | 19 | 15 |
| Valpromide | 20 | 20 | 28 |
| Piperonyl Butoxide | 20 | 21 | 8 |
| Mosapride | 20 | 22 | 11 |
| Levocetirizine Dihydrochloride | 20 | 23 | 98 |
| Indole-3-Carboxylic Acid | 20 | 24 | -29 |
| Pyridoxal 5-Phosphate Monohydrate | 20 | 25 | -123 |
| Lynestrenol | 20 | 26 | 0 |
| Sulfogaiacol | 20 | 27 | -13 |
| Fumaric Acid | 20 | 28 | 21 |
| Sulfalene (SMPZ) | 20 | 29 | 7 |
| Methylcobalamin | 20 | 30 | -38 |
| Tolmetin | 20 | 31 | -16 |
| Laurocapram | 20 | 32 | -92 |
| Flucloxacillin Sodium | 20 | 33 | -69 |
| Squalene | 20 | 34 | 12 |
| Cefazedone | 20 | 35 | -65 |
| Taurolidine | 20 | 36 | 6 |
| Propiverine Hydrochloride | 20 | 37 | 4 |
| Usnic Acid | 20 | 38 | -110 |
| Efonidipine | 20 | 39 | 52 |

| | | | |
|---|---|---|---|
| Tavaborole | 20 | 40 | 11 |
| Cefoxitin Sodium | 20 | 41 | -79 |
| Potassium Acetate | 20 | 42 | -1 |
| Tafluprost | 20 | 43 | 77 |
| Cefetamet Pivoxil Hydrochloride | 20 | 44 | 94 |
| Cephaprin Benzathine | 20 | 45 | 70 |
| Menbutone | 20 | 46 | -14 |
| Proxyphylline | 20 | 47 | 11 |
| Linalool | 20 | 48 | 60 |
| Azathramycin | 20 | 49 | -5 |
| Avermectin B1 | 20 | 50 | 98 |
| Propantheline Bromide | 20 | 51 | -91 |
| Cefcapene Pivoxil Hydrochloride | 20 | 52 | 85 |
| Gadopentetate Dimeglumine | 20 | 53 | 13 |
| Nicarbazin | 20 | 54 | 0 |
| Robenidine Hydrochloride | 20 | 55 | 59 |
| Nikethamide | 20 | 56 | 14 |
| Lithium Carbonate | 20 | 57 | -10 |
| Glycocholic Acid | 20 | 58 | 30 |
| Anamorelin | 20 | 59 | 57 |
| Tofacitinib Citrate | 20 | 60 | 9 |
| Aceclofenac | 20 | 61 | 20 |
| Rabeprazole | 20 | 62 | 35 |
| Ecabet Sodium | 20 | 63 | 24 |
| Propacetamol Hydrochloride | 20 | 64 | 28 |
| Eperisone Hydrochloride | 20 | 65 | 29 |
| Perospirone Hydrochloride | 20 | 66 | 73 |
| Asunaprevir | 20 | 67 | -31 |
| Lactobionic Acid | 20 | 68 | -6 |
| Sorbic Acid | 20 | 69 | -58 |
| Fingolimod HCl | 20 | 70 | -116 |
| Nilutamide | 20 | 71 | 56 |
| Meropenem Trihydrate | 20 | 72 | -17 |
| Bedaquiline Fumarate | 20 | 73 | 72 |
| Xanthinol Nicotinate | 20 | 74 | -1 |
| Neticonazole Hydrochloride | 20 | 75 | -66 |
| Bifendate | 20 | 76 | 83 |
| cis-Aconitic Acid | 20 | 77 | -10 |
| Buparvaquone | 20 | 78 | -4 |
| Tacrolimus | 20 | 79 | 96 |
| Pimecrolimus | 21 | 1 | 86 |
| Ioversol | 21 | 2 | 29 |
| Efinaconazole | 21 | 3 | 83 |
| Phenazine methosulfate | 21 | 4 | 36 |
| Thiocolchicoside | 21 | 5 | 25 |
| Daclatasvir Dihydrochloride | 21 | 6 | 97 |
| Rosuvastatin | 21 | 7 | 74 |

| | | | |
|---|---|---|---|
| Ceforanide | 21 | 8 | -44 |
| Febantel | 21 | 9 | 66 |
| Lanolin | 21 | 10 | -34 |
| Cefotiam hydrochloride | 21 | 11 | -1 |
| Crisaborole (AN2728) | 21 | 12 | 31 |
| Mebeverine Hydrochloride | 21 | 13 | 71 |
| Valethamate Bromide | 21 | 14 | 22 |
| Granisetron | 21 | 15 | -31 |
| Trelagliptin succinate | 21 | 16 | -67 |
| Donepezil | 21 | 17 | 60 |
| Vitamin K2 | 21 | 18 | 7 |
| Rafoxanide | 21 | 19 | -72 |
| Tylosin | 21 | 20 | 48 |
| Teprenone | 21 | 21 | 33 |
| Simeprevir | 21 | 22 | -67 |
| 4-Aminopyridine | 21 | 23 | 0 |
| Actarit | 21 | 24 | 4 |
| Rifamycin sodium salt | 21 | 25 | -57 |
| Ganciclovir sodium | 21 | 26 | -9 |
| Argatroban Monohydrate | 21 | 27 | 54 |
| Lentinan | 21 | 28 | -72 |
| SulfadiMethoxine sodium | 21 | 29 | -152 |
| Ademetionine | 21 | 30 | -11 |
| Delamanid | 21 | 31 | -31 |
| Isoprinosine | 21 | 32 | -26 |
| Etofylline | 21 | 33 | -34 |
| Tiamulin | 21 | 34 | 31 |
| Milbemycin Oxime | 21 | 35 | 89 |
| Pramipexole dihydrochloride | 21 | 36 | -98 |
| Acotiamide | 21 | 37 | -27 |
| Carbazochrome | 21 | 38 | -38 |
| alpha-Arbutin | 21 | 39 | 25 |
| Brivudine | 21 | 40 | 17 |
| Oxyclozanide | 21 | 41 | -52 |
| Dihydralazine sulphate | 21 | 42 | -33 |
| Difloxacin hydrochloride | 21 | 43 | -3 |
| Dinoprost tromethamine | 21 | 44 | 65 |
| Losartan | 21 | 45 | -76 |
| Xipamide | 21 | 46 | -26 |
| Azamethiphos | 21 | 47 | -42 |
| Pralidoxime Iodide | 21 | 48 | 5 |
| Propyl gallate | 21 | 49 | -3 |
| Indometacin Sodium | 21 | 50 | -2 |
| Indobufen | 21 | 51 | 12 |
| Mephenesin | 21 | 52 | -32 |
| Bevantolol hydrochloride | 21 | 53 | 30 |
| Revaprazan Hydrochloride | 21 | 54 | 51 |

| | | | |
|---|---|---|---|
| Dabrafenib Mesylate | 21 | 55 | 4 |
| Regorafenib Monohydrate | 21 | 56 | 10 |
| p-Anisaldehyde | 21 | 57 | 14 |
| Stachyose | 21 | 58 | -33 |
| Hydroquinine | 21 | 59 | -22 |
| Disodium Phosphate | 21 | 60 | 44 |
| Tilorone dihydrochloride | 21 | 61 | 16 |
| Terconazole | 21 | 62 | 76 |
| Benorylate | 21 | 63 | 49 |
| Pixantrone Maleate | 21 | 64 | -172 |
| Mupirocin calcium | 21 | 65 | 25 |
| Osimertinib mesylate | 21 | 66 | -289 |
| Tianeptine | 21 | 67 | 12 |
| lutein | 21 | 68 | 37 |
| Doramectin | 21 | 69 | 96 |
| Octenidine Dihydrochloride | 21 | 70 | 100 |
| Nadolol | 21 | 71 | -6 |
| Melitracen hydrochloride | 21 | 72 | 48 |
| Clonixin | 21 | 73 | 10 |
| Metadoxine | 21 | 74 | -2 |
| Duloxetine | 21 | 75 | 25 |
| Sitagliptin | 21 | 76 | 36 |
| Geranyl acetate | 21 | 77 | 52 |
| Proanthocyanidins | 21 | 78 | 78 |
| Olivetol | 21 | 79 | 72 |
| Phytol | 22 | 1 | -12 |
| Raffinose | 22 | 2 | -26 |
| Ropivacaine Mesilate | 22 | 3 | -17 |
| 4-Aminosalicylic Acid | 22 | 4 | -35 |
| Boldenone | 22 | 5 | 21 |
| Lenvatinib Mesylate | 22 | 6 | 94 |
| Cisapride | 22 | 7 | -38 |
| Canrenone | 22 | 8 | 12 |
| Estropipate | 22 | 9 | -52 |
| Tosufloxacin (P-Toluenesulfonic Acid) | 22 | 10 | 97 |
| Aleuritic Acid | 22 | 11 | -73 |
| Thymol | 22 | 12 | 22 |
| Diaveridine | 22 | 13 | -34 |
| Piroctone Olamine | 22 | 14 | 3 |
| Erythromycin Thiocyanate | 22 | 15 | 5 |
| Lapatinib Ditosylate Monohydrate | 22 | 16 | 79 |
| Dasatinib Hydrochloride | 22 | 17 | 98 |
| Tedizolid | 22 | 18 | -29 |
| Ceftezole | 22 | 19 | 12 |
| Carglumic Acid | 22 | 20 | 3 |
| Casanthranol | 22 | 21 | -75 |
| Doxycycline | 22 | 22 | -34 |

| | | | |
|---|---|---|---|
| Sulbutiamine | 22 | 23 | 13 |
| Imidocarb Dipropionate | 22 | 24 | -289 |
| Minoxidil Sulphate | 22 | 25 | 11 |
| Ruxolitinib Phosphate | 22 | 26 | 25 |
| Nicardipine | 22 | 27 | -9 |
| Fendiline Hydrochloride | 22 | 28 | -100 |
| Sulfaisodimidine | 22 | 29 | 5 |
| 6-Methoxy-2-Naphthoic Acid | 22 | 30 | 17 |
| D(-)-Arabinose | 22 | 31 | 1 |
| 7-Methoxycoumarin | 22 | 32 | 25 |
| Nilotinib Hydrochloride | 22 | 33 | 99 |
| Abacavir | 22 | 34 | 16 |
| Isosorbide Dinitrate | 22 | 35 | 7 |
| Raltegravir Potassium | 22 | 36 | 39 |
| Benproperine Phosphate | 22 | 37 | -98 |
| Dapiprazole Hydrochloride | 22 | 38 | -64 |
| Diazolidinyl Urea | 22 | 39 | 7 |
| Bromisoval | 22 | 40 | 20 |
| Citral | 22 | 41 | 20 |
| Benzyl Cinnamate | 22 | 42 | 9 |
| Benzylpenicillin Potassium | 22 | 43 | 19 |
| Arformoterol Tartrate | 22 | 44 | -99 |
| Fenretinide | 22 | 45 | 54 |
| Entecavir | 22 | 46 | 28 |
| Doxapram | 22 | 47 | 5 |
| Lactitol Monohydrate | 22 | 48 | 11 |
| Nimustine Hydrochloride | 22 | 49 | 8 |
| Propyphenazone | 22 | 50 | 13 |
| Pyrithioxin | 22 | 51 | 75 |
| Kojic Acid | 22 | 52 | -12 |
| Ciprofloxacin Hydrochloride Hydrate | 22 | 53 | 93 |
| Iodixanol | 22 | 54 | 23 |
| Nintedanib Ethanesulfonate Salt | 22 | 55 | 87 |
| Selamectin | 22 | 56 | 97 |
| Revefenacin | 22 | 57 | 25 |
| Adenosine 5-Monophosphate | 22 | 58 | 26 |
| Lincomycin Hydrochloride Monohydra | 22 | 59 | 13 |
| Pyridoxal Phosphate | 22 | 60 | -36 |
| Citropten | 22 | 61 | 41 |
| Fructose | 22 | 62 | 21 |
| Enoxacin Sesquihydrate | 22 | 63 | 31 |
| Phenylpiracetam | 22 | 64 | 36 |
| Solifenacin | 22 | 65 | -46 |
| Apatinib | 22 | 66 | 44 |
| Ufenamate | 22 | 67 | 27 |
| Ramatroban | 22 | 68 | -47 |
| Vidarabine Monohydrate | 22 | 69 | -12 |

| | | | |
|---|---|---|---|
| Octinoxate | 22 | 70 | 29 |
| Protoporphyrin IX | 22 | 71 | 33 |
| Naproxen | 22 | 72 | 27 |
| Sulfamethazine Sodium Salt | 22 | 73 | 34 |
| Adrafinil | 22 | 74 | 43 |
| Paroxetine Mesylate | 22 | 75 | -55 |
| Darunavir | 22 | 76 | 53 |
| Stiripentol | 22 | 77 | 63 |
| Tiletamine Hydrochloride | 22 | 78 | 16 |
| Amoxicillin Trihydrate | 22 | 79 | -18 |
| Sodium Gualenate | 22 | 80 | -78 |
| Nitisinone | 23 | 1 | -60 |
| Nerolidol | 23 | 2 | -24 |
| Butoconazole | 23 | 3 | 81 |
| Ethoxyquin | 23 | 4 | 46 |
| Betrixaban maleate | 23 | 5 | -174 |
| Dasabuvir(ABT-333) | 23 | 6 | 88 |
| Sultamicillin | 23 | 7 | -64 |
| Carbaryl | 23 | 8 | 44 |
| Hyperoside | 23 | 9 | 18 |
| Dantrolene sodium | 23 | 10 | -45 |
| Dolasetron | 23 | 11 | 64 |
| Cefpodoxime proxetil | 23 | 12 | 94 |
| Diflorasone | 23 | 13 | 19 |
| Ajmaline | 23 | 14 | -85 |
| Mepivacaine | 23 | 15 | -13 |
| Ombitasvir (ABT-267) | 23 | 16 | 92 |
| Ertugliflozin | 23 | 17 | 24 |
| Promazine hydrochloride | 23 | 18 | 38 |
| Saikosaponin D | 23 | 19 | 83 |
| Cloperastine hydrochloride | 23 | 20 | 73 |
| Meisoindigo | 23 | 21 | -2 |
| Cefmetazole sodium | 23 | 22 | -5 |
| Bendazac | 23 | 23 | 41 |
| Methyl Aminolevulinate Hydrochloride | 23 | 24 | 0 |
| Cyclofenil | 23 | 25 | 64 |
| Paritaprevir (ABT-450) | 23 | 26 | -24 |
| Diflucortolone valerate | 23 | 27 | 74 |
| Metoprolol | 23 | 28 | -16 |
| Curculigoside | 23 | 29 | 58 |
| Clidinium Bromide | 23 | 30 | -43 |
| Gamithromycin | 23 | 31 | 50 |
| Cefminox Sodium | 23 | 32 | -7 |
| Pikamilone | 23 | 33 | 36 |
| Dibutyl phthalate | 23 | 34 | -15 |
| Phenolphthalein | 23 | 35 | 57 |
| Propylparaben | 23 | 36 | 53 |

| | | | |
|---|---|---|---|
| Enoxaparin sodium | 23 | 37 | -59 |
| Quinacrine Dihydrochloride Dihydrate | 23 | 38 | -55 |
| Aucubin | 23 | 39 | 24 |
| Molindone hydrochloride | 23 | 40 | 6 |
| Ceftezole sodium | 23 | 41 | -33 |
| Cefpiramide sodium | 23 | 42 | -58 |
| Alogliptin | 23 | 43 | -7 |
| Dimethyl phthalate | 23 | 44 | 10 |
| Chlorhexidine | 23 | 45 | 37 |
| Sultamicillin Tosylate | 23 | 46 | 42 |
| Metyrapone | 23 | 47 | 1 |
| Berberine Sulfate | 23 | 48 | -73 |
| Saikosaponin A | 23 | 49 | -226 |
| Prilocaine hydrochloride | 23 | 50 | -23 |
| Sulbenicillin Sodium | 23 | 51 | 11 |
| Ceftiofur | 23 | 52 | -1 |
| Fipronil | 23 | 53 | 91 |
| Formate | 23 | 54 | 18 |
| Nefazodone hydrochloride | 23 | 55 | 85 |
| Squalane | 23 | 56 | 5 |
| Parecoxib Sodium | 23 | 57 | 29 |
| Triprolidine Hydrochloride | 23 | 58 | 28 |
| Pivmecillinam hydrochloride | 23 | 59 | 34 |
| Tribenzagan Hydrochloride | 23 | 60 | -57 |
| Metroprolol succinate | 23 | 61 | -61 |
| Safinamide | 23 | 62 | 45 |
| Ethyl Oleate | 23 | 63 | 30 |
| Imidafenacin | 23 | 64 | 1 |
| Chlorprothixene hydrochloride | 23 | 65 | -171 |
| Isoprene | 23 | 66 | -10 |
| 1,4-Cineole | 23 | 67 | 20 |
| Sofalcone | 23 | 68 | -101 |
| Rolapitant | 23 | 69 | 86 |
| Rimantadine Hydrochloride | 23 | 70 | 14 |
| Vanillic acid | 23 | 71 | -11 |
| Regadenoson | 23 | 72 | 0 |
| Lactitol | 23 | 73 | 9 |
| Betrixaban | 23 | 74 | -11 |
| Tegaserod Maleate | 23 | 75 | -118 |
| Chloramphenicol sodium succinate | 23 | 76 | -68 |
| Clindamycin alcoholate | 23 | 77 | 12 |
| Sanguinarine chloride | 23 | 78 | -25 |
| Gefarnate | 23 | 79 | -2 |
| Desipramine Hydrochloride | 23 | 80 | 24 |
| Fluorometholone | 24 | 1 | 89 |
| Iopanoic acid | 24 | 2 | 71 |
| Trimebutine maleate | 24 | 3 | 88 |

| | | | |
|---|---|---|---|
| Erythromycin estolate | 24 | 4 | 94 |
| 1, 10-Phenanthroline monohydrate | 24 | 5 | 48 |
| Pamabrom | 24 | 6 | 41 |
| Xylazine | 24 | 7 | 36 |
| Olmesartan | 24 | 8 | 21 |
| (1R)-(-)-Menthyl acetate | 24 | 9 | 85 |
| Bedaquiline | 24 | 10 | 58 |
| Cefoperazone sodium | 24 | 11 | 53 |
| Betahistine mesylate | 24 | 12 | 53 |
| Dehydroepiandrosterone acetate | 24 | 13 | 92 |
| Cinchocaine | 24 | 14 | 94 |
| D-Ribose | 24 | 15 | 52 |
| (-)-Sparteine Sulfate | 24 | 16 | 12 |
| Triflupromazine hydrochloride | 24 | 17 | 38 |
| Cytarabine hydrochloride | 24 | 18 | 49 |
| p-Cymene | 24 | 19 | 43 |
| Ammonium lactate | 24 | 20 | 30 |
| Fluorescein | 24 | 21 | 42 |
| Amodiaquine hydrochloride | 24 | 22 | 90 |
| 4-Aminophenol | 24 | 23 | 60 |
| Moxifloxacin | 24 | 24 | 36 |
| Sulfacetamide | 24 | 25 | 47 |
| D-Pantothenate Sodium | 24 | 26 | 38 |
| Dapagliflozin propanediol monohydrat | 24 | 27 | 93 |
| Cinnamyl acetate | 24 | 28 | 89 |
| Sodium cholate | 24 | 29 | 25 |
| Benzalkonium chloride | 24 | 30 | -38 |
| Disopyramide | 24 | 31 | 40 |
| Hydrocortisone acetate | 24 | 32 | 56 |
| Penicillin G Procaine | 24 | 33 | 26 |
| Tizanidine | 24 | 34 | 54 |
| Hydroxylammonium chloride | 24 | 35 | 29 |
| Tetrahydropalmatine | 24 | 36 | 90 |
| Trimethadione | 24 | 37 | 37 |
| Citronellal | 24 | 38 | 68 |
| Diphenylamine Hydrochloride | 24 | 39 | 92 |
| Amsacrine hydrochloride | 24 | 40 | 98 |
| Lomefloxacin | 24 | 41 | 45 |
| Ilaprazole sodium | 24 | 42 | 94 |
| Salmeterol | 24 | 43 | 90 |
| Tropisetron | 24 | 44 | 41 |
| Ethyl gallate | 24 | 45 | 58 |
| Midecamycin | 24 | 46 | 93 |
| Tocofersolan | 24 | 47 | -57 |
| Camphene | 24 | 48 | 57 |
| (+)-Longifolene | 24 | 49 | 77 |
| Cefotiam Hexetil Hydrochloride | 24 | 50 | 99 |

| | | | |
|---|---|---|---|
| Econazole | 24 | 51 | 82 |
| Ropivacaine | 24 | 52 | 66 |
| Acetophenone | 24 | 53 | 64 |
| Olprinone | 24 | 54 | 49 |
| Amenamevir | 24 | 55 | 88 |
| Ethacrynic Acid | 24 | 56 | 59 |
| Anisole | 24 | 57 | 58 |
| Vitamin A | 24 | 58 | 77 |
| Hippuric acid | 24 | 59 | 38 |
| Cefozopran hydrochloride | 24 | 60 | 53 |
| Atropine sulfate | 24 | 61 | 54 |
| 2’-deoxyuridine | 24 | 62 | 69 |
| Geraniol | 24 | 63 | 96 |
| Landiolol hydrochloride | 24 | 64 | 49 |
| Kasugamycin hydrochloride | 24 | 65 | 42 |
| 2-Hydroxybenzyl alcohol | 24 | 66 | 63 |
| Tilmicosin phosphate | 24 | 67 | 72 |
| α-Terpineol | 24 | 68 | 87 |
| Betahistine | 24 | 69 | 70 |
| Ceftriaxone Sodium | 24 | 70 | 36 |
| Salbutamol | 24 | 71 | 39 |
| Vortioxetine | 24 | 72 | -18 |
| Doripenem | 24 | 73 | 38 |
| Dimetridazole | 24 | 74 | 40 |
| Lanatoside C | 24 | 75 | 80 |
| Thioridazine hydrochloride | 24 | 76 | 62 |
| Arabic gum | 24 | 77 | 18 |
| (1S)-(-)-α-Pinene | 24 | 78 | 58 |
| Cilastatin | 24 | 79 | 47 |
| Emedastine Difumarate | 24 | 80 | 53 |
| Iguratimod | 25 | 1 | 46 |
| Diphenylpyraline Hydrochloride | 25 | 2 | -89 |
| Drofenine Hydrochloride | 25 | 3 | -109 |
| Methyl Linolenate | 25 | 4 | 23 |
| Phthalylsulfathiazole | 25 | 5 | 32 |
| Pravastatin | 25 | 6 | 39 |
| Oxantel Pamoate | 25 | 7 | -47 |
| Stearic Acid | 25 | 8 | -34 |
| Deferoxamine Mesylate | 25 | 9 | 31 |
| Citric Acid | 25 | 10 | 27 |
| Hydroxyzine Pamoate | 25 | 11 | -49 |
| Ertapenem Sodium | 25 | 12 | -99 |
| Moxisylyte Hydrochloride | 25 | 13 | -34 |
| Isoproterenol Sulfate Dihydrate | 25 | 14 | 29 |
| Alvimopan Dihydrate | 25 | 15 | 37 |
| Lurasidone | 25 | 16 | 25 |
| Dipchlorophene | 25 | 17 | 28 |

| | | | |
|---|---|---|---|
| Midodrine Hydrochloride | 25 | 18 | 16 |
| Morantel Tartrate | 25 | 19 | -15 |
| Methyl Oleate | 25 | 20 | 31 |
| Omeprazole Sodium | 25 | 21 | 41 |
| Fruquintinib | 25 | 22 | 54 |
| Desoximetasone | 25 | 23 | 39 |
| Fenipentol | 25 | 24 | 41 |
| Atorvastatin | 25 | 25 | -1 |
| Triiodothyronine | 25 | 26 | 8 |
| Midodrine | 25 | 27 | 32 |
| Chlorpromazine | 25 | 28 | -100 |
| D-Mannose | 25 | 29 | 38 |
| Dexrazoxane | 25 | 30 | 26 |
| Emedastine | 25 | 31 | -85 |
| Isoxuprine Hydrochloride | 25 | 32 | -9 |
| Tropic Acid | 25 | 33 | 28 |
| Carvedilol Phosphate | 25 | 34 | 58 |
| Abemaciclib | 25 | 35 | -208 |
| Tetryzoline | 25 | 36 | 27 |
| Benzathine Penicilline | 25 | 37 | -54 |
| Dihydroergotamine Mesylate | 25 | 38 | 94 |
| Sodium Dehydrocholate | 25 | 39 | -17 |
| Elbasvir | 25 | 40 | 100 |
| Tiaprofenic Acid | 25 | 41 | 38 |
| Chloropyramine Hydrochloride | 25 | 42 | -101 |
| Mexenone | 25 | 43 | 49 |
| Raceanisodamine | 25 | 44 | 39 |
| Acetohexamide | 25 | 45 | 48 |
| Delapril Hydrochloride | 25 | 46 | 37 |
| Diclofenac Epolamine | 25 | 47 | 49 |
| Baricitinib Phosphate | 25 | 48 | 42 |
| Alfuzosin | 25 | 49 | -30 |
| Indigo Carmine | 25 | 50 | -85 |
| Ranitidine | 25 | 51 | -34 |
| Mivacurium Chloride | 25 | 52 | -89 |
| Levomilnacipran Hydrochloride | 25 | 53 | -9 |
| Norgestrel | 25 | 54 | 70 |
| Acrivastine | 25 | 55 | -13 |
| Fofosal | 25 | 56 | 77 |
| Nevibolol | 25 | 57 | -62 |
| Methyl Stearate | 25 | 58 | 43 |
| Aliskiren | 25 | 59 | 10 |
| Indacaterol | 25 | 60 | 16 |
| Minaprine Dihydrochloride | 25 | 61 | 17 |
| Dolasetron Mesylate | 25 | 62 | 20 |
| Isopropamide Iodide | 25 | 63 | 30 |
| Ambroxol | 25 | 64 | -5 |

| | | | |
|---|---|---|---|
| Ceftizoxime Sodium | 25 | 65 | -10 |
| Alimemazine Tartrate | 25 | 66 | -96 |
| Palonosetron | 25 | 67 | -30 |
| Isoeugenol | 25 | 68 | 37 |
| Fenoterol Hydrobromide | 25 | 69 | 10 |
| Venlafaxine | 25 | 70 | 4 |
| Orphenadrine Hydrochloride | 25 | 71 | -80 |
| (-)-Verbenone | 25 | 72 | 38 |
| Ketorolac Tromethamine Salt | 25 | 73 | 42 |
| Glecaprevir | 25 | 74 | -45 |
| Sebacic Acid | 25 | 75 | 30 |
| Quetiapine | 25 | 76 | -15 |
| Methyl Linoleate | 25 | 77 | 55 |
| Fenoterol | 25 | 78 | 14 |
| Proflavine | 26 | 1 | -136 |
| Dronedarone | 26 | 2 | -131 |
| Citronellyl acetate | 26 | 3 | -4 |
| Cortisone | 26 | 4 | -2 |
| Bictegravir | 26 | 5 | -32 |
| Ataluren (PTC124) | 26 | 6 | -26 |
| Dibutyl sebacate | 26 | 7 | -13 |
| Doxycycline monohydrate | 26 | 8 | -83 |
| p-Toluenesulfonic acid monohydrate | 26 | 9 | -34 |
| Phthalic acid | 26 | 10 | -24 |
| Esmolol | 26 | 11 | -60 |
| Rasagiline | 26 | 12 | -28 |
| Cabergoline | 26 | 13 | -56 |
| Trans-Tranilast | 26 | 14 | 15 |
| Celiprolol hydrochloride | 26 | 15 | -181 |
| Imidazole | 26 | 16 | -6 |
| Lactose | 26 | 17 | -8 |
| Diclofenac acid | 26 | 18 | -16 |
| 4-Nitrophenol | 26 | 19 | -12 |
| o-Toluic acid | 26 | 20 | -26 |
| Trimetazidine | 26 | 21 | -18 |
| Alprenolol hydrochloride | 26 | 22 | -34 |
| Cinitapride Hydrogen Tartrate | 26 | 23 | 76 |
| Estradiol dipropionate(17-Beta-Estradi | 26 | 24 | 61 |
| Olanexidine Hydrochloride semihydrat | 26 | 25 | 95 |
| Bisphenol A | 26 | 26 | 63 |
| (+)-(S)-Carvone | 26 | 27 | -3 |
| 2-Methylhexanoic acid | 26 | 28 | 25 |
| Dimethylamine hydrochloride | 26 | 29 | -3 |
| Sodium lauryl sulfate | 26 | 30 | -4 |
| Prazosin | 26 | 31 | -113 |
| Allopregnanolone | 26 | 32 | 71 |
| Vilazodone | 26 | 33 | 85 |

| | | | |
|---|---|---|---|
| Scopolamine HBr trihydrate | 26 | 34 | 11 |
| Olodaterol hydrochloride | 26 | 35 | 1 |
| Sodium L-lactate | 26 | 36 | -2 |
| Saccharin sodium salt hydrate | 26 | 37 | 33 |
| (±)-α-Tocopherol | 26 | 38 | -20 |
| o-Cresol | 26 | 39 | 17 |
| Methyl cinnamate | 26 | 40 | 1 |
| Raloxifene | 26 | 41 | 85 |
| Relugolix | 26 | 42 | -10 |
| Lercanidipine | 26 | 43 | 2 |
| L-Carnitine hydrochloride | 26 | 44 | -10 |
| Pitolisant hydrochloride | 26 | 45 | -5 |
| Isonicotinic acid | 26 | 46 | -4 |
| Phenylglyoxylic acid | 26 | 47 | -21 |
| Terpinen-4-ol | 26 | 48 | 6 |
| Butylated hydroxytoluene | 26 | 49 | -45 |
| Triethyl citrate | 26 | 50 | -13 |
| Doxazosin | 26 | 51 | 46 |
| Choline Fenofibrate | 26 | 52 | 6 |
| Metoclopramide | 26 | 53 | -125 |
| Edrophonium chloride | 26 | 54 | -99 |
| Proguanil | 26 | 55 | 7 |
| p-Cresol | 26 | 56 | 44 |
| (S)-(−)-Limonene | 26 | 57 | 21 |
| Maltotriose | 26 | 58 | 19 |
| Levulinic acid | 26 | 59 | 23 |
| 4-Methyl-2-pentanone | 26 | 60 | 2 |
| Montelukast | 26 | 61 | 22 |
| pyrvinium | 26 | 62 | -135 |
| Metronidazole Benzoate | 26 | 63 | -20 |
| Canagliflozin hemihydrate | 26 | 64 | 14 |
| Alvimopan | 26 | 65 | 17 |
| p-Benzoquinone | 26 | 66 | -58 |
| β-Caryophyllene | 26 | 67 | -23 |
| Brucine sulfate heptahydrate | 26 | 68 | -101 |
| Ethanolamine hydrochloride | 26 | 69 | -21 |
| Methyl nicotinate | 26 | 70 | -10 |
| Vancomycin | 26 | 71 | -57 |
| Linoleic acid | 26 | 72 | -237 |
| Anagliptin | 26 | 73 | -25 |
| Fingolimod | 26 | 74 | -79 |
| 2-Naphthol | 26 | 75 | 24 |
| 2,4-dichlorobenzyl alcohol | 26 | 76 | 21 |
| Amylmetacresol | 26 | 77 | 74 |
| Furfural | 26 | 78 | 6 |
| Tartaric acid | 26 | 79 | -11 |
| Pentadecanoic acid | 27 | 1 | -449 |

| | | | |
|---|---|---|---|
| Rutin hydrate | 27 | 2 | -167 |
| N-Acetyl-L-tyrosine | 27 | 3 | -96 |
| Duvelisib (IPI-145, INK1197) | 27 | 4 | -3 |
| BAF312 (Siponimod) | 27 | 5 | -386 |
| Triapine | 27 | 6 | 79 |
| Niraparib (MK-4827) tosylate | 27 | 7 | -202 |
| Monomethyl auristatin E (MMAE) | 27 | 8 | 47 |
| Afatinib (BIBW2992) Dimaleate | 27 | 9 | -56 |
| CP21R7 (CP21) | 27 | 10 | -92 |
| Sodium dehydroacetate | 27 | 11 | -92 |
| Glycolic acid | 27 | 12 | -57 |
| 5-Methoxytryptamine | 27 | 13 | 12 |
| Tezacaftor?(VX-661) | 27 | 14 | 17 |
| Edoxaban | 27 | 15 | 95 |
| (S)-crizotinib | 27 | 16 | 26 |
| Lomitapide Mesylate | 27 | 17 | 93 |
| Gilteritinib (ASP2215) | 27 | 18 | -89 |
| Cyclo (-RGDfK) | 27 | 19 | -68 |
| Larotrectinib (LOXO-101) sulfate | 27 | 20 | 93 |
| Pyrrolidine | 27 | 21 | -66 |
| (−)-β-Pinene | 27 | 22 | -40 |
| m-Cresol | 27 | 23 | -21 |
| Cilengitide?trifluoroacetate | 27 | 24 | -132 |
| Osimertinib (AZD9291) | 27 | 25 | -1046 |
| Trelagliptin | 27 | 26 | -110 |
| Lomitapide | 27 | 27 | 78 |
| Sunitinib | 27 | 28 | 30 |
| Cyclo(RGDyK) | 27 | 29 | -169 |
| Favipiravir (T-705) | 27 | 30 | -3 |
| L-Lactic acid | 27 | 31 | -140 |
| Sodium Thiocyanate | 27 | 32 | -54 |
| Nonadecanoic acid | 27 | 33 | -341 |
| Ceritinib (LDK378) | 27 | 34 | -94 |
| Rilpivirine | 27 | 35 | 2 |
| Lorlatinib?(PF-6463922) | 27 | 36 | -26 |
| Peficitinib (ASP015K, JNJ-54781532) | 27 | 37 | 19 |
| Dasatinib Monohydrate | 27 | 38 | 95 |
| Eliglustat | 27 | 39 | 78 |
| Ripasudil (K-115) hydrochloride dihydr | 27 | 40 | -47 |
| Terephthalic acid | 27 | 41 | 12 |
| Tetraethylammonium bromide | 27 | 42 | -35 |
| D-Glucuronic acid | 27 | 43 | -124 |
| Zotarolimus(ABT-578) | 27 | 44 | 96 |
| Sorafenib | 27 | 45 | 41 |
| Erythromycin Cyclocarbonate | 27 | 46 | 83 |
| Obeticholic Acid | 27 | 47 | 24 |
| Combretastatin A4 | 27 | 48 | 89 |

| | | | |
|---|---|---|---|
| Tenofovir Alafenamide (GS-7340) | 27 | 49 | 60 |
| Vonoprazan Fumarate (TAK-438) | 27 | 50 | -56 |
| N-Acetylglucosamine | 27 | 51 | 48 |
| Chlorhexidine diacetate | 27 | 52 | 11 |
| L-(+)-Arabinose | 27 | 53 | -27 |
| Marimastat (BB-2516) | 27 | 54 | -4 |
| Ascomycin (FK520) | 27 | 55 | 55 |
| Darolutamide (ODM-201) | 27 | 56 | 5 |
| Ruboxistaurin (LY333531 HCl) | 27 | 57 | 74 |
| Fumagillin | 27 | 58 | -36 |
| Dibutyryl-cAMP (Bucladesine) | 27 | 59 | -26 |
| Vortioxetine (Lu AA21004) HBr | 27 | 60 | -107 |
| 1,2-Propanediol | 27 | 61 | -112 |
| Octanoic acid | 27 | 62 | -16 |
| Androsterone | 27 | 63 | -81 |
| abemaciclib (LY2835219) | 27 | 64 | -91 |
| Puromycin 2HCl | 27 | 65 | -135 |
| Ledipasvir (GS5885) | 27 | 66 | 89 |
| Picropodophyllin (PPP) | 27 | 67 | 98 |
| Erlotinib | 27 | 68 | -17 |
| Oleuropein | 27 | 69 | -86 |
| Empagliflozin (BI 10773) | 27 | 70 | -8 |
| Ammonium formate | 27 | 71 | -133 |
| Catechol | 27 | 72 | -174 |
| Binimetinib (MEK162, ARRY-162, ARRY | 27 | 73 | -29 |
| Glasdegib (PF-04449913) | 27 | 74 | -99 |
| Ribociclib (LEE011) | 27 | 75 | -61 |
| Abscisic Acid (Dormin） | 27 | 76 | -161 |
| Sacubitril/valsartan (LCZ696) | 27 | 77 | -38 |
| Docetaxel Trihydrate | 27 | 78 | 95 |
| Y-39983 HCl | 27 | 79 | -128 |
| Apremilast (CC-10004) | 27 | 80 | 96 |
| Cobimetinib | 28 | 1 | -93 |
| Riociguat | 28 | 2 | 45 |
| Melphalan | 28 | 3 | 0 |
| Tirofiban Hydrochloride | 28 | 4 | 37 |
| Sanguinarine | 28 | 5 | -302 |
| Aristolochic Acid A | 28 | 6 | 14 |
| Topotecan | 28 | 7 | -6 |
| Oxalic Acid | 28 | 8 | 0 |
| Yangonin | 28 | 9 | -102 |
| Isocarboxazid | 28 | 10 | -16 |
| Venetoclax | 28 | 11 | -24 |
| Sivelestat | 28 | 12 | 37 |
| Olmutinib | 28 | 13 | 98 |
| Sodium Dichloroacetate | 28 | 14 | 24 |
| Wedelolactone | 28 | 15 | 32 |

| | | | |
|---|---|---|---|
| Melamine | 28 | 16 | 48 |
| Scopolamine | 28 | 17 | 44 |
| ADP | 28 | 18 | 31 |
| Uridine 5'-Monophosphate | 28 | 19 | 31 |
| Demecarium Bromide | 28 | 20 | 10 |
| Macitentan | 28 | 21 | -12 |
| Halofuginone | 28 | 22 | 9 |
| Erdafitinib | 28 | 23 | -7 |
| Ipragliflozin | 28 | 24 | 44 |
| Berberine | 28 | 25 | -53 |
| Dipotassium Glycyrrhizinate | 28 | 26 | 30 |
| Carboprost | 28 | 27 | -58 |
| Neryl Acetate | 28 | 28 | 1 |
| Undecanoic Acid | 28 | 29 | -41 |
| Methysergide Maleate | 28 | 30 | 54 |
| Vorapaxar | 28 | 31 | 9 |
| Mitomycin C | 28 | 32 | 33 |
| Troglitazone | 28 | 33 | 63 |
| Anlotinib Dihydrochloride | 28 | 34 | 92 |
| Harringtonine | 28 | 35 | 47 |
| Sinensetin | 28 | 36 | 38 |
| Orcinol | 28 | 37 | 43 |
| 5,7-dihydroxy-4-methylcoumarin | 28 | 38 | -9 |
| Phensuximide | 28 | 39 | 10 |
| Methenamine Hippurate | 28 | 40 | 9 |
| CB-5083 | 28 | 41 | 72 |
| Pimavanserin | 28 | 42 | -192 |
| TAS-102 | 28 | 43 | 34 |
| Malic Acid | 28 | 44 | 47 |
| Pulegone | 28 | 45 | 56 |
| Isofraxidin | 28 | 46 | 30 |
| Octyl Gallate | 28 | 47 | -41 |
| Thymine | 28 | 48 | 47 |
| Ergoloid Mesylates | 28 | 49 | 98 |
| Thiothixene | 28 | 50 | 39 |
| Acalabrutinib | 28 | 51 | 91 |
| Oclacitinib Maleate | 28 | 52 | 44 |
| Tofogliflozin | 28 | 53 | 63 |
| L-Fucose | 28 | 54 | 29 |
| Berbamine | 28 | 55 | 85 |
| Securinine | 28 | 56 | 31 |
| 1-Indanone | 28 | 57 | 47 |
| Methyl Palmitate | 28 | 58 | 41 |
| Mecamylamine Hydrochloride | 28 | 59 | 50 |
| Haloperidol Decanoate | 28 | 60 | 65 |
| Resiquimod | 28 | 61 | 44 |
| Enasidenib | 28 | 62 | 42 |

| | | | |
|---|---|---|---|
| Omariglipton | 28 | 63 | 86 |
| R-(-)-Mandelic Acid | 28 | 64 | 31 |
| Sparteine | 28 | 65 | -10 |
| 3-n-Butylphathlide | 28 | 66 | -18 |
| D-(+)-Raffinose Pentahydrate | 28 | 67 | 25 |
| Ligustilide | 28 | 68 | -20 |
| Ethotoin | 28 | 69 | -7 |
| Penbutolol Sulfate | 28 | 70 | -30 |
| Radotinib | 28 | 71 | 93 |
| Ivosidenib | 28 | 72 | 62 |
| Tucidinostat | 28 | 73 | 62 |
| 2-Deoxyguanosine | 28 | 74 | 20 |
| Ammonium Glycyrrhizate | 28 | 75 | 15 |
| Germacrone | 28 | 76 | -42 |
| 2-Deoxyadenosine | 28 | 77 | 49 |
| Rhynchophylline | 28 | 78 | 39 |
| Benzonatate | 28 | 79 | 39 |
| Oxtriphylline | 28 | 80 | 17 |
| Valbenazine tosylate | 29 | 1 | 31 |
| Farrerol | 29 | 2 | 46 |
| Quillaic acid | 29 | 3 | 46 |
| Isorhamnetin | 29 | 4 | -32 |
| Dracohodin perochlorate | 29 | 5 | 77 |
| Clofibrate | 29 | 6 | -6 |
| levalbuterol tartrate | 29 | 7 | -48 |
| Benzamidine HCl | 29 | 8 | -33 |
| Khellin | 29 | 9 | 18 |
| Butenafine | 29 | 10 | 48 |
| Madecassic acid | 29 | 11 | 45 |
| Vincristine | 29 | 12 | 81 |
| AKBA | 29 | 13 | 68 |
| Eriodictyol | 29 | 14 | 28 |
| Peimine | 29 | 15 | 27 |
| Seratrodast(AA-2414, ABT-001) | 29 | 16 | 64 |
| Antazoline HCl | 29 | 17 | 21 |
| Ethacrynate Sodium | 29 | 18 | 56 |
| Cyclopentolate Hydrochloride | 29 | 19 | 52 |
| Amodiaquine | 29 | 20 | 63 |
| PA-824 | 29 | 21 | -75 |
| Anisodamine Hydrobromide | 29 | 22 | -29 |
| Cimifugin | 29 | 23 | 43 |
| Harpagoside | 29 | 24 | 72 |
| Fargesin | 29 | 25 | 48 |
| Phenprocoumon | 29 | 26 | -28 |
| Clopamide | 29 | 27 | 26 |
| Frovatriptan Succinate | 29 | 28 | -30 |
| (-)-Fenchone | 29 | 29 | 3 |

| | | | |
|---|---|---|---|
| Cefoxitin | 29 | 30 | -55 |
| Danazol | 29 | 31 | 54 |
| Bepridil hydrochloride | 29 | 32 | -252 |
| Boldine | 29 | 33 | -29 |
| Ruscogenin | 29 | 34 | -6 |
| Casticin | 29 | 35 | 12 |
| Abiraterone Acetate | 29 | 36 | -9 |
| Josamycin | 29 | 37 | 15 |
| Cariprazine HCl | 29 | 38 | -83 |
| Cefazolin | 29 | 39 | -28 |
| Baloxavir marboxil | 29 | 40 | 54 |
| (−)-Norepinephrine | 29 | 41 | 3 |
| Indinavir Sulfate | 29 | 42 | 61 |
| Irisflorentin | 29 | 43 | 97 |
| Vitexin | 29 | 44 | -21 |
| Gelsemine | 29 | 45 | -2 |
| Evodiamine | 29 | 46 | 70 |
| Dimenhydrinate | 29 | 47 | 23 |
| Xylometazoline | 29 | 48 | 65 |
| Benazepril | 29 | 49 | 24 |
| Glucosamine | 29 | 50 | 4 |
| Brimonidine | 29 | 51 | 4 |
| Sapropterin Dihydrochloride | 29 | 52 | 98 |
| Dihydrocapsaicin | 29 | 53 | 77 |
| Pneumocandin B0 | 29 | 54 | 98 |
| Triptonide | 29 | 55 | -46 |
| Tetrandrine | 29 | 56 | 88 |
| Succimer | 29 | 57 | -70 |
| Trifluoperazine | 29 | 58 | -316 |
| Bendamustine | 29 | 59 | 57 |
| 2-Aminoethanethiol | 29 | 60 | -49 |
| Tiapride Hydrochloride | 29 | 61 | 15 |
| Fedratinib (SAR302503, TG101348) | 29 | 62 | -81 |
| Magnolin | 29 | 63 | 54 |
| Dehydroandrographolide Succinate | 29 | 64 | -82 |
| Bacitracin Zinc | 29 | 65 | -90 |
| Ethisterone | 29 | 66 | 89 |
| Cefotaxime | 29 | 67 | -6 |
| Trazodone | 29 | 68 | 68 |
| Bepotastine | 29 | 69 | -24 |
| Chloramine-T | 29 | 70 | 26 |
| Protriptyline hydrochloride | 29 | 71 | 35 |
| Homoharringtonine | 29 | 72 | 58 |
| Morin | 29 | 73 | 21 |
| Homoorientin | 29 | 74 | -23 |
| Acetylcholine Chloride | 29 | 75 | 60 |
| Niclosamide | 29 | 76 | -70 |

| | | | |
|---|---|---|---|
| Anethole trithione | 29 | 77 | 20 |
| DL-Menthol | 29 | 78 | 85 |
| Bromhexine | 29 | 79 | 15 |
| Salicylamide | 29 | 80 | 10 |
| Octisalate | 30 | 1 | 28 |
| Mequitazine | 30 | 2 | 65 |
| Ibuprofen Piconol | 30 | 3 | 73 |
| Nifurtimox | 30 | 4 | 47 |
| Tinoridine Hydrochloride | 30 | 5 | 90 |
| Lasmiditan Succinate | 30 | 6 | 16 |
| Hydrocortisone Butyrate | 30 | 7 | 76 |
| Etofenamate | 30 | 8 | -47 |
| Hydroxyprogesterone | 30 | 9 | 65 |
| Temocapril | 30 | 10 | 73 |
| Tulobuterol Hydrochloride | 30 | 11 | 45 |
| Talniflumate | 30 | 12 | 64 |
| Apronal | 30 | 13 | 56 |
| Permethrin | 30 | 14 | 65 |
| Pinaverium Bromide | 30 | 15 | 76 |
| Doravirine | 30 | 16 | 85 |
| (+)-Equol | 30 | 17 | 59 |
| Omadacycline Tosylate | 30 | 18 | -2 |
| 3,4-Diaminopyridine | 30 | 19 | 23 |
| Roquinimex | 30 | 20 | 42 |
| Hexetidine | 30 | 21 | -195 |
| Halazone | 30 | 22 | 82 |
| Delavirdine | 30 | 23 | 95 |
| Amezinium (Methylsulfate) | 30 | 24 | 11 |
| Eicosapentaenoic Acid | 30 | 25 | 73 |
| Fosamprenavir Calcium Salt | 30 | 26 | 50 |
| Cyclothiazide | 30 | 27 | 40 |
| Letermovir (AIC246) | 30 | 28 | -27 |
| 5,5-Dimethyloxazolidine-2,4-dione | 30 | 29 | 58 |
| Tirapazamine | 30 | 30 | 26 |
| Zucapsaicin | 30 | 31 | 73 |
| Riboflavin Tetrabutyrate | 30 | 32 | 70 |
| Bicyclol | 30 | 33 | 42 |
| Tafamidis | 30 | 34 | 7 |
| Octodrine (2-Amino-6-Methylheptane | 30 | 35 | 46 |
| Ozenoxacin | 30 | 36 | 30 |
| Fursultiamine | 30 | 37 | 53 |
| Fadrozole | 30 | 38 | 24 |
| Taurocholic Acid Sodium Sal Hydrate | 30 | 39 | -38 |
| Isopropyl Myristate | 30 | 40 | 47 |
| Clemizole | 30 | 41 | 51 |
| Chlorphenesin | 30 | 42 | 52 |
| Docosahexaenoic Acid | 30 | 43 | 63 |

| | | | |
|---|---|---|---|
| Fosfluconazole | 30 | 44 | 49 |
| Fipexide Hydrochloride | 30 | 45 | 44 |
| Apraclonidine HCl | 30 | 46 | 25 |
| Avatrombopag | 30 | 47 | -27 |
| 4-Aminohippuric Acid | 30 | 48 | 29 |
| Embelin | 30 | 49 | 39 |
| Sevoflurane | 30 | 50 | 69 |
| Nimorazole | 30 | 51 | 41 |
| Chromium Picolinate | 30 | 52 | 30 |
| Clebopride (Malate) | 30 | 53 | 65 |
| Trandolapril | 30 | 54 | 53 |
| Fosphenytoin (Disodium) | 30 | 55 | 62 |
| Clobetasone Butyrate | 30 | 56 | 91 |
| Brequinar | 30 | 57 | 53 |
| Azilsartan | 30 | 58 | 49 |
| Pexidartinib | 30 | 59 | 58 |
| 2-Aminoethyl Diphenylborinate | 30 | 60 | 53 |
| Cevimeline | 30 | 61 | 10 |
| Triclocarban | 30 | 62 | 28 |
| Glycyrrhetinic Acid | 30 | 63 | 46 |
| Thonzylamine | 30 | 64 | 24 |
| Uridine Triacetate | 30 | 65 | 53 |
| Methylprednisolone Hemisuccinate | 30 | 66 | 44 |
| Belotecan Hydrochloride | 30 | 67 | 45 |
| Sodium 4-Aminosalicylate | 30 | 68 | 53 |
| Entrectinib | 30 | 69 | 90 |
| Ebselen | 30 | 70 | 57 |
| Udenafil | 30 | 71 | 87 |
| Carazolol | 30 | 72 | 36 |
| Flurbiprofen Axetil | 30 | 73 | 41 |
| Fluralaner | 30 | 74 | 65 |
| Fluticasone Furoate | 30 | 75 | 26 |
| Norethisterone Enanthate | 30 | 76 | 73 |
| Besifovir | 30 | 77 | 97 |
| 9-Aminocridine | 30 | 78 | 2 |
| Voxelotor | 30 | 79 | 54 |
| Myristic Acid | 30 | 80 | 0 |
| Perifosine (KRX-0401) | 31 | 1 | -40 |
| Galanthamine HBr | 31 | 2 | -18 |
| Cephalexin | 31 | 3 | -65 |
| Gadodiamide Hydrate | 31 | 4 | -6 |
| Procarbazine HCl | 31 | 5 | -13 |
| Gabapentin | 31 | 6 | 6 |
| Methacycline HCl | 31 | 7 | -6 |
| Zanamivir | 31 | 8 | -2 |
| (R)-baclofen | 31 | 9 | 10 |
| Thiamine HCl (Vitamin B1) | 31 | 10 | 9 |

| | | | |
|---|---|---|---|
| Palbociclib (PD-0332991) HCl | 31 | 11 | 66 |
| Granisetron HCl | 31 | 12 | -59 |
| Perindopril Erbumine | 31 | 13 | 21 |
| Nedaplatin | 31 | 14 | -60 |
| D-Cycloserine | 31 | 15 | 13 |
| Kanamycin sulfate | 31 | 16 | -43 |
| Lomefloxacin HCl | 31 | 17 | -8 |
| Plerixafor 8HCl (AMD3100 8HCl) | 31 | 18 | 100 |
| Caspofungin Acetate | 31 | 19 | 64 |
| Citicoline sodium | 31 | 20 | 5 |
| Pemetrexed | 31 | 21 | -7 |
| Biapenem | 31 | 22 | -27 |
| Cidofovir | 31 | 23 | 7 |
| Penicillamine | 31 | 24 | -23 |
| Sodium butyrate | 31 | 25 | -9 |
| Chondroitin sulfate | 31 | 26 | -39 |
| Amiloride HCl dihydrate | 31 | 27 | -50 |
| Geneticin (G418 Sulfate) | 31 | 28 | -49 |
| Creatinine | 31 | 29 | 17 |
| Cesium chloride | 31 | 30 | -7 |
| Gemcitabine HCl | 31 | 31 | -1 |
| Dorzolamide HCl | 31 | 32 | -14 |
| Ibuprofen Lysine | 31 | 33 | -10 |
| Etidronate | 31 | 34 | -8 |
| Taurine | 31 | 35 | -2 |
| Donepezil HCl | 31 | 36 | -18 |
| Oxacillin sodium monohydrate | 31 | 37 | -7 |
| Solifenacin succinate | 31 | 38 | 62 |
| Amikacin hydrate | 31 | 39 | -44 |
| Ceftazidime | 31 | 40 | -49 |
| Carboplatin | 31 | 41 | -1 |
| Mizoribine | 31 | 42 | -4 |
| Palbociclib (PD0332991) Isethionate | 31 | 43 | 72 |
| Tranexamic Acid | 31 | 44 | 6 |
| Clindamycin Phosphate | 31 | 45 | -10 |
| Neostigmine Bromide | 31 | 46 | -15 |
| Neomycin sulfate | 31 | 47 | -58 |
| Palonosetron HCl | 31 | 48 | -54 |
| Tripelennamine HCl | 31 | 49 | -32 |
| Penicillin V potassium salt | 31 | 50 | -18 |
| Leucovorin Calcium Pentahydrate | 31 | 51 | 3 |
| Polymyxin B sulphate | 31 | 52 | -25 |
| Alendronate sodium trihydrate | 31 | 53 | 8 |
| Levamisole hydrochloride | 31 | 54 | 3 |
| Lisinopril | 31 | 55 | 6 |
| Salbutamol Sulfate | 31 | 56 | -9 |
| Streptomycin sulfate | 31 | 57 | -37 |

| | | | |
|---|---|---|---|
| Miltefosine | 31 | 58 | -70 |
| Ibandronate sodium | 31 | 59 | 7 |
| Pirenzepine dihydrochloride | 31 | 60 | 7 |
| Pamidronate Disodium | 31 | 61 | -1 |
| Teicoplanin | 31 | 62 | 38 |
| Cytarabine | 31 | 63 | -3 |
| Ticlopidine HCl | 31 | 64 | -25 |
| Fosinopril Sodium | 31 | 65 | -35 |
| Tobramycin | 31 | 66 | -23 |
| Vancomycin HCl | 31 | 67 | 59 |
| Danofloxacin Mesylate | 31 | 68 | -39 |
| Abacavir sulfate | 31 | 69 | 11 |
| Imipenem | 31 | 70 | 10 |
| Gabapentin HCl | 31 | 71 | 20 |
| Varenicline Tartrate | 31 | 72 | -60 |
| L-Glutamine | 31 | 73 | 5 |
| ATP | 31 | 74 | -24 |
| Fudosteine | 31 | 75 | 7 |
| NAD+ | 31 | 76 | 19 |
| Hygromycin B | 31 | 77 | -41 |
| Amikacin disulfate | 31 | 78 | -23 |
| L-Arginine HCl (L-Arg) | 31 | 79 | 10 |
| Fondaparinux Sodium | 31 | 80 | -39 |
| Cangrelor Tetrasodium | 32 | 1 | -57 |
| DL-Glutamine | 32 | 2 | -12 |
| Netilmicin Sulfate | 32 | 3 | -103 |
| Amprolium HCl | 32 | 4 | -64 |
| Capreomycin Sulfate | 32 | 5 | -95 |
| Ceftazidime Pentahydrate | 32 | 6 | -57 |
| Pralidoxime chloride | 32 | 7 | -4 |
| D-(+)-Cellobiose | 32 | 8 | 6 |
| Manganese chloride | 32 | 9 | 5 |
| Methotrexate disodium | 32 | 10 | 13 |
| Acamprosate Calcium | 32 | 11 | 4 |
| L-SelenoMethionine | 32 | 12 | 12 |
| Bismuth Subcitrate Potassium | 32 | 13 | -56 |
| Bacitracin | 32 | 14 | -33 |
| Proflavine Hemisulfate | 32 | 15 | -55 |
| Guanethidine Sulfate | 32 | 16 | -76 |
| Eflornithine hydrochloride hydrate | 32 | 17 | 13 |
| L-Glutamic acid monosodium salt | 32 | 18 | 11 |
| Sodium carbonate | 32 | 19 | 26 |
| Homotaurine | 32 | 20 | 24 |
| isoleucine | 32 | 21 | -2 |
| Pemirolast potassium | 32 | 22 | 25 |
| Tetramisole HCl | 32 | 23 | -34 |
| Chloroquine Phosphate | 32 | 24 | -62 |

| | | | |
|---|---|---|---|
| Sodium ascorbate | 32 | 25 | 2 |
| Tolmetin Sodium | 32 | 26 | 11 |
| Captisol (SBE-β-CD) | 32 | 27 | 3 |
| Sodium ferulate | 32 | 28 | -4 |
| L(+)-Asparagine monohydrate | 32 | 29 | 26 |
| Calcium folinate | 32 | 30 | 26 |
| L-Leucine | 32 | 31 | 12 |
| Sodium Monofluorophosphate | 32 | 32 | 18 |
| Clodronate Disodium | 32 | 33 | 12 |
| Ceftriaxone Sodium Trihydrate | 32 | 34 | 5 |
| Apramycin Sulfate | 32 | 35 | -29 |
| Potassium Canrenoate | 32 | 36 | 12 |
| Glutathione | 32 | 37 | -28 |
| Danshensu | 32 | 38 | -32 |
| L-Threonine | 32 | 39 | 10 |
| D-Pantethine | 32 | 40 | -11 |
| L-Citrulline | 32 | 41 | 15 |
| Flavoxate HCl | 32 | 42 | 48 |
| Histamine Phosphate | 32 | 43 | -9 |
| Sodium Gluconate | 32 | 44 | -6 |
| Isepamicin Sulphate | 32 | 45 | -36 |
| Hydroxychloroquine Sulfate | 32 | 46 | -85 |
| L-Ornithine | 32 | 47 | -9 |
| Creatine phosphate disodium salt | 32 | 48 | 14 |
| Timonacic | 32 | 49 | -1 |
| Ozagrel sodium | 32 | 50 | 26 |
| L-Theanine | 32 | 51 | -7 |
| Dexamethasone Sodium Phosphate | 32 | 52 | 0 |
| Succinylcholine Chloride Dihydrate | 32 | 53 | -3 |
| Nefopam HCl | 32 | 54 | -51 |
| Micafungin Sodium | 32 | 55 | 31 |
| Dihydrostreptomycin sulfate | 32 | 56 | -71 |
| Cefradine | 32 | 57 | 7 |
| Glycylglycine | 32 | 58 | 7 |
| Nylidrin Hydrochloride | 32 | 59 | -39 |
| Sodium Demethylcantharidate | 32 | 60 | 0 |
| Colistin Sulfate | 32 | 61 | -12 |
| Terbutaline Sulfate | 32 | 62 | -16 |
| Paromomycin Sulfate | 32 | 63 | -60 |
| Amifostine | 32 | 64 | -52 |
| Sisomicin sulfate | 32 | 65 | -69 |
| Sildenafil | 32 | 66 | 51 |
| VitaMin U | 32 | 67 | 6 |
| Fosfomycin Disodium | 32 | 68 | -2 |
| Dimemorfan phosphate | 32 | 69 | -12 |
| L-Lysine hydrochloride | 32 | 70 | 14 |
| Gentamicin Sulfate | 32 | 71 | -34 |

| | | | |
|---|---|---|---|
| Eprazinone 2HCl | 32 | 72 | 34 |
| Ribostamycin Sulfate | 32 | 73 | -2 |
| Calcium Gluceptate | 32 | 74 | -19 |
| Eprodisate disodium | 32 | 75 | 8 |
| Choline bitartrate | 32 | 76 | -2 |
| Glycine | 32 | 77 | 8 |
| Pipemidic acid | 32 | 78 | -39 |
| Cephradine monohydrate | 32 | 79 | 8 |
| UTP, Trisodium Salt | 33 | 1 | 1 |
| DL-Arginine | 33 | 2 | -7 |
| L-Aspartic Acid | 33 | 3 | -9 |
| Penetrate Calcium Trisodium | 33 | 4 | 0 |
| Sodium Phytate Hydrate | 33 | 5 | 0 |
| Gadoversetamide | 33 | 6 | -14 |
| Gonadorelin Acetate | 33 | 7 | -25 |
| Angiotensin II Human Acetate | 33 | 8 | -15 |
| Sugammadex | 33 | 9 | -19 |
| Loxoprofen Sodium | 33 | 10 | -3 |
| L-Asparagine | 33 | 11 | 2 |
| L-Methionine | 33 | 12 | -1 |
| L-Hydroxyproline | 33 | 13 | -14 |
| Aprotinin | 33 | 14 | -24 |
| Bivalirudin Trifluoroacetate | 33 | 15 | -19 |
| Oxytocin | 33 | 16 | 99 |
| Carperitide Acetate | 33 | 17 | 82 |
| Gadoxetate Sodium | 33 | 18 | -20 |
| Sodium Succinate | 33 | 19 | 3 |
| Creatine | 33 | 20 | -1 |
| L-Arginine | 33 | 21 | 8 |
| Gastrodenol | 33 | 22 | -20 |
| Rilmenidine Phosphate | 33 | 23 | -14 |
| Eptifibatide Acetate | 33 | 24 | -16 |
| Salmon Calcitonin Acetate | 33 | 25 | 59 |
| Hyaluronic Acid | 33 | 26 | 42 |
| Latamoxef Sodium | 33 | 27 | -6 |
| Guanethidine Monosulfate | 33 | 28 | 11 |
| L-Cysteine | 33 | 29 | 7 |
| Bendazac L-Lysine | 33 | 30 | 13 |
| Pemetrexed Disodium Hydrate | 33 | 31 | 30 |
| Leuprorelin Acetate | 33 | 32 | -2 |
| Terlipressin Acetate | 33 | 33 | -8 |
| Foscarnet Sodium | 33 | 34 | -9 |
| Estramustine Phosphate Sodium | 33 | 35 | -21 |
| Spermine Tetrahydrochloride | 33 | 36 | -34 |
| L-Valine | 33 | 37 | -26 |
| Clodronate Disodium Tetrahydrate | 33 | 38 | 14 |
| Creatine Monohydrate | 33 | 39 | -4 |

| | | | |
|---|---|---|---|
| DL-Methionine | 33 | 40 | 5 |
| Lypressin Acetate | 33 | 41 | -19 |
| GHRP-2 | 33 | 42 | 2 |
| Citric Acid Trilithium Salt Tetrahydrate | 33 | 43 | 19 |
| β-Alanine | 33 | 44 | 25 |
| L-Proline | 33 | 45 | 2 |
| Lodoxamide Tromethamine | 33 | 46 | -16 |
| Malachite Green | 33 | 47 | -19 |
| L-Serine | 33 | 48 | -7 |
| Octreotide Acetate | 33 | 49 | 4 |
| Nafarelin Acetate | 33 | 50 | 79 |
| Somatostatin Acetate | 33 | 51 | -49 |
| Tobramycin Sulfate | 33 | 52 | -9 |
| L-Lysine | 33 | 53 | 1 |
| Edetate Trisodium | 33 | 54 | 4 |
| (R)-Serine | 33 | 55 | 12 |
| Adenosine Disodium Triphosphate | 33 | 56 | -2 |
| Alarelin Acetate | 33 | 57 | -15 |
| ε-Aminocaproic Acid | 33 | 58 | 40 |
| Goserelin Acetate | 33 | 59 | -30 |
| DL-Serine | 33 | 60 | 66 |
| L-Alanine | 33 | 61 | 27 |
| Edetate Calcium Disodium | 33 | 62 | -6 |
| (S)-Glutamic Acid | 33 | 63 | 10 |
| Mangafodipir Trisodium | 33 | 64 | 14 |
| Atosiban Acetate | 33 | 65 | -11 |
| Desmopressin Acetate | 33 | 66 | 52 |
| Chlorophyllin | 33 | 67 | 63 |
| Erlotinib HCl (OSI-744) | 34 | 1 | 68 |
| Irbesartan | 34 | 2 | -13 |
| Beta Carotene | 34 | 3 | 37 |
| Vinpocetine | 34 | 4 | 66 |
| Itraconazole | 34 | 5 | 37 |
| Sodium Aescinate | 34 | 6 | -59 |
| Ebastine | 34 | 7 | 9 |
| Alectinib hydrochloride | 34 | 8 | 77 |
| Zofenopril calcium | 34 | 9 | -10 |
| Vandetanib (ZD6474) | 34 | 10 | -76 |
| Norfloxacin | 34 | 11 | 33 |
| Flubendazole | 34 | 12 | 76 |
| Neratinib (HKI-272) | 34 | 13 | 84 |
| 7-Aminocephalosporanic acid | 34 | 14 | 13 |
| Sodium Houttuyfonate | 34 | 15 | -181 |
| Folic acid | 34 | 16 | 24 |
| Amantadine | 34 | 17 | 3 |
| Ofloxacin | 34 | 18 | 21 |
| Imiquimod | 34 | 19 | -43 |

| | | | |
|---|---|---|---|
| Risperidone | 34 | 20 | -51 |
| Oxibendazole | 34 | 21 | 33 |
| Sitafloxacin Hydrate | 34 | 22 | -26 |
| Trazodone HCl | 34 | 23 | -30 |
| Nuciferine | 34 | 24 | 39 |
| Netupitant | 34 | 25 | 70 |
| Balofloxacin Dihydrate | 34 | 26 | 27 |
| Droxidopa | 34 | 27 | 30 |
| Camptothecin | 34 | 28 | 31 |
| Sulfapyridine | 34 | 29 | -29 |
| Irsogladine | 34 | 30 | 26 |
| R788 (Fostamatinib) Disodium | 34 | 31 | 7 |
| Oseltamivir Phosphate | 34 | 32 | -14 |
| Vincamine | 34 | 33 | 2 |
| Aluminium hydroxide | 34 | 34 | 57 |
| Xanthopterin Hydrate | 34 | 35 | -49 |
| Dolutegravir Sodium | 34 | 36 | 9 |
| Ketoconazole | 34 | 37 | 83 |
| Methyldopa | 34 | 38 | 33 |
| Sarafloxacin HCl | 34 | 39 | 72 |
| Ketanserin | 34 | 40 | 20 |
| Mozavaptan | 34 | 41 | 81 |
| Retinyl (Vitamin A) Palmitate | 34 | 42 | 34 |
| Cetilistat | 34 | 43 | -51 |
| Garenoxacin | 34 | 44 | 18 |
| Mirogabalin | 34 | 45 | -20 |
| Cefoselis Sulfate | 34 | 46 | -16 |
| Torsemide | 34 | 47 | 5 |
| Meclizine 2HCl | 34 | 48 | 65 |
| (-)-Huperzine A (HupA) | 34 | 49 | 37 |
| Enrofloxacin | 34 | 50 | 11 |
| Clopidol | 34 | 51 | 1 |
| Mebhydrolin napadisylate | 34 | 52 | 69 |
| Cefadroxil hydrate | 34 | 53 | -15 |
| Brigatinib (AP26113) | 34 | 54 | -17 |
| Prazosin HCl | 34 | 55 | -87 |
| Eplerenone | 34 | 56 | -4 |
| Epalrestat | 34 | 57 | -32 |
| Diosmin | 34 | 58 | 74 |
| Ambroxol HCl | 34 | 59 | 32 |
| Chlortetracycline HCl | 34 | 60 | -34 |
| 6-Aminopenicillanic acid | 34 | 61 | -8 |
| Methenamine | 34 | 62 | 22 |
| Gefitinib hydrochloride | 34 | 63 | 9 |
| Marbofloxacin | 34 | 64 | 5 |
| Paliperidone | 34 | 65 | 5 |
| Lornoxicam | 34 | 66 | 14 |

| | | | |
|---|---|---|---|
| Hypoxanthine | 34 | 67 | -3 |
| Bilastine | 34 | 68 | 15 |
| Amoxapine | 34 | 69 | 19 |
| Sennoside A | 34 | 70 | 53 |
| Sulfaquinoxaline sodium | 34 | 71 | -72 |
| Oxaliplatin | 35 | 1 | 60 |
| Calcium gluconate | 35 | 2 | -5 |
| Sodium Hyaluronate | 35 | 3 | -13 |
| Heparin sodium | 35 | 4 | 13 |
| Deferiprone | 35 | 5 | 35 |
| Nisin | 35 | 6 | -86 |
| Risedronate Sodium | 35 | 7 | 10 |
| D-Phenylalanine | 35 | 8 | 70 |
| Disodium Cromoglycate | 35 | 9 | 19 |
| Minocycline HCl | 35 | 10 | -27 |
| Adenine sulfate | 35 | 11 | 38 |
| Alosetron Hydrochloride | 35 | 12 | -19 |
| Azasetron HCl | 35 | 13 | 35 |
| Piperaquine phosphate | 35 | 14 | 57 |
| Hydralazine HCl | 35 | 15 | 70 |
| Icatibant Acetate | 35 | 16 | 66 |
| Besifloxacin HCl | 35 | 17 | -25 |
| Plerixafor (AMD3100) | 35 | 18 | -91 |
