## Supplementary material for "A repurposed drug screen identifies compounds that inhibit the binding of the COVID-19 spike protein to ACE2": Suppl. Table 2

| Drug Name | Plate Number | Well Location | % Inhibition | EC50 (corrected) |
|---|---|---|---|---|
| Thiostrepton | 17 | 3 | 99 | 3.95E-06 |
| Oxytocin | 33 | 16 | 99 | 4.17E-06 |
| Nilotinib (AMN-107) | 1 | 52 | 99 | 4.20E-06 |
| (S)-10-Hydroxycamptothecin | 8 | 77 | 100 | 7.07E-06 |
| Hydroxy Camptothecine | 13 | 37 | 99 | 7.25E-06 |
| Nilotinib Hydrochloride | 22 | 33 | 99 | 8.38E-06 |
| Selamectin | 22 | 56 | 97 | 8.46E-06 |
| Picropodophyllin (PPP) | 27 | 67 | 98 | 9.97E-06 |
| Docetaxel | 1 | 5 | 98 | 1.03E-05 |
| Doramectin | 21 | 69 | 96 | 1.28E-05 |
| Anidulafungin | 16 | 29 | 99 | 1.31E-05 |
| Velpatasvir | 12 | 76 | 93 | 1.52E-05 |
| Zotarolimus(ABT-578) | 27 | 44 | 96 | 1.73E-05 |
| Estradiol Benzoate | 15 | 73 | 93 | 1.73E-05 |
| Aprepitant | 1 | 16 | 94 | 1.73E-05 |
| Irisflorentin | 29 | 43 | 97 | 1.75E-05 |
| Avermectin B1 | 20 | 50 | 98 | 1.80E-05 |
| Posaconazole | 1 | 40 | 97 | 1.87E-05 |
| Edoxaban | 27 | 15 | 95 | 1.87E-05 |
| Cefditoren Pivoxil | 4 | 70 | 96 | 1.88E-05 |
| Olmutinib | 28 | 13 | 98 | 1.91E-05 |
| Docetaxel Trihydrate | 27 | 78 | 95 | 2.05E-05 |
| Triamterene | 15 | 1 | 91 | 2.35E-05 |
| Biotin (Vitamin B7) | 11 | 5 | 91 | 2.74E-05 |
| Cabazitaxel | 10 | 58 | 86 | 2.88E-05 |
| Lenvatinib Mesylate | 22 | 6 | 94 | 3.49E-05 |
| Daclatasvir Dihydrochloride | 21 | 6 | 97 | 3.55E-05 |
| Arbidol HCl | 7 | 14 | 95 | 3.68E-05 |
| Dihydroergotamine Mesylate | 25 | 38 | 94 | 3.73E-05 |
| Cefotiam Hexetil Hydrochloride | 24 | 50 | 99 | 3.74E-05 |
| Daclatasvir | 3 | 3 | 99 | 4.50E-05 |
| Teniposide | 5 | 22 | 97 | 4.63E-05 |
| Cefetamet Pivoxil Hydrochloride | 20 | 44 | 94 | 5.07E-05 |
| Tipifarnib | 3 | 2 | 98 | 5.74E-05 |
| Entrectinib | 30 | 69 | 90 | 5.84E-05 |
| Crystal Violet | 5 | 70 | 98 | 6.00E-05 |
| Celecoxib | 1 | 80 | 93 | 6.15E-05 |
| Lomitapide Mesylate | 27 | 17 | 93 | 6.23E-05 |
| Avanafil | 14 | 73 | 99 | 6.59E-05 |
| Everolimus (RAD001) | 1 | 54 | 94 | 6.60E-05 |
| Cepharanthine | 16 | 54 | 94 | 6.77E-05 |
| Pneumocandin B0 | 29 | 54 | 98 | 7.28E-05 |
| Ombitasvir (ABT-267) | 23 | 16 | 92 | 7.28E-05 |
| Vilazodone HCl | 16 | 66 | 99 | 7.69E-05 |
| Temsirolimus (CCI-779, NSC 68386 | 1 | 23 | 96 | 8.00E-05 |
| Ciprofloxacin Hydrochloride Hydra | 22 | 53 | 93 | 8.02E-05 |

| | | | | |
|---|---|---|---|---|
| Fipronil | 23 | 53 | 91 | 9.31E-05 |
| Tadalafil | 3 | 64 | 96 | 1.00E-04 |
| Octenidine Dihydrochloride | 21 | 70 | 100 | 1.26E-04 |
| Tosufloxacin (P-Toluenesulfonic Ac | 22 | 10 | 97 | 1.64E-04 |
| Elbasvir | 25 | 40 | 100 | 2.12E-04 |
| Tinoridine Hydrochloride | 30 | 5 | 90 | 2.57E-04 |
| Mometasone furoate | 6 | 4 | 98 | 2.76E-04 |
| Amodiaquine hydrochloride | 24 | 22 | 90 | 5.00E-04 |
| Apremilast (CC-10004) | 27 | 80 | 96 | 1.39E-02 |
| Nelfinavir Mesylate | 16 | 68 | 94 | 7.40E-02 |
