## Supplementary material for "A repurposed drug screen identifies compounds that inhibit the binding of the COVID-19 spike protein to ACE2": Suppl. Table 3

| Drug | Bond Type |
|---|---|
| **Thiostrepton** | H-bond |
| | π -alkyl bond |
| | π -Sulfur bond |
| **Oxytocin** | H-bond |
| | π-alkyl bond |
| | Alkyl bond |
| | Salt bridge |
| **Nilotinib (AMN-107)** | H-Bond |
| | π-π T-shaped |
| | π-sigma |
| | Halogen bond |
| | π-anion |
| **S-(10)-Hydroxycamptothecin** | H-bond |
| | π-stacked |
| | Alkyl bond |
| | π-cation |
| **Hydroxycamptothecin** | H-bond |
| | π-stacked |
| | Alkyl bond |
| | π-cation bond |
| **Nilotinib HCl** | H- bond |
| | π-π T-shaped |
| | π-sigma bond |
| | Halogen bond |
| | π-anion bond |
| **Selamectin** | H-bond |
| | Alkyl bond |
| | π-alkyl bond |
| **Picropodophyllin** | H-bond |
| | π-stacked |
| | π-cation bond |
| **Docetaxel** | H-bond |
| | π -alkyl bond |
| **Doramectin** | H-bond |
| | π-sigma bond |
| | Alkyl bond |
| **Anidulafungin** | H-bond |
| | π-π T-shaped |
| | π -alkyl bond |

| | |
|---|---|
| | π –anion bond |
| **Estradiol Benzoate** | H-bond |
| | π-alkyl bond |
| | Alkyl bond |
|  | π-π stacked |

| 6VSB Glide score & Interactions | ACE2 Glide score & Interactions |
|---|---|
| Arg403, Glu406, **Lys417**, Asp420, **Tyr453**, Asn460, **Gln493**, **Tyr505** | **Asp30**, **Asp38**, **Arg393** |
| Tyr495 | **His34**, Ala387 |
| ---- | **His34** |
| Arg403, Glu406, **Lys417**, Gly447, **Gln493 Gln498**, Asn501 | **His34**, **Glu35**, **Gln42**, Glu75 |
| Tyr449 | ---- |
| ---- | **Lys31** |
| ---- | **Asp38** |
| Asn501 | **Arg393** |
| **Tyr449**, Tyr495, **Gly496** | Tyr349 |
| **Gln493** | ---- |
| **Phe490**, Leu492 | Ala348, Asp382, His401 |
| ---- | Asp350, Asp382 |
| Arg403, Gln414, **Lys417** | **Asp30**, **Glu37** |
| ---- | Gln388 |
| ---- | Lys26, Pro389 |
| ---- | **Arg393** |
| Arg403, Gln414, **Lys417** | **Asp30**, **Glu37** |
| ---- | Gln388 |
| ---- | Lys26, Pro389 |
| ---- | **Arg393** |
| Asn501 | **Arg393** |
| **Tyr449**, Tyr495, **Gly496** | Tyr349 |
| **Gln493** | ---- |
| **Phe490**, Leu492 | Ala348, Asp382, His401 |
| ---- | Asp350, Asp382 |
| Arg403, Gln409, Gln414, **Lys417**, **Tyr453** | **Asp30**, **Lys353**, **Gly354**, **Arg393** |
| **Lys417** | ---- |
| ---- | **His34** |
| Lys458, **Gln474** | Asp350 |
| Ala475 | Phe40 |
| Lys458 | |
| **Tyr449**, **Gln493**, **Asn501** | Asn394, **Arg393**, Lys562 |
| **Tyr449** | Leu73, **Arg393**, His401 |
| Ile468, Thr470, Leu492, **Ser494** | **Gly354** |
| **Phe490** | ---- |
| Leu452, **Leu455** | Leu45, Asn49 |
| Asp405, Arg408, | **Glu35**, Glu75 |
| **Tyr505** | |
| **Tyr449** | **Lys353**, Ala386 |

| | |
|---|---|
| **----** | **Glu35** |
| **Phe490** | Gln102 |
| Cys488 | **Arg393** |
| **Phe486** | Leu73, Ala99 |
| **----** | Phe40 |
